# Discovery of a taxusin-mediated route to baccatin III enables its complete biosynthesis in engineered microbes

**DOI:** 10.64898/2026.07.31.742161

**Authors:** Chengshuai Yang, Zhenhua Li, Linjie Yu, Yan Wang, Liangzhen Zheng, Xing Yan, Wenping Wei, Bingtong Feng, Tong Zhang, Jianxu Li, Pingping Wang, Zhihua Zhou

## Abstract

Taxol (paclitaxel) is a frontline anticancer drug widely applied for the treatment of breast, ovarian and lung cancers. Currently, its supply mainly relies on the semi-synthesis using baccatin III from *Taxus* plants. Heterologous biosynthesis of baccatin III in microorganisms offers a promising solution to alleviate global Taxol supply shortage, but remains challenging due to pathway complexity. Here, we report a novel taxusin-mediated biosynthetic pathway for baccatin III production *via* the identification of C13 deacetylase, elucidation of the exact sequence underlying C1 hydroxylation, and stepwise enzymatic functional validation. Through protein engineering of the promiscuous C1 and C5 hydroxylases, coupled with the distribution of pathway modules in *Saccharomyces cerevisiae* and *Escherichia coli*, we achieved the *de novo* biosynthesis of baccatin III. Collectively, our findings remodel the current biosynthetic framework governing the formation of Taxol precursors and highlight the great potential of microbial cell factories for the production of complex plant-derived therapeutic compounds.

**Highlights:** • Discovery of C13 deacetylase reveals a novel biosynthetic route to baccatin III *via* taxusin

• Stepwise verification of the complete biosynthetic route to baccatin III through taxusin and baccatin VI

• Single-site mutation reversed the product selectivity of T1OH and converted T5OH into a specific taxoid C5 hydroxylase

• Complete biosynthesis of baccatin III in engineered microbes

## INTRODUCTION

Taxol (paclitaxel), originally isolated from *Taxus* species, is a potent and widely used anticancer agent^1–5^. Taxol only accumulates at an extremely low level in yew tissues (∼0.01% of dry weight), and no scalable total chemical synthesis method is available for industrial production^6–24^. For these reasons, currently the global supply of Taxol hinges on the semisynthetic production^5,25^. In the semisynthetic workflow, baccatin III (**19**) or 10-deacetyl baccatin III are first extracted from the needles of *Taxus* trees, followed by industrial chemical derivatization to yield Taxol^12,26^. Though the semi-synthesis strategy avoids felling entire yew trees, it is still restricted by limited wild and cultivated *Taxus* biomass resources^5,27^. Heterologous biosynthesis of Taxol or its precursor baccatin III (**19**) in microbial or plant chassis represents a promising strategy to overcome Taxol supply constraints and facilitates the development of new-generation taxane anticancer drugs, as this sustainable, eco-friendly approach eliminates reliance on natural yew raw materials^1,27–33^.

Over the past three years, substantial breakthroughs have been made in deciphering the Taxol biosynthetic pathway. Key advances include the identification of several previously uncharacterized essential enzymes namely oxetane synthase^27,31,34^, C1^29,35,36^/C9^27,29,31^/C2^37^ hydroxylases, C7 acetylase^27^ and C7/C9 deacetylases^29^; the full elucidation of the biosynthetic route constructing Taxol’s tetracyclic core skeleton^27^ (Figures 1A and S14); the successful *de novo* biosynthesis of baccatin III (**19**) in *Nicotiana benthamiana*^29,31,35^; and the clarification of the formation mechanism for governing Taxol’s vital oxetane ester moiety *via* isotope labeling experiments assays^27,34^ and computational modeling^27,31,34^. Collectively, these discoveries lay a solid foundation for complete elucidation of the biosynthetic pathway of Taxol in yew and its *de novo* biosynthesis in heterologous microbial chassis, and thus expedite the development of robust microbial cell factory dedicated to Taxol heterologous synthesis.

**Figure 1.**
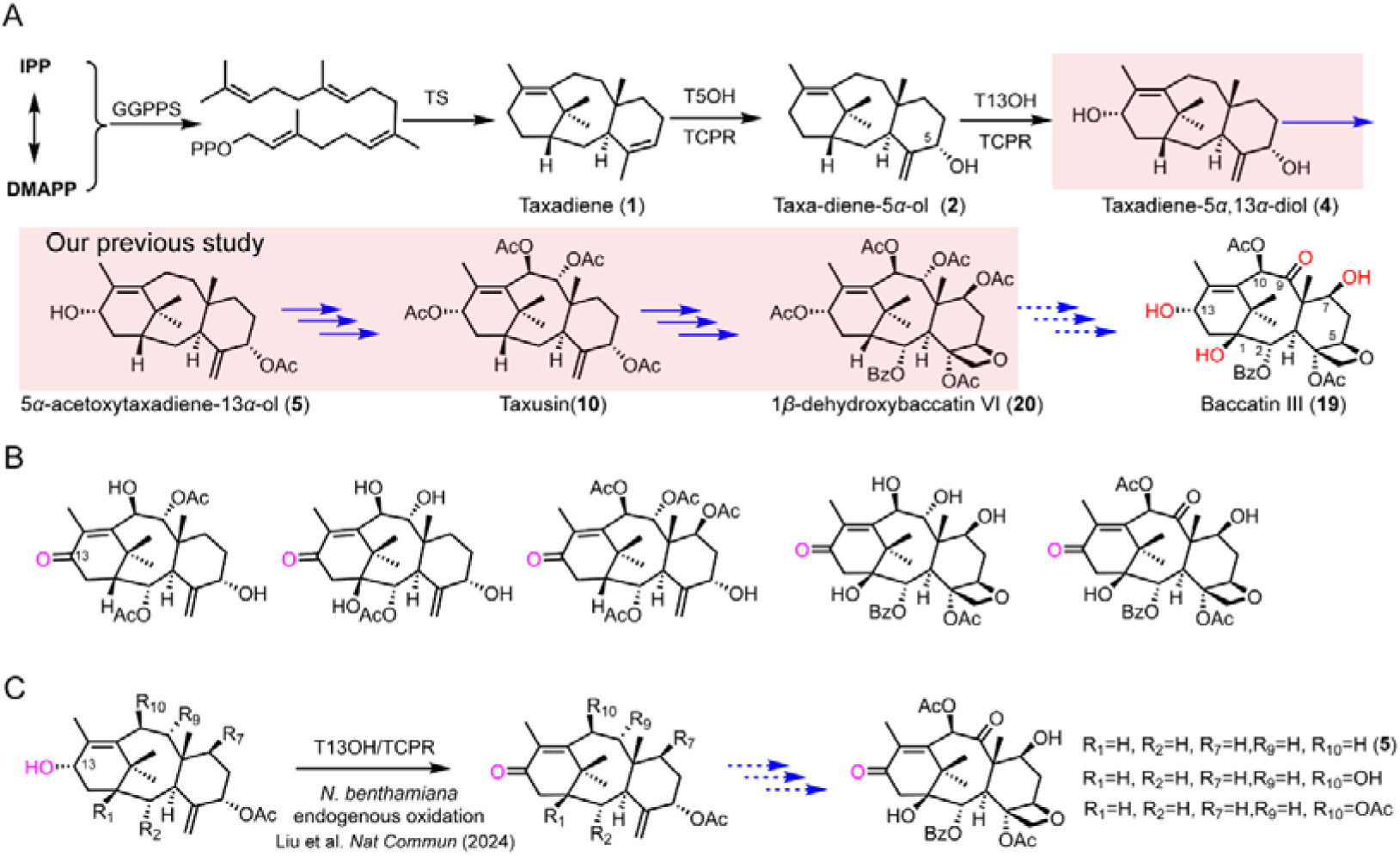
The proposed biosynthetic pathway of baccatin III and taxoids with ketone at C13. (A) The proposed baccatin III biosynthetic pathway *via* taxusin (**10**). The biosynthetic pathway of 1β-dehydroxybaccatin VI (**20**) in pink box was confirmed in our previous study^27^. (B) Representative taxoids with ketone at C13 isolated from *Taxus* plant^39^. (C) T13OH and endogenous enzymes from *Nicotiana benthamiana* catalyze oxidation of the C13-hydroxyl group of taxoids to yield the corresponding C13-ketone taxoids^30^, which may account for the formation of C13-ketone-containing taxoids in planta.

Nevertheless, previous reconstructions of baccatin III biosynthetic pathway in *N. benthamiana* and functional characterizations of its key enzymes has predominantly conducted *via* multiplexed gene co-expression in *N. benthamiana*^29,31,35^, which could identify products of a group of enzymes rather than that of the single-gene expression coupled with exogenous substrate feeding. This approach obscures stepwise validation of enzymatic reactions in the biosynthetic pathway. Meanwhile, it could not exclude the contribution of endogenous enzymes and metabolisms from *N. benthamiana*^30,31,37^. Ambiguity regarding the native or preferred reaction sequence within this intricate pathway may lead to misidentification of the authentic rate-limiting steps. Stepwise validation of enzymatic reactions in the biosynthetic pathway is crucial for the confirmation of biosynthetic pathway, discovery of limiting steps and heterologous biosynthesis.

In 2024, we elucidated a stepwise taxusin-mediated biosynthetic pathway converting taxadiene-5α-ol (**2**) to the highly oxygenated tetracyclic metabolite 1β-dehydroxybaccatin VI (**20**)^27^ (Figures 1A and S14). In 2025, Conor James McClune et al. proposed three potential biosynthetic routes to baccatin III (**19**). Based on metabolites detected in a multiplexed gene co-expression system in *N. benthamiana*, one putative route features 5α,10β-diacetoxytaxadien-13α-ol, 13-deacetyltaxusin and 13-deacetyl-1β-dehydroxybaccatin VI (**21**) as key intermediates^29^. For the other two proposed branching biosynthetic routes toward baccatin III (**19**), only putative intermediates —taxusin (**10**), baccatin VI (**16**), and 9-dihydro-13-acetylbaccatin III (**17**)— were indicated, without detailed stepwise enzymatic reactions being suggested^29^.

Accumulating evidence from multiple studies has demonstrated that the baccatin III (**19**) biosynthetic pathway is a complex network rather than a simple linear cascade^27,30,34,38^. The discovery of C7 and C9 deacetylases indicates that the acetyl moieties at C7 and C9 positions likely act as protecting groups to stabilize highly oxygenated intermediates during baccatin III (**19**) biosynthesis. A variety of C13-ketone taxoids have been isolated from the *Taxus* plant^39^ (Figure 1B), and several such C13-ketone taxoids have also been detected in the *N. benthamiana* engineered with the upstream paclitaxel biosynthetic genes^30^ (Figure 1C). Furthermore, we also observed the taxoid 13α-hydroxylase (T13OH) catalyzed 5α-acetoxytaxadiene-13α-ol (**5**) to yield 5α-acetoxytaxadiene-13*-*one (**6**) in yeast cell (Figures 1C and S1). In addition, taxusin-13-one was unexpectedly obtained during the isolation and purification of taxusin (**10**) for NMR analysis in our previous study^27^. Since the free C13 hydroxy group is susceptible to oxidation to yield a ketone, the installation and subsequent removal of a protecting group at the C13 position would facilitate baccatin III (**19**) biosynthesis. Accordingly, identification of the hitherto uncharacterized C13 deacetylase is essential to validate this hypothesis.

The biosynthetic pathway of baccatin III (**19**) is intricate, consisting of at least 17 enzymatic reactions mediated by 18 distinct enzymes. These enzymes include taxadiene synthase, which constructs the tricyclic skeleton taxadiene (**1**)^40–43^; sixteen tailoring enzymes (seven hydroxylases targeting C5^44,45^, C13^46^, C10^40,47^, C9^27,29,31^, C7^48^, C2^49^ and C1^29,36^, five acyltransferases modifying the hydroxyl groups at C2^49^, C5^50,51^, C7^27,29^, C9^52^ and C10^53^, two taxane deacetylases acting on C7-OAc and C9-OAc^29^, as well as an oxetane synthase^27,31,34^ and an oxidase for C9-OH^29,35^) ; and the taxane oxidation facilitator FoTO1^29^.

Several enzymes within this pathway generate promiscuous products, such as the C5 hydroxylase^29,44,45,54–58^, C1 hydroxylase^29,36^ and oxetane synthase^27,31,34^. Meanwhile, other enzymes exhibit broad substrate tolerance, including the C10^27,40,47^, C7^48^ and C1^29,36^ hydroxylases, and alongside the two taxane deacetylases^29^. The discovery of FoTO1 markedly boosts the catalytic activity of CYP725A4 (T5OH)^29^. Even so, the desired product taxadiene-5α-ol (**2**) only accounts for only ∼40% of the total catalytic output, as an equivalent quantity of the by-product 4α,20-epoxy-taxadiene-5α-ol (**3**) and miscellaneous side products are simultaneously produced^29^. Such catalytic promiscuity diverts metabolic flux away from baccatin III (**19**) biosynthesis toward dead-end metabolites, impeding the development of high-efficiency microbial factories. Enhancing the substrate and product selectivity of these promiscuous enzymes is therefore essential to achieve *de novo* heterologous biosynthesis of baccatin III (**19**) in engineered cell factories. To date, only early-stage di-oxygenated taxoids intermediates have been successfully *de novo* biosynthesized in microbial hosts^55,59,60^. We have previously built a yeast cell factory to produce 1β-dehydroxybaccatin VI (**20**) by feeding taxadiene-5α-ol (**2**). The heterologous production of baccatin III (**19**) remains unattainable in microbial chassis, in which a bottleneck likely stems from the extreme complexity of the paclitaxel biosynthetic cascade and poor functional compatibility of plant-derived enzymes within microbial heterologous systems.

In this study, we identified a C13 deacetylase from yew, enabling us to establish a stepwise validated biosynthetic pathway to baccatin III (**19**) *via* taxusin (**20**) and baccatin VI (**16**). We successfully realized *de novo* biosynthesis of intermediate baccatin VI (**16**) in yeast cell factory, followed by biotransformation of baccatin VI (**16**) into baccatin III (**19**) in *Escherichia coli* cell. Collectively, these findings lay a solid foundation for engineering microbial cell factories to produce baccatin III (**19**) and Taxol.

## RESULTS

### Integrating mass spectrometry imaging and single-cell transcriptome enables the discovery of C13 deacetylase

Our previous study has elucidated the biosynthetic pathway of 1β-dehydroxybaccatin VI (**20**) with the tetracyclic core skeleton of Taxol^27^, in which taxusin (**10**) is a key intermediate. Building on this foundation, we further explored the tailoring modifications that convert 1β-dehydroxybaccatin VI (**20**) to baccatin III (**19**) (Figure 1A) in this study. This process was proposed to be involved in deacetylation reactions at C7, C9, and C13, oxidation at C9 and hydroxylation at C1. Given that taxusin (**10**) and 1β-dehydroxybaccatin VI (**20**) possess an additional acetyl group at C13 position relative to baccatin III (**19**), we hypothesized that a C13 deacetylase participates in the biosynthesis of baccatin III (**19**). We therefore attempted to mine the C13 deacetylase involved in this biosynthetic route. We first visualized the spatial distribution of baccatin III (**19**) in stem of *Taxus× media via* mass spectrometry imaging. The result revealed specific accumulation of baccatin III (**19**) in stem epidermis (Figure 2A and S2A-S2C), suggesting that the enzymes responsible for the late-stage biosynthesis of baccatin III (**19**) may be specifically expressed in epidermis. Subsequent single-cell transcriptomic analysis confirmed that the known key taxiod C9 oxidase (T9ox)^29^ exhibited epidermis-specific expression in stem (Figures 2B and S2D). Accordingly, we hypothesized that the C13 deacetylase involved in this pathway might be also specifically expressed in stem epidermis cells.

**Figure 2.**
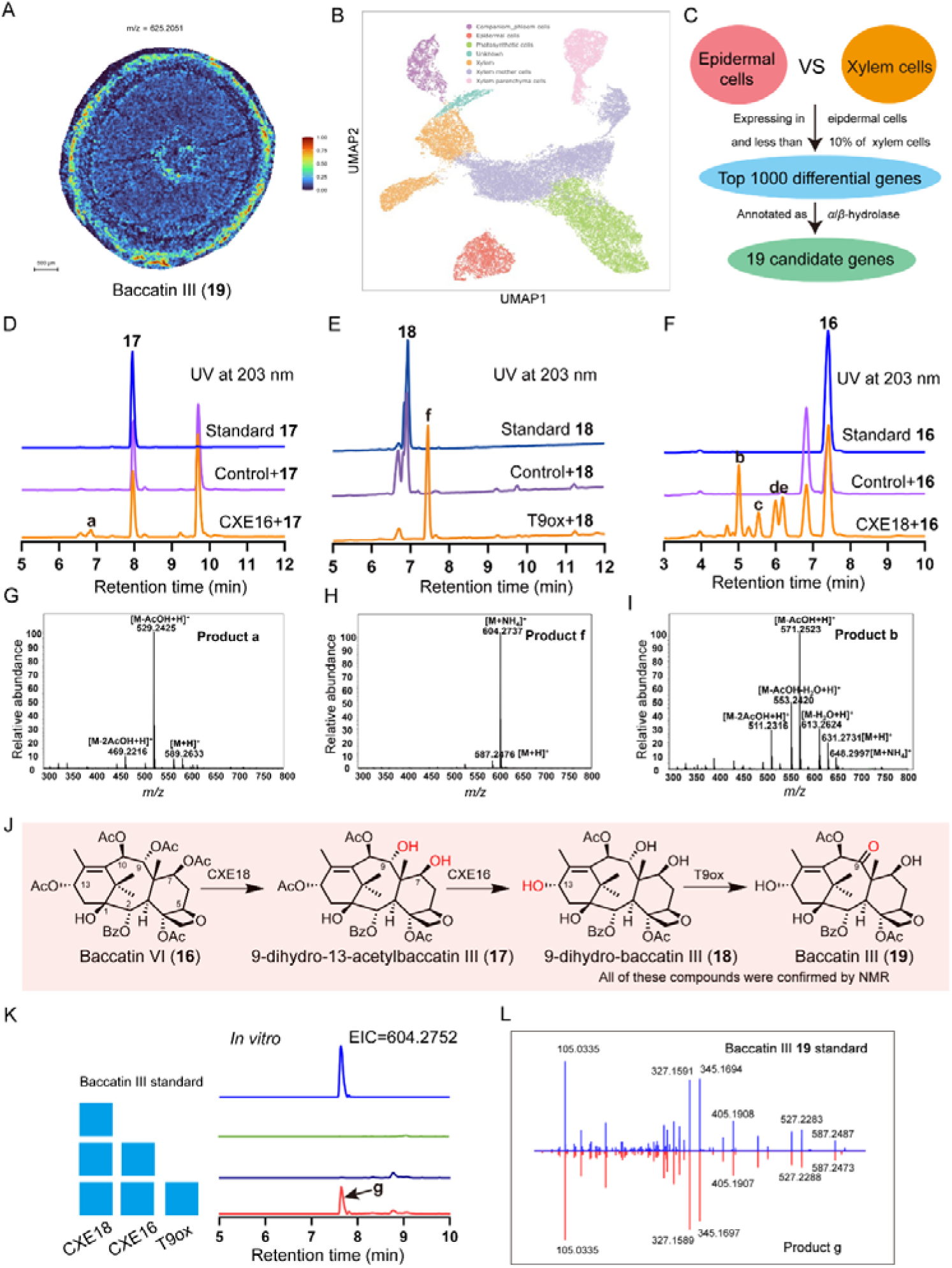
Biosynthesis of baccatin III (19) from baccatin VI (16). (A) The distribution of baccatin III in stem, detected by MALDI analysis. The color scale ranges from 0 to 1. (B) Visualization of cell clusters from stem using the UMAP method. Different cell clusters were marked with different colors. (C) Strategy used to mine candidate genes for C13 deacetylase. (D) HPLC analysis of *in vitro* reaction using crude enzyme of *E. coli* expressing CXE16 with 9-dihydro-13-acetylbaccatin III (**17**) as substrate. Crude enzyme of *E. coli* expressing empty vector was used as control. (E) HPLC analysis of *in vitro* reaction using crude enzyme of *E. coli* expressing T9ox with 9-dihydro-baccatin III (**18)** as substrate. (F) HPLC analysis of *in vitro* reaction using crude enzyme of *E. coli* expressing CXE18 with baccatin VI (**16**) as substrate. Crude enzyme of *E. coli* expressing empty vector was used as control. (G-I) Mass spectra (ESI) of products a, b and f. (J) The biosynthetic pathway of baccatin III (**19**) from baccatin VI (**16**). (K) The combinatorial catalysis of CXE16, CXE18 and T9ox to produce baccatin III (**19**) from baccatin VI (**16**). Crude enzymes of *E. coli* respectively expressing CXE16, CXE18, and T9ox were used to perform *in vitro* combinatorial catalysis with baccatin VI (**16**) as substrate. (I) MS/MS fragmentation patterns of product g from *in vitro* combinatorial catalysis of CXE16, CXE19 and T9ox using baccatin VI (**16**) as substrate compared to that of baccatin III (**19**) standard.

We then screened the potential C13 deacetylase from 19 candidate genes belonging to α/β-hydrolases family in the top 1000 differential genes (Figure 2C). These candidate genes were functionally characterized through *in vitro* enzymatic assays using crude proteins extracted from *E. coli* expressing individual candidates, with the corresponding substrate supplied for each reaction. High performance liquid chromatography (HPLC) analysis demonstrated that the candidate protein CXE16 could catalyze the conversion of 9-dihydro-13-acetylbaccatin III (**17)** into a more polar product (Figure 2D). Combined spectra analyses, including high-resolution electrospray ionization mass spectrometry (HR-ESIMS) and nuclear magnetic resonance (NMR) spectroscopy such as ^1^H, ^13^C (Figures 2G and 2J), we identified this product as 9-dihydro-baccatin III (**18**), which lacks the C13 acetyl group. These results confirm that CXE16 functions as a C13-specific deacetylase. Further enzymatic assays showed that CXE16 also could remove the C13 acetyl group of 1β-dehydroxybaccatin VI (**20)** to produce 13-deacetyl-1β-dehydroxybaccatin VI (**21)** while exhibiting no catalytic activity towards baccatin VI (**16)** (Figures S2E-H). The identification of CXE16 indicates that reversible the C13 acetylation and deacetylation may serve as a protective mechanism for unstable C13-deacetylated intermediates to avoid being oxidated to corresponding C13-ketone taxoids, exemplified by 13-deacetyltaxusin.

To verify whether 9-dihydro-baccatin III (**18**) serves as an intermediate en route to baccatin III (**19**), we tested the activity of T9ox toward 9-dihydro-baccatin III (**18**). T9ox was previously characterized to oxidize 9-dihydro-7-acetylbaccatin III to generate 7-acetylbaccatin III^29^. HPLC analysis revealed that T9ox could also oxidize 9-dihydro-baccatin III (**18**) and produce a less polar metabolite. Based on HR-ESIMS and NMR spectroscopic analyses, this product was identified as baccatin III (**19**) (Figures 2E, 2H and 2J), confirming that 9-dihydro-13-acetylbaccatin III (**17**) and 9-dihydro-baccatin III (**18**) are intermediates for baccatin III (**19**).

We subsequently sought candidate C7 and C9 deacetylases to catalyze baccatin VI (**16**) to 9-dihydro-13-acetylbaccatin III (**17**). *In vitro* enzymatic assays were carried out with baccatin VI (**16**) as the substrate against 19 candidate α/β-hydrolases. These candidates were either highly expressed in high-Taxol-producing cell lines, or exhibited upregulated expression levels relative to low-taxol-yielding cell lines (Figure S3A)^61^. HPLC analysis showed that CXE18 catalyzed the transformation of baccatin VI (**16**) into multiple products (Figure 2F). The predominant product was assigned as 9-dihydro-13-acetylbaccatin III (**17**) by HR-ESIMS and NMR spectroscopic analyses (Figures 2F, 2I, and 2J). Collectively, these data confirm that CXE18 is a bifunctional enzyme capable of simultaneously removing the acetyl groups at C7 and C9 positions of baccatin VI (**16**). Sequence alignment indicated low amino acid sequence identity between CXE18 and the previously characterized taxane 9α-O-deacetylase (74.3%) and taxane 7β-O-deacetylase (56%)^29^. Additional substrate profiling confirmed that CXE18 exhibits a broad substrate spectrum. In addition to baccatin VI (**16**), CXE18 catalyzes C7 and C9 deacetylation of other taxoids, including 1β-dehydroxybaccatin VI (**20**), 13-deacetyl-1β-dehydroxybaccatin VI (**21**), and 13-deacetylbaccatin VI (**22**) (Figure S4). This finding suggests that the late-stage biosynthetic steps toward baccatin III follow a non-linear route.

Accordingly, we elucidated the biosynthetic route toward baccatin III (**19**) starting from baccatin VI (**16**) involving the deacetylations at C7, C9, and C13 positions together with the C9 oxidation. Further combinatorial catalysis was performed using crude preparations of *E. coli*-expressed CXE16, CXE18 and T9ox with baccatin VI (**16**) as the substrate. The *in vitro* reaction successfully produced baccatin III (**19**), thereby validating this biosynthetic pathway from baccatin VI (**16**) to baccatin III (**19**) (Figures 2K and 2L).

### Identification of a novel taxusin-derived biosynthetic route to baccatin III *via* **distinct C1 hydroxylation and stepwise enzymatic validation**

We have elucidated the biosynthetic pathway of 1β-dehydroxybaccatin VI (**20**) previouly^27^ and newly characterized CXE16 could catalyze the C13 deacetylation of 1β-dehydroxybaccatin VI (**20**) to generate 13-deacetyl-1β-dehydroxybaccatin VI (**21**), . Accordingly, we next sought to identify the key enzyme responsible for C1 hydroxylation of either 13-deacetyl-1β-dehydroxybaccatin VI (**21**) or 1β-dehydroxybaccatin VI (**20**) for the formation of 13-deacetyl-baccatin VI (**22**) or baccatin VI (**16**) to fill the last gap of baccatin III (**19**) biosynthetic pathway. Therefore, 1β-dehydroxybaccatin VI (**20**) and its C13 deacetylated derivate 13-deacetyl-1β-dehydroxybaccatin VI (**21**) were selected as substrates to identify the C1 hydroxylase. We screened 51 candidate P450 genes of CYP725A subfamily via *in vivo* feeding assays, together with 18 candidate 2-oxoglutarate-dependent dioxygenase (2OGD) genes that were highly expressed in high-taxol-yielding cell lines, or exhibited upregulated tendency relative to low-taxol-yielding cell lines (Figure S3B) by *in vitro* enzymatic reactions. Reactions of these candidate genes were analyzed by HPLC, however, we failed to identify the target C1 hydroxylase.

Considering numerous natural tricyclic taxoids possess a C1 hydroxyl group and lack an oxetane moiety^39^, we screened for the C1 hydroxylase using highly oxygenated tricyclic taxoids 7β-acetoxytaxusin-2α-ol (**12**) as a substrate. HPLC analysis revealed that the 2OGD TOGD5 efficiently catalyzes the conversion of taxoids 7β-acetoxytaxusin-2α-ol (**12**) into two products. The major and minor products were identified as 5,10,13-acetyl-10-debenzoylbrevifoliol-2α-ol (**14**, a) and 7β-acetoxytaxusin-1β,2α-diol (**13**, b), respectively, based on HR-ESIMS and NMR spectroscopic analyses (Figures 3A, 3D, and 3I), confirming TOGD5 as a C1 hydroxylase hereafter renamed T1OH. T1OH shares 98.4% amino acid identity with T1βH-184^29^, which was previously supposed to catalyze the C1 hydroxylation of 13-deacetyl-1β-dehydroxybaccatin VI (**21**) in a putative biosynthetic route toward baccatin III (**19**). However, T1OH displayed no activity toward three compounds bearing a C2 benzoyl group, 13-deacetyl-1β-dehydroxybaccatin VI (**21**), 2α-benzoyloxy-7β-acetoxytaxusin (**23**) and 1β-dehydroxybaccatin VI (**20**), as confirmed by HPLC and LC-MS analyses (Figure S5), suggesting that the bulky benzoyl substituent suppresses C1 hydroxylation. Taken together, these results demonstrate that T1OH-mediated C1 hydroxylation takes place before C2 benzoylation and oxetane ring formation, which is fundamentally distinct from the previously proposed biosynthetic scheme^29^.

**Figure 3.**
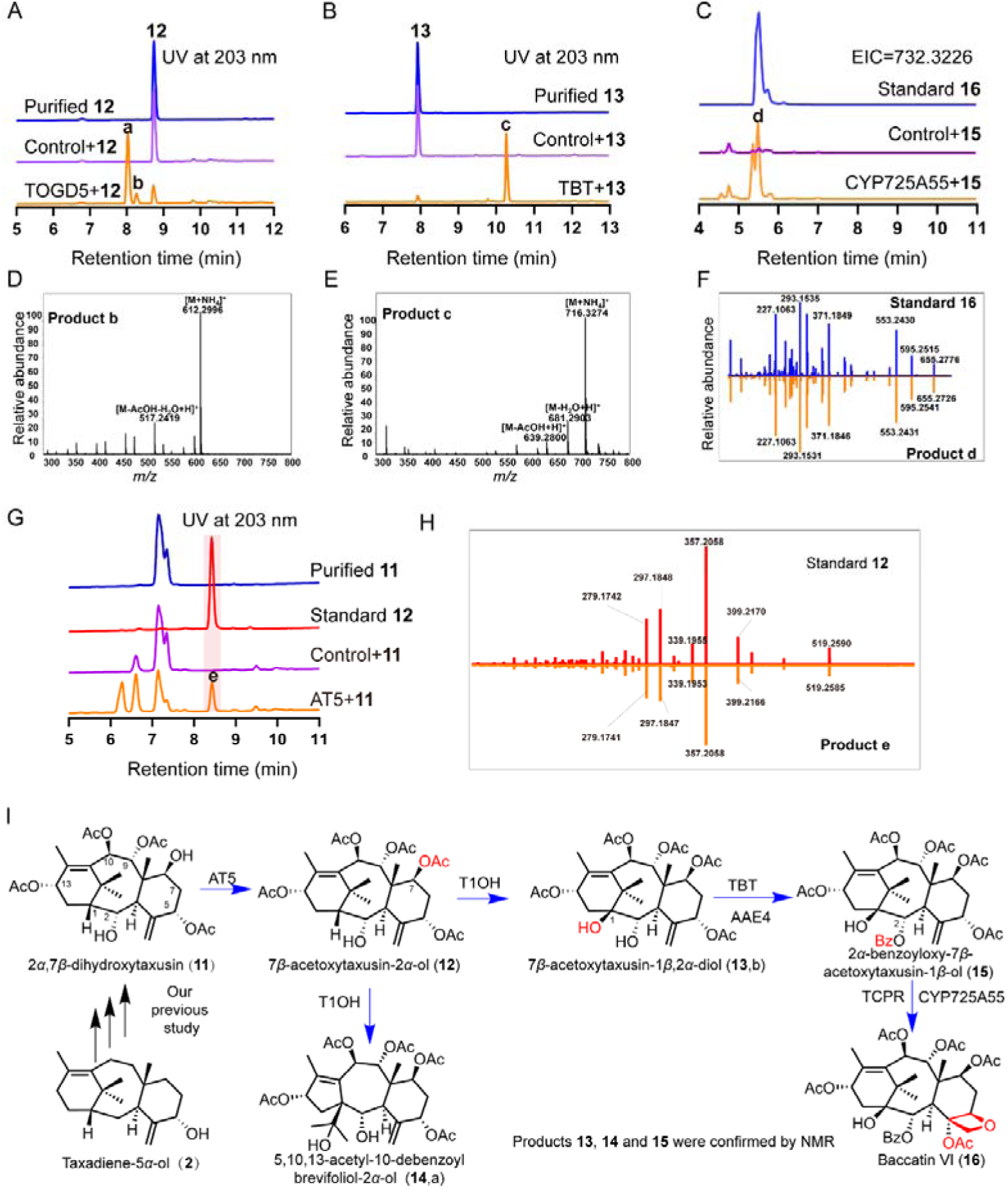
The complete biosynthetic reaction order of baccatin VI (16). (A) HPLC analysis of *in vitro* reaction using crude enzyme of *E. coli* expressing TOGD5 with 7β-acetoxytaxusin-2α-ol (**12**) as substrate. Crude enzyme of *E. coli* expressing empty vector was used as control. (B) HPLC analysis of *in vitro* reaction using crude enzyme of *S. cerevisiae* strain expressing TBT and AAE4 with 7β-acetoxytaxusin-2α,1β-diol (**13**) as substrate. Crude enzyme of *S. cerevisiae* strain YBD80 was used as control. (C) LC-MS analysis of *in vivo* feeding experiment using *S. cerevisiae* strain YCYP725A55 expressing CYP725A55 with 2α-benzoyloxy-7β-acetoxytaxusin-1β-ol (**15**) as substrate. Extracted ion chromatogram (EIC) =732.3226. *S. cerevisiae* strain YTCPR was used as the control. (D-E) Mass spectra (ESI) of products b and c. (F) MS/MS fragmentation patterns of product d from *in vivo* feeding experiment using 2α-benzoyloxy-7β-acetoxytaxusin-1β-ol (**15**) as substrate compared to that of baccatin VI (**16**) standard. (G) HPLC analysis of *in vitro* reaction using crude enzyme of *E. coli* expressing AT5 with 2α,7β-dihydroxytaxusin (**11**) as substrate. Crude enzyme of *E. coli* expressing empty vector was used as control. (H) MS/MS fragmentation patterns of product e from *in vitro* reaction using crude enzyme of *E. coli* expressing AT5 with 2α,7β-dihydroxytaxusin (**11**) as substrate compared to that of 7β-acetoxytaxusin-2α-ol (**12**) standard. (I) The biosynthetic pathway of baccatin VI (**16**) from taxadiene-5α-ol (**2**).

We further verified the stepwise biosynthetic pathway from 7β-acetoxytaxusin-2α,1β-diol (**12**) to baccatin VI (**16**) through the stepwise conversion. *In vitro* enzymatic reactions showed that TBT^52^ benzoylated 7β-acetoxytaxusin-1β,2α-diol (**13**) to generate 2α-benzoyloxy-7β-acetoxytaxusin-1β-ol (**15**), which was structurally assigned by HR-ESIMS and NMR spectroscopy analyses (Figures 3B, 3E, and 3I). *In vivo* feeding experiments further revealed that the oxetane synthase CYP725A55 oxygenated 2α-benzoyloxy-7β-acetoxytaxusin-1β-ol (**15**) to form baccatin VI (**16**), consistent with LC-MS and MS/MS fragmentation patterns analyses (Figures 3C, 3F, and 3I). As we reported earlier, 2α,7β-dihydroxytaxusin (**11**) undergoes acetylation catalyzed by the characterized C7-OH acetyltransferase AT5 to yield 7β-acetoxytaxusin-2α-ol (**12**) (Figures 3G-3I). Besides, A stepwise biosynthetic pathway to generate 2α,7β-dihydroxytaxusin (**11**) from taxadiene-5α-ol (**2**) has been characterized in our previous study ^27^ (Figure S14).

Taken together, we elucidated a novel biosynthetic pathway for baccatin III (**19**) *via* taxusin (**10**) and baccatin VI (**16**), and provided an experimentally verified stepwise route toward baccatin III (**19**).

### Single site mutation reversed T1OH product selectivity

Wild-type T1OH primarily produces the dead-end metabolite 5,10,13-acetyl-10-debenzoyl brevifoliol-2α-ol (**14**) while the target product 7β-acetoxytaxusin-1β,2α-diol (**13**) accounts for only 13.6 % of the total products (Figures 4C and 4D). This low target product yield represents a well-recognized bottleneck limiting the efficient biosynthesis of baccatin III. To enhance the proportion of the target compound catalyzed by T1OH, we conducted heterologous expression and protein purification of T1OH, and followed by crystallization screening to resolve its three-dimensional crystal structure. Despite extensive screening attempts, no ligand-bound complex crystals were successfully obtained. Therefore, the machine learning-based protein ligand-binding site prediction model PointSite^62^ was employed to identify potential residues based on the apo crystal structure of T1OH (Figure 4A). Nineteen candidate sites with high pocket confidence scores were screened out and individually mutated to alanine for functional verification. Two mutants, T1OH_M1 and T1OH_M2, exhibited improved product selectivity relative to wild-type T1OH (Figure S6). Among them, T1OH_M1 displaying the optimal selectivity by increasing the target product proportion from 13.6% to 43.8% (Figures 4C and S6). Subsequent saturation mutants of these two key active sites further reversed the product distribution ratio between the dead-end metabolite and the target product (Figures 4C, S7, and S8). Specifically, mutant T1OH_M3 produced 7β-acetoxytaxusin-1β,2α-diol (**13**) as the predominant product, with a significantly elevated proportion of 65.2% (Figures 4C and 4D). Nevertheless, further combinational mutagenesis based on T1OH_M3 failed to achieve additional improvements in product selectivity (Figure S9B).

**Figure 4.**
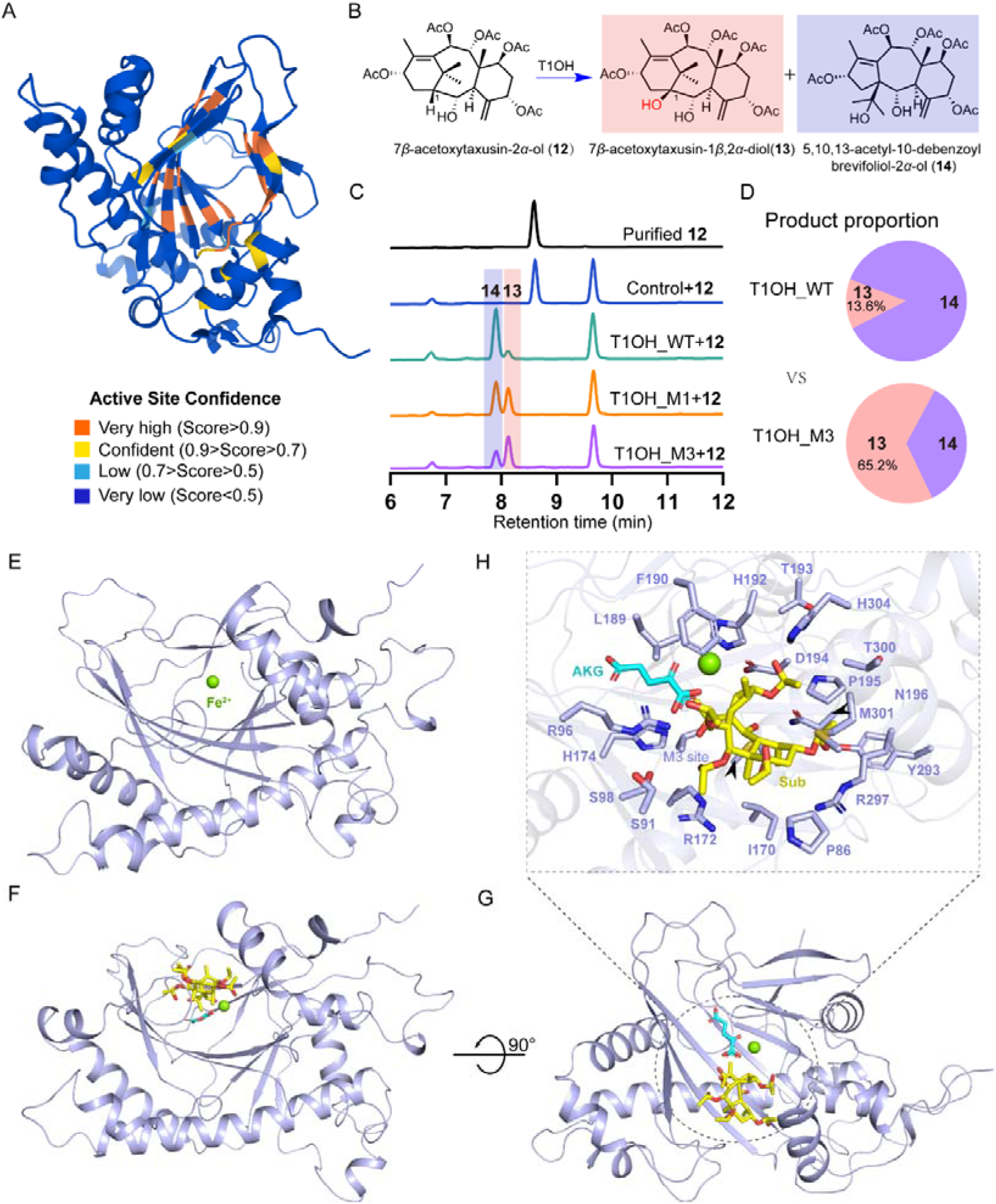
Enzymatic engineering and crystal structure of T1OH. (A) Crystal structure of T1OH and proposed active sites of T1OH predicted by PointSite^62^. (B) Hydroxylation reaction catalyzed T1OH. (C) HPLC analysis of *in vitro* reaction using crude enzyme of *E. coli* expressing T1OH and its mutants using 7β-acetoxytaxusin-2α-ol (**12**) as substrate. Crude enzyme of *E. coli* expressing empty vector was used as control. (D) The product proportion of wild type and mutant of T1OH. (E) Crystal structure of T1OH. (F-G) Top view and side view of docking analysis of the α-ketoglutarate and substrate 7β-acetoxytaxusin-2α-ol (**12**) with T1OH. The substrate and α-ketoglutarate molecules were marked with yellow and cyan, respectively. (H) Zoomed-in view of the intermolecular interactions at T1OH. The interacting residues are identified in stick representations.

To elucidate the structural mechanism underlying the altered product selectivity of T1OH, molecular docking simulations of the catalytic substrate and α-ketoglutarate (AKG) were performed using the solved apo structure as the receptor model (Figures 4E-4G). Docking analysis demonstrated that the target product exhibits favorable spatial compatibility with the catalytic pocket of T1OH and can be stably accommodated within the active-site cavity. Subsequent structural characterization of the amino acid residues lining the catalytic pocket further revealed that this binding cavity is predominantly composed of hydrophobic residues (Figure 4H). The T1OH_M3 mutation replaces the original residue with bulky side chain by a smaller amino acid, thereby effectively expanding the internal volume of the active-site cavity (Figure S9C). This structural alteration disrupts the canonical binding mode of AKG and remodels its coordination geometry within the catalytic pocket. Consequently, the trajectory of the catalytic oxidation reaction is subtly modulated, and the enzymatic catalytic partitioning is redirected. This remodeling enables the mutant enzyme to preferentially generate the target product 7β-acetoxytaxusin-1β,2α-diol (**13**) rather than the dead-end metabolite 5,10,13-acetyl-10-debenzoyl brevifoliol-2α-ol (**14**).

In summary, we successfully obtained a high-performance T1OH mutant (T1OH_M3) that produces the target compound as the dominant product, and systematically elucidated the structural mechanism responsible for the enhanced product selectivity of this mutant.

### Engineering promiscuous T5OH as a specificity taxoid C5 hydroxylase

The catalytic promiscuity of T5OH, which leads to the accumulation of numerous undesired byproducts (Figure 5A), has long been a critical bottleneck restricting Taxol biosynthesis as reported in previous studies^44,45,55–57,59^. The nuclear transport factor 2 (NTF2)-like protein FoTO1 was previously identified to facilitate the conversion of taxadiene (**1**) to taxadiene-5α-ol (**2**)^29^. However, this reaction is accompanied by the equivalent production of the byproduct 4α,20-epoxy-taxadiene-5α-ol (**3**) alongside the target product taxadiene-5α-ol (**2)** (Figure 5B). Therefore, improving the product selectivity of T5OH to produce taxadiene-5α-ol(**2**) is essential to redirect more diterpene precursor flux into the Taxol biosynthesis. To identify the key residues governing the product selectivity of T5OH, we performed virtual mutagenesis screening. A total of 26 T5OH mutants were designed *via* structural prediction using FoldX, followed by molecular docking simulations with Autodock Vina docking (ten independent replicates) to screen for mutants with elevated near-attack conformation ratios. All these designed mutants were functionally tested *via* individual expression in a yeast cell factory that produces the precursor taxadiene (**1**). GC-FID analysis demonstrated that T5OH_M1 mutant exhibited excellent product selectivity, predominantly generating the target product taxadiene-5α-ol (**2**) (Figure S10).

**Figure 5.**
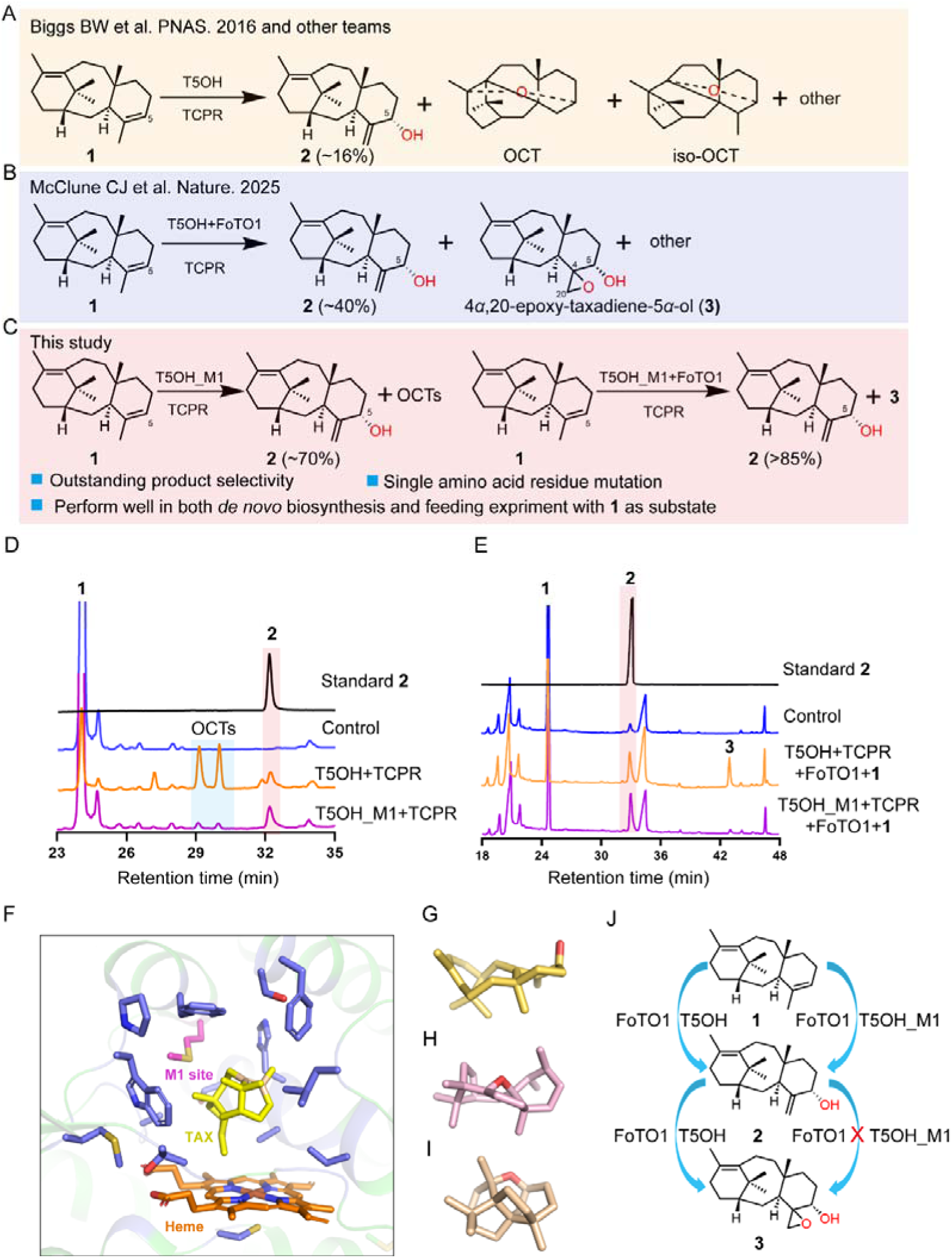
Enzymatic engineering of T5OH. (A) The promiscuous products of T5OH. (B) Co-expressing T5OH with FoTO1 changes the product distribution of T5OH. (C) The product proportion of T5OH mutant without or with FoTO1. (D) GC-FID analysis of products of *S. cerevisiae* strains YT5OH and YT5OH_M1 respectively expressing T5OH and its mutant. Yeast cell factory YSP producing taxadiene (**1**) was used as control. (E) GC-FID analysis of products of *S. cerevisiae* strains YFT5OH and YFT5OH_M1 respectively expressing FoTO1 with T5OH or its mutant fed with taxadiene (**1**). *S. cerevisiae* strain YTCPR was used as the control. (F) Docking analysis of the substrate taxadiene (**1**) and heme with previously reported crystal structure of T5OH (PDB code: 8X1W)^63^. (G-I) Computational structures of products taxadiene-5α-ol (**2**), OCT and iso-OCT. (J) Proposed formation process of 4α,20-epoxy-taxadien-5α-ol (**3**) catalyzed by FoTO1 and T5OH.

Subsequent overexpression of T5OH_M1 *via* multi-copy genomic integration in the taxadiene-producing yeast confirmed that T5OH_M1 prioritizes the synthesis of taxadiene-5α-ol (**2**) in *de novo* biosynthesis systems, with the proportion of the desired product elevated to nearly 70% (Figures 5C and 5D). Whole-cell catalytic assays *via* co-expression of wild-type T5OH or T5OH_M1 mutant with FoTO1 by feeding substrate taxadiene (**1**) further verified the improved selectivity of T5OH_M1. The T5OH_M1/FoTO1 system efficiently converted substrate (**1**) into the target product (**2**) (>85%), whereas wild type of T5OH/FoTO1 system yielded equal amounts of the target product taxadiene-5α-ol (**2**) and 4α,20-epoxy-taxadiene-5α-ol (**3**) (Figure 5E). Consistently, Taxadiene-5α-ol (**2**) became the dominant product in yeast strains co-expressing T5OH_M1 and FoTO1(Figure S11C). Further assays revealed that the co-expression of wild-type T5OH and FoTO1 can catalyze the secondary oxidation of taxadiene-5α-ol (**2**) to generate 4α,20-epoxy-taxadiene-5α-ol (**3**) (Figures S11A and S11D). This indicates that the enhanced product selectivity of the T5OH_M1 /FoTO1 system is attributed to the reduced catalytic activity of T5OH_M1 towards the target product taxadiene-5α-ol (**2**), which prevents secondary byproduct formation (Figures 5J and S11B). Collectively, these results confirm that the single-site mutation of T5OH_M1 confers superior product selectivity, effectively eliminating byproduct generation regardless of the presence or absence of FoTO1.

Combined with structural comparisons of substrate and product conformational features (Figures 5F-5I), we propose a structural mechanism underlying the altered catalytic selectivity of T5OH_M1. In wild-type T5OH, the short aliphatic side chain at the key mutation site imposes negligible steric hindrance, providing minimal spatial restriction on catalytic intermediates and nascent products. Following catalysis, reaction intermediates undergo unimpeded intramolecular folding and cyclization-induced conformational collapse, preferentially forming the compact, sterically bulky polycyclic architecture corresponding to the off-pathway by-product. In contrast, the aromatic benzyl side chain introduced by the T5OH _M1 mutation substantially increases local steric volume. This substitution thus remodels the steric microenvironment of the active-site cleft and introduces significant steric clashes, which block the intramolecular compaction and structural rearrangement required for by-product formation. Meanwhile, the target product adopts an extended, planar backbone conformation with a small spatial footprint, which is structurally compatible with the constricted active-site cavity of the T5OH_M1 mutant. In summary, T5OH_M1 mutation reshapes the steric topography of the enzyme active site, differentially regulating the conformational feasibility of two competing catalytic pathways. This structural remodeling redirects metabolic flux toward target product synthesis and effectively suppressing aberrant by-product generation.

### Biosynthesis of baccatin III in microorganisms

CXE16, CXE18 and T9ox have been verified to catalyze the deacetylation and oxidation reactions during the biosynthesis of baccatin III (**19**) from baccatin VI (**16**). Nevertheless, CXE16 and CXE18 exhibit substrate promiscuity and can remove acetyl moieties at C9 or C10 positions of several putative key intermediates e.g. taxusin (**10**) and 2α-hydroxytaxusin (Figures S2, S4, and S12). These unintended catalytic activity towards such putative intermediates may drive the generation of dead-end products, thereby impeding the *de novo* baccatin III biosynthesis. Furthermore, all three enzymes display pH-dependent activity (Figure S13). Specifically, CXE16 and CXE18 prefer alkaline conditions (Figure S13). However, the intracellular pH of *S. cerevisiae* becomes acidic (< 6) in the late stage of fermentation^64^, which disfavors the deacetylation reactions catalyzed by CXE16 and CXE18. To address this bottleneck, we split the biosynthetic route toward baccatin III (**19**) into two modules: *de novo* biosynthesis of baccatin VI (**16**) in yeast (owing to the requirement of multiple P450 enzymes for the baccatin VI (**16**) biosynthesis), followed by bioconversion of baccatin VI (**16**) to baccatin III (**19**) in *E. coli*. The intracellular environment of *E. coli* maintains an alkaline pH ^65^, which accommodates the activity of these three enzymes (Figure 6A).

**Figure 6.**
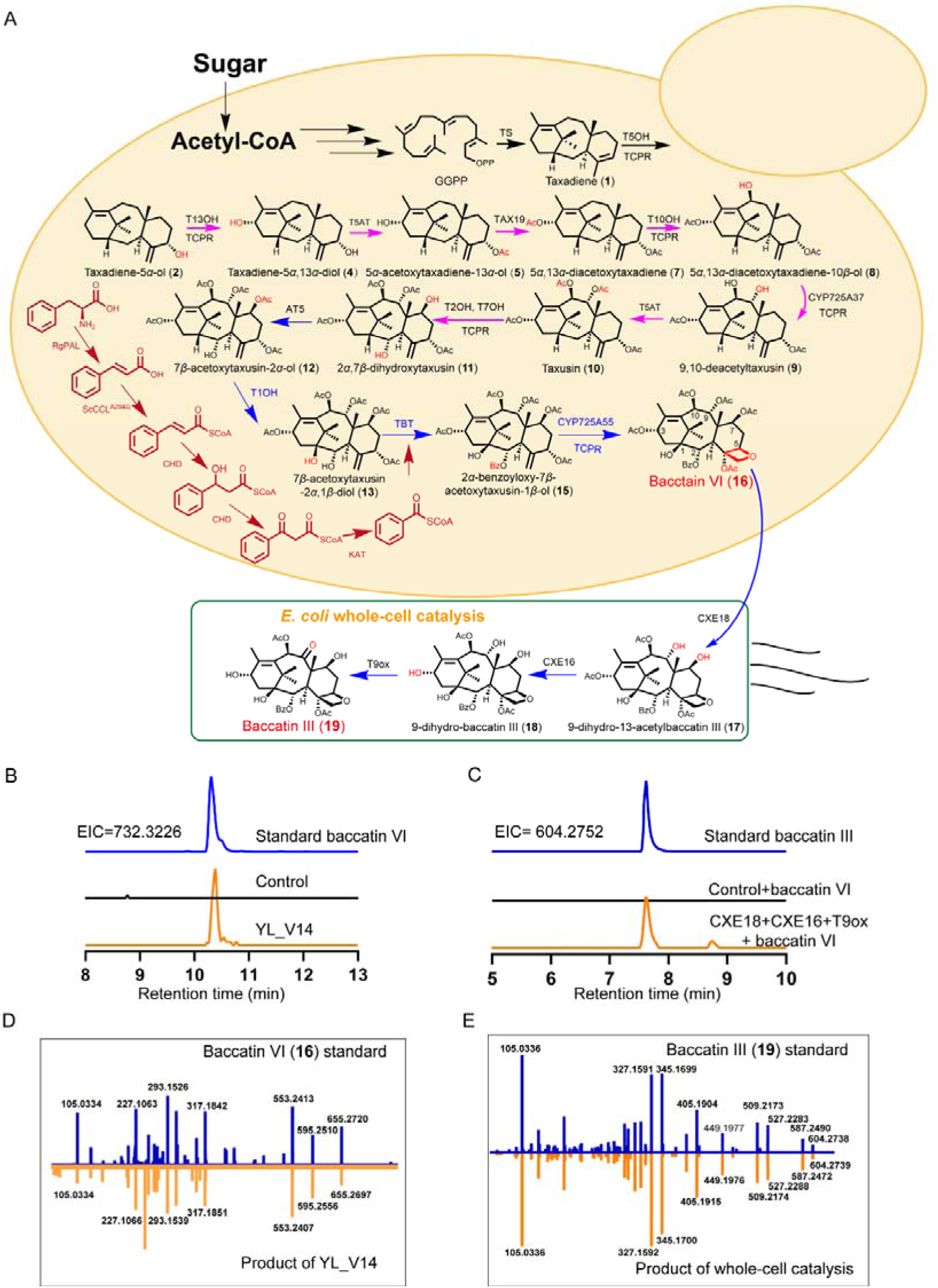
Biosynthesis baccatin III in microorganisms. (A) Biosynthesis of baccatin III (**16**) *via de novo* biosynthesis of baccatin VI in *S. cerevisiae* and bioconversion of baccatin III (**19**) from baccatin VI (**16**) using *E. coli* simultaneously expressing CXE16, CXE18 and T9ox. Reactions marked with magenta arrows were characterized and validated in our previous study. Reactions marked with blue arrows were characterized in this study. (B) LC-MS analysis of product of *S. cerevisiae* cell factory YL_V14 for biosynthesis baccatin VI (**16**). Extracted ion chromatogram (EIC) = 732.3226. *S. cerevisiae* cell factory CHD18 for biosynthesis 1β-dehydroxybaccatin VI (**20**) was used as control. (C) LC-MS analysis of product of whole-cell catalysis of *E. coli* strain EBAIII simultaneously expressing CXE16, CXE18 and T9ox using baccatin VI (**16**) as substrate. Extracted ion chromatogram (EIC) = 604.2752. *E. coli* strain EBK expressing empty vectors was used as control. (D) MS/MS fragmentation patterns of product from product of *S. cerevisiae* cell factory YL_V14 compared to that of baccatin VI standard (**16**). (E) MS/MS fragmentation patterns of product from whole cell catalysis of *E. coli* EBAIII using baccatin VI (**16**) as substrate compared to that of baccatin III standard (**19**).

To achieve *de novo* biosynthesis of baccatin VI (**16**) in yeast, we selected our previous reported strain YDBVI as the starting chassis. This strain harbors 12 *Taxus*-derived genes that convert mono-oxygenated taxoid taxadiene-5α-ol (**2**) to construct the tetracyclic core skeleton of Taxol. A series of metabolic engineering strategies were implemented for the biosynthesis of baccatin VI (**16**): (1) introduction of an upstream module enabling biosynthesis of precursor taxadiene-5α-ol (**2**) from glucose; (2) integrating an artificial benzoyl-CoA biosynthesis pathway assembled with enzymes (including RgPAL, ScCCL^A294G^, PhCHD and PhKAT) from *Rhodotorula glutinis*, *Streptomyces coelicolor* and *Petunia hybrida*^66,67^ to boost benzoyl-CoA supply; (3) overexpression of downstream module consisting of a T1OH_M3 mutant, DBAT and additional copy of T2OH (Figures 6A and S14A). LC-MS analysis confirmed the presence of baccatin VI (**16**) in the engineered yeast strain YL_V14, which validated the reconstructed biosynthetic route toward Baccatin VI (**16**) (Figures 6A, 6B and 6D). Nevertheless, substantial accumulation of by-product 1β-dehydroxybaccatin VI (**20**) was also observed (Figure S14). Further overexpression of T5OH_M1 and T1OH_M3 mutants, together with reinforcement of the biosynthesis of taxadiene (**1**), significantly increased the titers of baccatin VI (**16**) and by-product 1β-dehydroxybaccatin VI (**20**) (Fiugre 14D). These results suggest that the relatively weaker hydroxylation activity of the T1OH_M3 mutant relative to the benzoylation activity of TBT in the downstream pathway constitutes the major bottleneck in cell factory YL_V14 for baccatin VI (**16**) biosynthesis. When *E. coli* whole-cells co-expressing CXE16, CXE18 and T9ox were fed with baccatin VI (**16**), baccatin III (**19**) was detected by LC-MS analysis (Figures 6A, 6C, and 6E). This result demonstrates the feasibility of *de novo* production of baccatin III (**19**) from simple carbon feedstocks using engineered microbes.

## DISCUSSION

In this study, we uncovered a previously uncharacterized taxusin-derived biosynthetic route to baccatin III (**19**) through the identification of a novel C13 deacetylase, resolution of the precise sequence of C1 hydroxylation, and stepwise enzymatic validation. We engineered promiscuous enzymes to boost catalytic specificity and limit flux diversion to off-pathway metabolites. By partitioning the pathway between *S. cerevisiae* and *E. coli*, we achieved full biosynthesis of baccatin III in engineered microbial cell factories.

Taxusin (**10**) has been isolated from *Taxus* plants and identified as an abundant metabolite in the heartwood of yew previously^39^. However, it has been regarded as a dead-end metabolite rather than a biosynthetic intermediate en route to baccatin III^48,68^. In 2024, we elucidated a stepwise biosynthetic pathway converting taxadiene-5α-ol (**2**) to the highly oxygenated tetracyclic metabolite 1β-dehydroxybaccatin VI (**20**) (Figure 1A), establishing taxusin as a key intermediate in Taxol biosynthesis for the first time^27^. Taxusin was largely overlooked in other investigations of Taxol biosynthesis, primarily because C13 acetylation was presumed to shunt the taxane scaffold away from the productive biosynthetic pathway^48,50,68^. The discovery of the C13 deacetylase CXE16 revised this prevailing view: by removing the C13 acetyl moiety from late-stage tetracyclic taxanes, CXE16 reconnects taxusin-derived intermediates to baccatin III formation (Figure 6). Furthermore, the identification of C13, C7 and C9 deacetylases reveals that acetylation–deacetylation-mediated protection and deprotection exert crucial functions during baccatin III biosynthesis, serving as a reversible protecting-group strategy to protect unstable intermediates and inhibit the formation of byproducts like other reported biosynthesis of complex plant natural products^41–43^.

Resolving the exact order of C1 hydroxylation and C2 benzoylation is essential to decipher the taxusin-derived biosynthetic route toward baccatin III. In an earlier proposed biosynthetic route to baccatin III (**19**) built upon metabolite profiling from a multiplexed gene co-expression system in *N. benthamiana*, 13-deacetyl-1β-dehydroxybaccatin VI (**21**) was hypothesized to be the key intermediate and substrate of the C1 hydroxylase^29^. However, we have verified that the C1 hydroxylases characterized herein (T1OH) and the previously reported T1βH-184 fail to catalyze C1 hydroxylation of taxanes bearing a C2 benzoyl group, including 13-deacetyl-1β-dehydroxybaccatin VI (**21**) and 1β-dehydroxybaccatin VI (**20**) (Figure S5). This inactivity arises from steric interference imposed by the bulky C2 benzoyl substituent. Therefore, C1 hydroxylation occurs prior to C2 benzoylation and oxetane ring formation, rendering 1β-dehydroxybaccatin VI (**20**) a byproduct instead of a biosynthetic intermediate toward baccatin III (**19**) (Figures 6 and S14).

Plants have evolved promiscuous enzymes to generate vast chemical diversity. Meanwhile, they develop sophisticated spatial organization including tissue-, cell-and organelle-level compartmentalization to construct specialized microenvironments supporting distinct enzymatic reactions. These two evolutionary adaptations create substantial obstacles for microbial heterologous biosynthesis: the catalytic promiscuity of plant enzymes frequently drives unwanted byproduct formation, whereas the lack of plant-like spatial partitioning in microbial hosts impedes precise control of enzyme activity and pathway coordination. Accordingly, engineering key enzymes such as T1OH and T5OH to improve substrate specificity represents an urgent challenge for heterologous baccatin III production. Notably, our results demonstrate that the promiscuity of T1OH and T5OH can be substantially attenuated via single amino acid substitutions (Figures 4 and 5), providing engineered enzyme candidates for the *de novo* microbial synthesis of baccatin III.

To recapitulate the spatial compartmentalization of plant biosynthetic pathways, we implemented a yeast–*E. coli* microbial relay strategy for baccatin III biosynthesis (Figure 6). Yeast was engineered for *de novo* synthesis of baccatin VI, which requires multiple cytochrome P450-mediated oxidation reactions. *E. coli* was deployed for late-stage pathway transformations, as deacetylases exhibit broad substrate promiscuity and acid sensitivity, which are incompatible with the baccatin VI biosynthetic cascade and the acidic intracellular environment of yeast. This labor division mimics the compartmentalized nature of plant specialized metabolism and offers a generalizable strategy for reconstructing biosynthetic pathways that cannot be efficiently reconstituted within a single cellular chassis. Collectively, these approaches enabled the complete microbial biosynthesis of baccatin III.

Beyond protein engineering of rate-limiting enzymes, spatiotemporal precise regulation of biosynthetic networks is required to mitigate byproduct generation and improve the biosynthesis efficiency of target products. As the C2 benzoyl moiety suppresses T1OH-catalyzed C1 hydroxylation; accordingly, TBT-mediated C2 benzoylation should follow the C1 hydroxylation. Tuning the timing of TBT-catalyzed C2 benzoylation—via modulation of TBT expression or spatial segregation of TBT from the shared precursor 7β-acetoxytaxusin-2α-ol (**12**) can channel this common precursor toward the baccatin III biosynthetic pathway. Furthermore, the construction of taxoid-specific biosensors facilitates real-time detection of pathway intermediates and byproducts, enabling refined spatiotemporal control over biosynthetic networks.

Although further optimization of enzyme properties and interspecies metabolic coordination is still required to reach industrially viable titers, our work bridges a major gap in paclitaxel biosynthesis and establishes a framework for engineering distributed microbial consortia to produce paclitaxel and other novel taxoids.

## METHODS

### Media, strains and chemicals

*Escherichia coli* TOP10 and BL21 (DE3) were used to clone and functional characterize deacetylases and 2OGDs. They were cultured in Luria-Bertani (LB) medium with appropriate antibiotics at 37 □. For *Saccharomyces cerevisiae* engineered strains, growth were performed at 30 □ in SC minimal medium (6.7g/L yeast nitrogen base without amino acids, 20g/L glucose, and the corresponding deficient amino acid) or YPD medium (10 g/Lyeast extract,20 g/L bacteriological peptone, and 20 g/L glucose).

### Mass spectrometry imaging

During the cutting stage, frozen tissue samples were fixed using three drops of distilled water. Leica CM1950 cryostat (Leica Microsystems GmbH, Wetzlar, Germany) was used to section the tissues at 25 μm thickness under -20 □. Then, the tissue sections were arranged in groups on indium tin oxide (ITO)-coated glass slides, and the slides with the tissue sections were dried via a vacuum desiccator for 30 min. Desiccated tissue sections located on ITO glass slides were sprayed through a matrix sprayer using 15 mg/mL DHB (2,5-dihydroxybenzoic acid), dissolved in 90%:10% Acetonitrile:water. The sprayer temperature was 60 □, with a flow rate of 0.12 mL/min and pressure of 10 psi. 20 passes of the matrix were applied to slides with 5 s of drying time between each pass. MALDI timsTOF MSI experiments were carried out on a prototype Bruker timsTOF flex MS system (Bruker Daltonics, Bremen, Germany) with a 10 kHz smartbeam 3D laser. Throughout the entire experiment, the laser power was set to 80%. The mass spectra data were acquired over a mass range from m/z 50-1300 Da in positive mode. The imaging spatial resolution was set to 20 μm, and each spectrum contained 400 laser shots. MALDI mass spectra were imported into SCiLS Lab software, with root mean square (RMS) normalization, and the signal intensity in each image was shown as the normalized intensity. MS/MS fragmentations from the timsTOF flex MS system in the MS/MS mode were chosen for further structural confirmation of these identified metabolites.

### Single-cell transcriptome sequence

The tissue was sliced into 1–2 mm wide strips using a fresh razor blade in a sterile petri dish, then quickly transferred into a sterile beaker containing 20 mL of enzyme solution. It was placed under vacuum in a vacuum pump for 30 minutes, after which the enzyme solution was kept in darkness at 27 □ with shaking at 60 rpm. After digestion was complete, the enzyme solution was filtered through a mesh into a 50 mL centrifuge tube, and the mesh was rinsed twice with washing buffer. The filtrate was centrifuged at 100g for 5 minutes, the supernatant was carefully removed, and the protoplasts were washed twice with washing buffer. The viability of the isolated protoplasts was determined using FDA (fluorescein diacetate) staining. The concentration of protoplasts suspension was adjusted to 3-3.5□×□105 cells/mL in washing buffer. Then loaded onto a microfluidic chip (GEXSCOPE® Single cell RNA-seq Kit, Singleron Biotechnologies) and scRNA-seq libraries were constructed according to the manufacturer’s instructions (Singleron Biotechnologies). The resulting scRNA-seq libraries were sequenced on an Illumina NovaSeq Xplus instrument with 150□bp paired end reads.

### Cloning of candidate genes

Candidate genes were cloned from *Taxus* twig cDNA with PrimeSTAR HS DNA Polymerase kit and then linked into the pTOPO-Blunt Simple Vector via pTOPO-Blunt Simple Cloning Kit. These recombinant vectors were transformed into *E. coli* TOP10 cells and were plated for selection on LB plates containing 100 μg/mL ampicillin. Finally, positive transformants were verified by colony PCR and Sanger sequencing.

### Construction of yeast strains

Our previous constructed *S. cerevisiae* YDBVI was used as the starting strain to construct the cell factory for baccatin VI. Codon-optimized genes for RgPAL, ScCCL^A294G^, CHD and KAT were synthesized by Shanghai Saiheng Biotechnology.co., LTD. PrimeSTAR HS DNA Polymerase and HiPure Gel DNA Pure Mini kits were used for PCR amplification and DNA products purification according to their manufacturer’s instructions. Yeast strain construction was based on the CRISPR-Cas9 system to achieve gene integration *via* homologous recombination The construction of yeast strains was performed as described in our previous paper^27^. For the functional characterization of T5OH mutants, these mutants respectively were introduced into yeast chassis strain YSP. A single colony of positive transformant was incubated in 4mL YPD medium and then grown 16h at 30 □, 250 rpm. Then, 100 μL culture was incubated into 10 mL YPD medium with 1 mL dodecane and then grown 4 days at 30 □, 250 rpm.100 μL dodecane was used for GC-FID analysis.

### *In vivo* feeding and *in vitro* enzymatic assays

The in vivo feeding and in vitro enzymatic assays were performed according to our previous study^27^. For in vivo feeding assays, a single colony of *S. cerevisiae* strain YCYP725A55 was incubated in 1mL YPD medium and then grown 24h at 30 □, 250 rpm. Then, 100uM substrate was fed to the culture. Metabolite was extracted with equal volume ethyl acetate (EA) at 48 h and the pooled extracts were evaporated. The residue was dissolved in EA and analyzed by LC-MS. YTCPR was used as the control strain. These *in vitro* enzymatic assays of deacetylases were conducted in 300 μL reactions of containing 100 mM potassium phosphate buffer (pH 7.4), 100 μM substrate and 1 mg *E. coli* crude protein lysate. These *in vitro* enzymatic assays of 2OGDs were conducted in 300 μL reactions of containing 100 mM potassium phosphate buffer (pH 7.4), 100 μM substrate, 1mM FeSO_4_, 1mM α-ketoglutarate and 1 mg *E. coli* crude protein lysate. The crude protein lysate from *E. coli* strain expressing empty vector was treated in parallel as control. The reaction system was mixed at 30 □ for 12 h. The reactants were extracted with equal volume EA and analyzed by HPLC or LC-MS. For the activity analysis of deacetylases and 2OGDs in different pH, buffer solution with different pH were used to obtain *E. coli* crude protein lysate and perform reaction.

### Whole-cell catalysis using *E.coli*

For whole-cell catalysis assays, a single colony of *E.coli* strain EBAIII expressing CXE16, CEX18 and T9ox was incubated in 4mL LB medium and then grown 8h at 37 □, 200 rpm. Then, 2ml culture was incubated in 200mL LB medium and then grown 2h at 37 □, 200 rpm. 1mM isopropyl β-D-thiogalactopyranoside (IPTG) was added to the culture and then grown 16h at 16 □, 110 rpm. Cells were collected by centrifuge and resuspended in 1ml LB buffered with PBS (0.1M pH 7.4). 100μM Substrate, 1mM FeSO_4_ and ketoglutarate were added to the culture and the reaction was incubated at 30 □, 250 rpm. Metabolite was extracted with equal volume EA at 24 h and the pooled extracts were evaporated. The residue was dissolved in EA and analyzed by LC-MS.

### GC-FID analysis

A GC2010 pro (Shimadzu, Suzhou, China) equipped with a flame ionization detector was used to GC-FID analysis with a TG-5MS capillary column (30 m × 0.25 mm × 0.25 µm, Thermo Fisher Scientific). The oven temperature was set at 160 □ for 2 min, increased to 200 □ at a rate of 1 □/min, then increased to 280 □ at a rate of 5 □/min and finally held 280 □ for 5 min. The temperature of inlet and detection were set at 260 □ and 300 □, respectively. The sample volume of per injection was 2 µL in split ratio of 20. The flow rate of carrier gas (nitrogen) was kept at 1.05 mL/min.

### HPLC and LC-MS analysis

HPLC analysis was performed on a Shimadzu LC-20A system (Shimadzu, Kyoto, Japan) equipped with a diode array detector, an auto-sampler, a binary pump, and a thermostatically controlled column compartment. Separation was achieved on a Welch Boltimate™ C18 column (2.7 μm, 2.1 mm × 100 mm). The mobile phase consisted of water (A) and acetonitrile (B) using a gradient elution program as follows: 0–2 min (15% B), 15 min (98% B), and 15.01–17 min (15% B). The flow rate was maintained at 0.45 mL/min, and the column compartment temperature was set to 35□. Detection was performed at 203 nm. For LC-MS analysis the equipment and methods are adopted as our previous study^27^. The chromatographic analysis was performed on a Dionex Ultimate 3000 RSLC (HPG) ultra-performance liquid chromatography system (Thermo Fisher Scientific) with a Welch BoltimateTM C18 column (2.7 μm, 2.1 mm × 100 mm). The HR-ESIMS data were obtained using a Q Exactive quadrupole orbitrap high resolution mass spectrometry with a HESI ionization. Xcalibur 4.2 software was used for compound analysis.

### Isolation and purification of standards

For the identification of products, the in vitro enzymatic assays were scaled up correspondingly. Product was extracted with EA and purified with the silica gel column using different ratios of PE and EA as eluent. The purity of the product was judged by NMR analysis. If necessary, Further purification was carried out on an Agilent 1260 Infinity II preparative liquid chromatography system equipped with a Welch Ultimate XB□C18 preparative column (5 μm, 21.2 mm × 20 mm). A gradient elution system consisting of water (A) and acetonitrile (B) was used and product was detected at 203 nm.

### NMR analysis

All ^1^H, ^13^C, 2D-COSY, 2D-NOE, 2D-HSQC, and 2D-HMBC NMR spectra were acquired on Bruker AM-400, 500-MHz or Bruker AV Neo 500-, 600-MHz spectrometers. The samples were dissolved in CDCl_3_. Data were processed and analyzed using MestReNova (version 14.1.0). ^1^H NMR chemical shifts (δ) are reported in ppm relative to TMS (δ = 0.00 ppm), CHCl_3_ (δ = 7.26 ppm). ^13^C NMR chemical shifts (δ) are reported in ppm relative to CDCl_3_ (δ = 77.16 ppm) All NMR spectra, along with key COSY, NOE, and HMBC correlations, are provided in the supplementary information.

### Gene cloning and protein purification

All the plasmids were transformed into the *E. coli* BL21 (DE3) strain. The transformed bacterial cells were grown in Luria-Bertani (LB) medium suppltied with kanamycin sulfate at 37 □ to an OD_600_ of 0.8, and then induced by 0.25 mM IPTG for 12 h at 18 □. The cells were harvested and resuspended in buffer A (20 mM Tris-HCl, pH 8.0, 100 mM NaCl) supplemented with 1 mM phenylmethane sulfonyl fluoride, 2 mM magnesium chloride, and 5 μg/mL DNase. Cells were lysed by a high-pressure cell disruptor at 15,000 p.s.i. (pounds per square inch), and the lysate was centrifuged at 20,000 × g for 45 min. The supernatant was loaded onto a Ni^2+^-NTA affinity column (Qiagen, Cat No.: 30230) and washed with buffer A plus 25 mM imidazole. Proteins were eluted by buffer A plus 250 mM imidazole and purified by gel filtration using a Superdex 200 Increase column (Cytiva, Cat No.: 28990944) in buffer A on an ÄKTA pure system. The sample was eluted at a flow rate of 0.4 mL/min. The elution profile was monitored by measuring the absorbance at 280 nm. Peak fractions were collected and concentrated for subsequent structural and biochemical studies.

### Crystallization, data collection, structure determination and molecular docking

For crystallization screening, protein was concentrated to 10 mg/mL, followed by the addition of 5.72 mM FeSO_4_·7H_2_O. The mixture was then incubated on ice for 30 min. Crystals were grown at 20 □ using the sitting-drop vapor diffusion method by mixing 0.5 μL of protein with 0.5 μL of reservoir solution containing 0.2 M Ammonium fluoride, 20% (w/v) polyethylene glycol 3350. All crystals were transferred into cryoprotectant solution containing their respective mother liquor plus 30% (v/v) glycerol, before being flash frozen in liquid nitrogen for storage. Data collection was performed at 02U1 beamline of the Shanghai Synchrotron Radiation Facility (SSRF) under 100 K liquid nitrogen stream (wavelength = 0.9798 Å). The data were processed with HKL3000^69^, and the initial phase was determined by molecular replacement. All of the models were manually built in Coot^70^, and were refined by iterative rounds of manual adjustment with Coot and refinement with Phenix^71^. Molecular docking between the proteins and the ligands was performed using Autodock Vina software version 1.1.2 (https://github.com/ccsb-scripps/AutoDock-Vina). The potential key amino acid sites were predicted based on the binding positions. PyMOL and Adobe Photoshop software were utilized to enhance and present the images.

## ACKNOWLEDGEMENTS

This work was financially supported by the National Key Research and Development Program of China (Grant Nos. 2023YFA0915500), the Key Research and Development Program of Guangdong (Grant Nos. 2024B1111140001), Shanghai Municipal Science and Technology Major Project, the National Natural Science Foundation of China (Grant No. 32571470), and Shanghai Municipal Commission of Science and Technology (Grant No. 24HC2810800) as well as GsynBioT (Shanghai) Co., Ltd.

## AUTHOR CONTRIBUTIONS

C.Y., Z.L., L.Y., W. W., J.L., and B. F. performed the biological experiments; C.Y. and Y.W. carried out the chemical experiments; L.Z., C.Y., T.Z. Y.X. and performed the bioinformatics analyses; T.Z., J.L., P.W., and Z.Z. supervised and coordinated the experiments; C.Y., Y.W., T.Z., P.W., and Z.Z. wrote the manuscript. C.Y., Z.L., L.Y., and Y.W. contributed equally to this work.

## COMPETING INTERESTS

These authors declare no competing interests.

## Notes

### Competing Interest Statement

The authors have declared no competing interest.

