## Supplementary material for "Discovery of a taxusin-mediated route to baccatin III enables its complete biosynthesis in engineered microbes": SI

The file includes:

Figures S1 to S80

Tables S1 to S9

NMR data

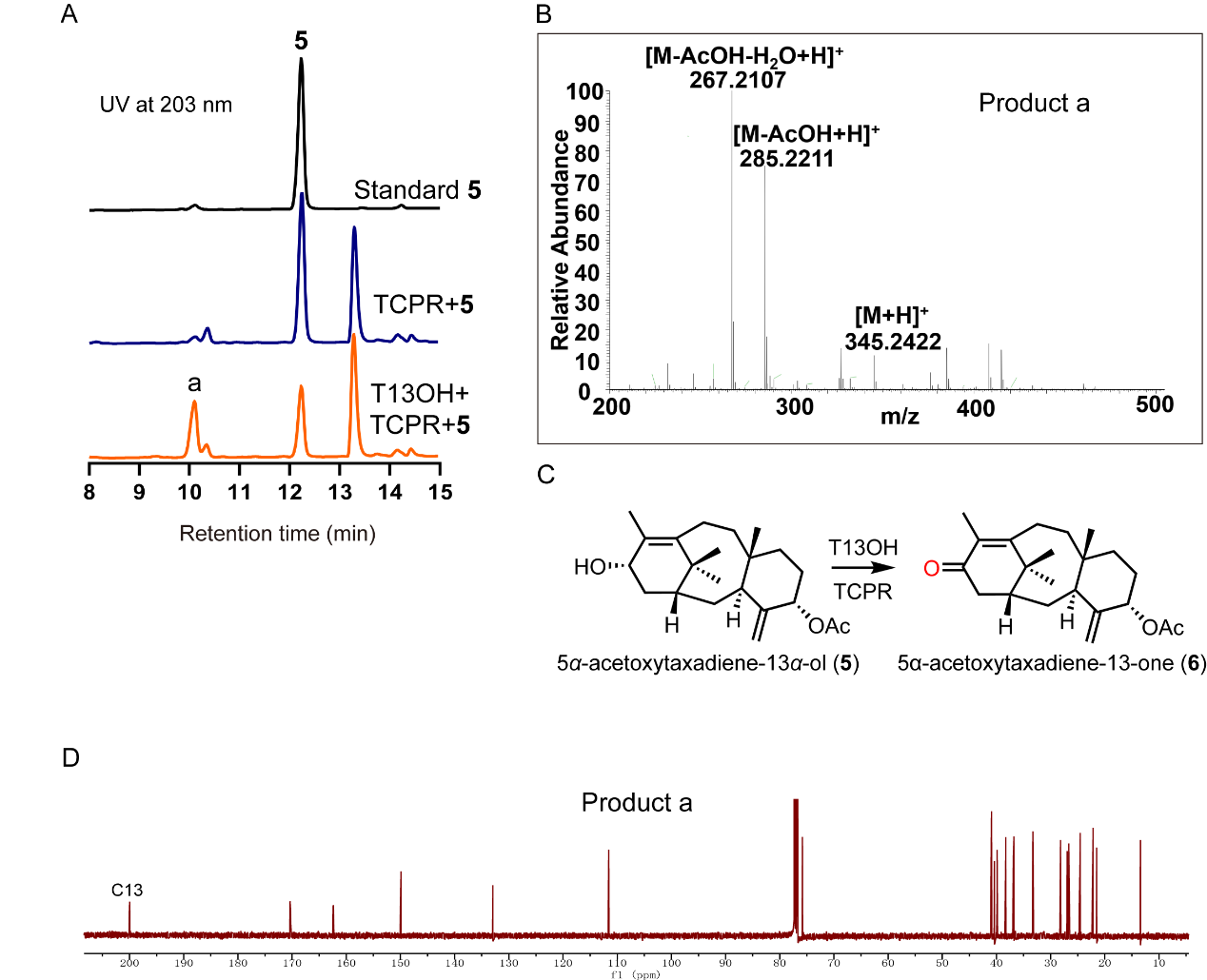

**Figure S1. T13OH oxidated C13 hydroxyl of taxoid to form corresponding C13 ketone** **taxoid, related to Figure 1.**

(A) HPLC analysis of *in vivo* feeding experiment of *S. cerevisiae* strain YT13OH expressing T13OH and TCPR using 5*α*-acetoxytaxadiene-13*α*-ol (**5**) as substrate. *S. cerevisiae* strain YTCPR expressing TCPR was used as the control.

(B) Mass spectra (ESI) of product a.

(C) T13OH could oxidate the C13 hydroxyl of 5*α*-acetoxytaxadiene-13*α*-ol (**5**) to C13 ketone taxoid 5*α*-acetoxytaxadiene-13-one (**6**).

(D) ^13^C-NMR spectrum of product a.

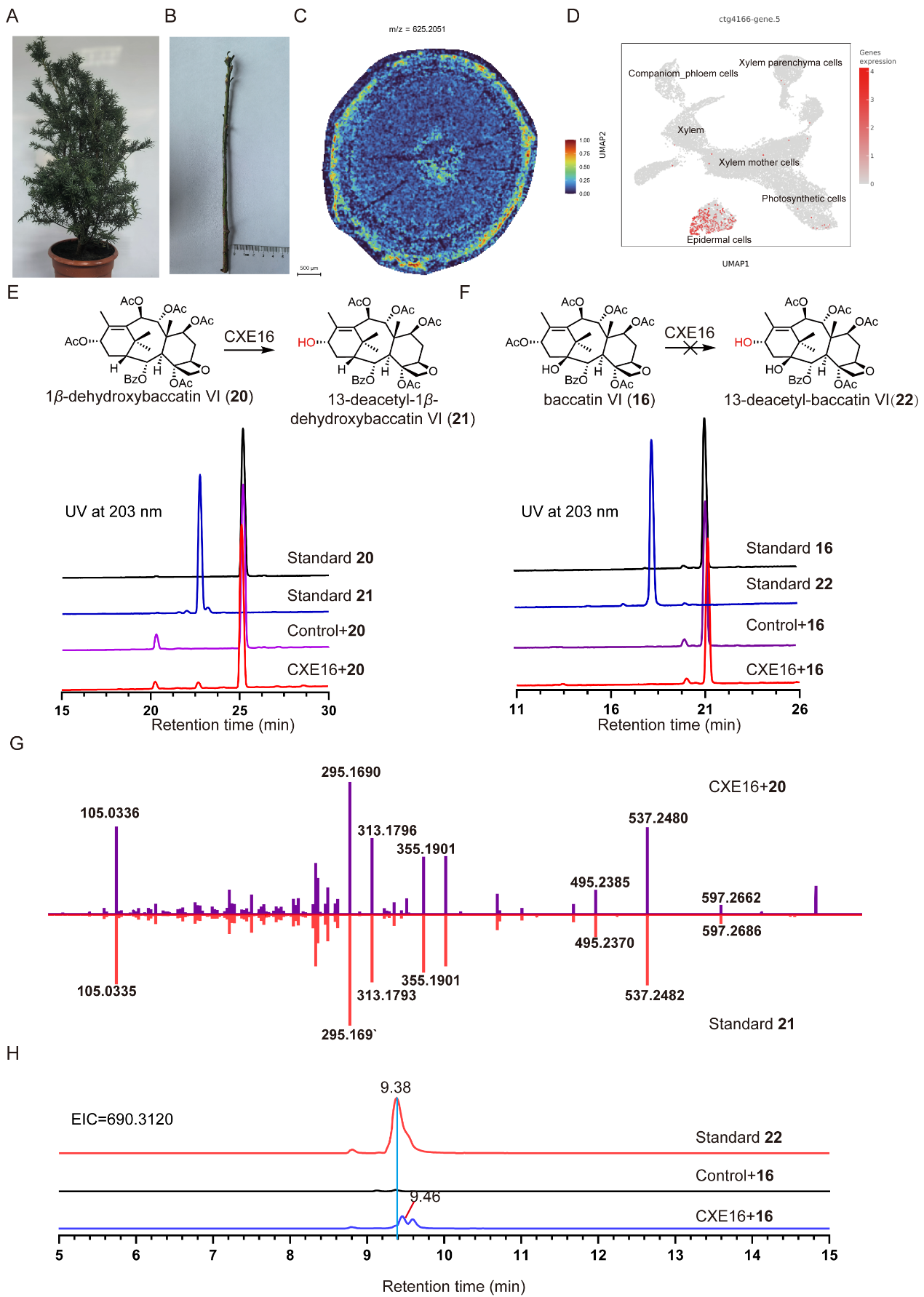

**Figure S2. Discovery and functional characterization of C13 deacetylase, related to Figure 2.**

(A-B) *Taxus* plant and stem sample used for mass spectrometry imaging and single-cell transcriptome sequence.

(C) Replicate sample for the distribution of baccatin III (**19**) in stem, detected by MALDI analysis. The color scale ranges from 0 to 1.

(D) The expression distribution of T9ox (ctg4166-gene.5) in the cell clusters.

(E) HPLC analysis of *in vitro* reaction using crude enzyme of *E. coli* expressing CXE16 with 1*β*-dehydroxybaccatin VI (**20**) as substrate. Crude enzyme of *E. coli* expressing empty vector was used as control.

(F) HPLC analysis of *in vitro* reaction using crude enzyme of *E. coli* expressing CXE16 with baccatin VI (**16**) as substrate. Crude enzyme of *E. coli* expressing empty vector was used as control.

(G) MS/MS fragmentation patterns of product from *in vitro* reaction using crude enzyme of *E. coli* expressing CXE16 with 1*β*-dehydroxybaccatin VI (**20**) as substrate compared to that of 13-deacetyl-1*β*-dehydroxybaccatin VI (**21**) standard.

(H) LC-MS analysis of *in vitro* reaction using crude enzyme of *E. coli* expressing CXE16 with baccatin VI (**16**) as substrate. Crude enzyme of *E. coli* expressing empty vector was used as control, EIC = 690.3120.

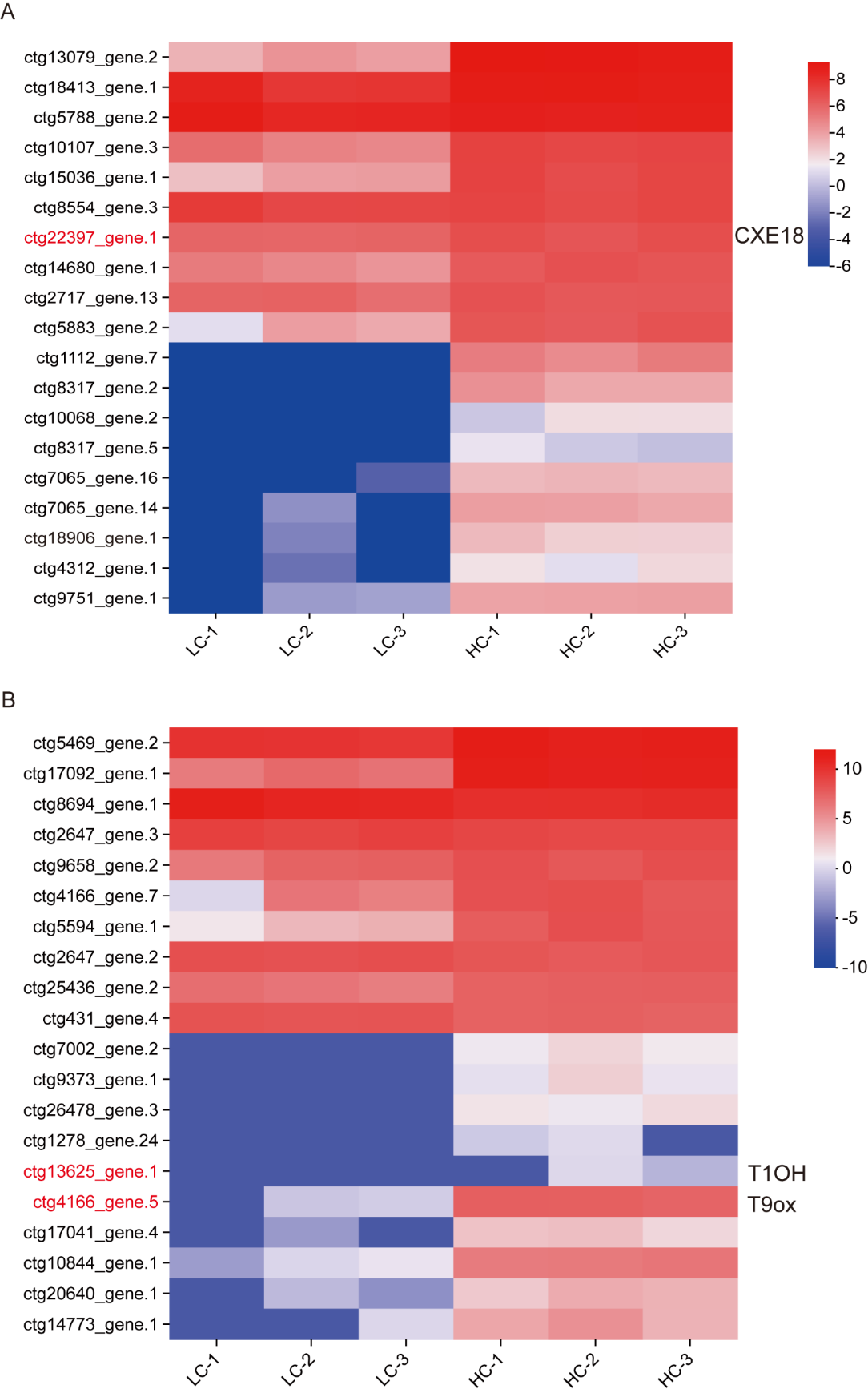

**Figure S3. Deacetylase and 2OGD candidate genes, related to Figure 2.**

(A) 19 *α*/*β*-hydrolases that were highly expressed in high Taxol-yielding cell line or upregulated compared with low Taxol-yielding cell line^1^.

(B) 20 2OGDs that were highly expressed in high Taxol-yielding cell line or upregulated compared with low Taxol-yielding cell line. HC indicates high Taxol-yielding line while LC indicates low Taxol-yielding line. The gene expression data is from reference^1^.

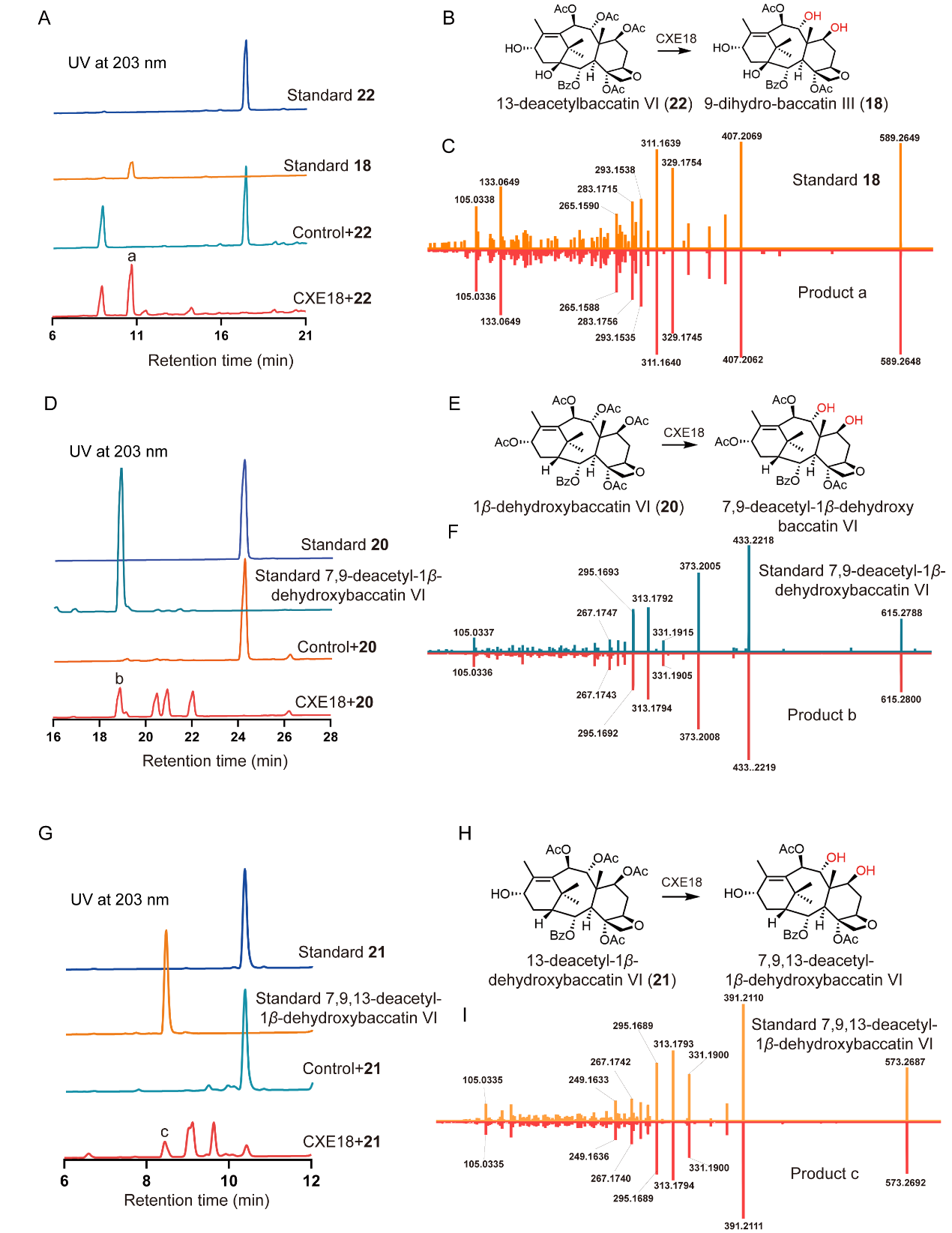

**Figure S4. Functional characterization of deacetylase CXE18, related to Figure 2.**

(A) HPLC analysis of *in vitro* reaction using crude enzyme of *E. coli* expressing CXE18 with 13-deacetylbaccatin VI (**22**) as substrate. Crude enzyme of *E. coli* expressing empty vector was used as control.

(B) CXE18 could catalyze the deacetylation at C7 and C9 of 13-deacetylbaccatin VI (**22**).

(C) MS/MS fragmentation patterns of product a from *in vitro* reaction using crude enzyme of *E. coli* expressing CXE18 with 13-deacetylbaccatin VI (**22**) as substrate compared to that of 9-dihydro-baccartin III (**18**) standard.

(D) HPLC analysis of *in vitro* reaction using crude enzyme of *E. coli* expressing CXE18 with 1*β*-dehydroxybaccatin VI (**20**) as substrate. Crude enzyme of *E. coli* expressing empty vector was used as control.

(E) CXE18 could catalyze the deacetylation at C7 and C9 of 1*β*-dehydroxybaccatin VI (**20**).

(F) MS/MS fragmentation patterns of product b from *in vitro* reaction using crude enzyme of *E. coli* expressing CXE18 with 1*β*-dehydroxybaccatin VI (**20**) as substrate compared to that of 7,9-deacetyl-1*β*-dehydroxybaccatin VI standard.

(G) HPLC analysis of *in vitro* reaction using crude enzyme of *E. coli* expressing CXE18 with 13-deacetyl-1*β*-dehydroxybaccatin VI (**21**) as substrate. Crude enzyme of *E. coli* expressing empty vector was used as control.

(H) CXE18 could catalyze the deacetylation at C7 and C9 of 13-deacetyl-1*β*-dehydroxybaccatin VI (**21**).

(I) MS/MS fragmentation patterns of product C from *in vitro* reaction using crude enzyme of *E. coli* expressing CXE18 with 13-deacetyl-1*β*-dehydroxybaccatin VI (**21**) as substrate compared to that of 7,9,13-deacetyl-1*β*-dehydroxybaccatin VI standard.

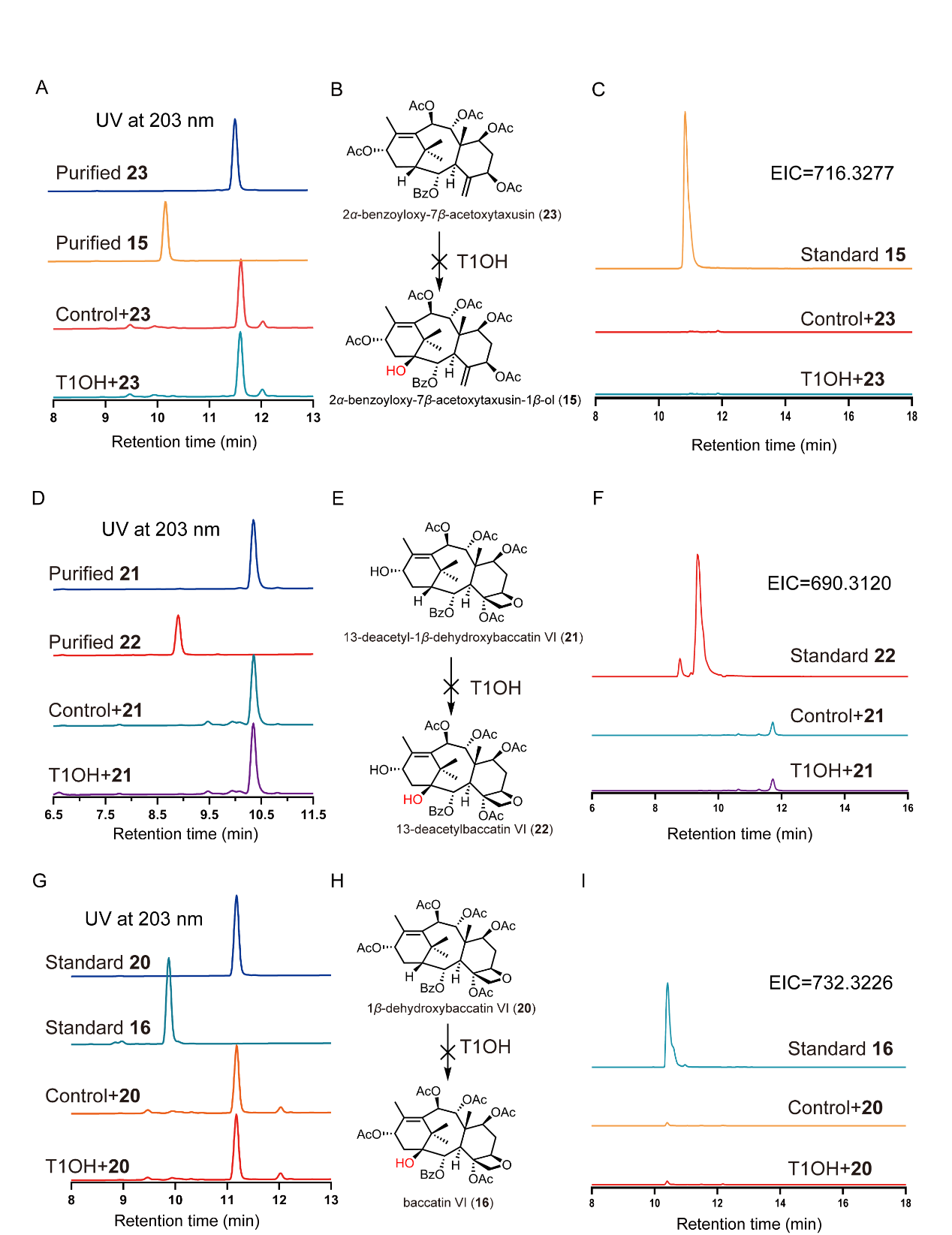

**Figure S5. Functional characterization of T1OH,** **related to Figure 3.**

(A) HPLC analysis of *in vitro* reaction using crude enzyme of *E. coli* expressing T1OH with 2*α*-benzoyloxy-7*β*-acetoxytaxusin (**23**) as substrate. Crude enzyme of *E. coli* expressing empty vector was used as control.

(B) T1OH couldn’t catalyze the hydroxylation at C1 of 2*α*-benzoyloxy-7*β*-acetoxytaxusin (**23**).

(C) LC-MS analysis of *in vitro* reaction using crude enzyme of *E. coli* expressing T1OH with 2*α*-benzoyloxy-7*β*-acetoxytaxusin (**23**) as substrate. Crude enzyme of *E. coli* expressing empty vector was used as control, EIC=716.3277.

(D) HPLC analysis of *in vitro* reaction using crude enzyme of *E. coli* expressing T1OH with 13-deacetyl-1*β*-dehydroxybaccatin VI (**21**) as substrate. Crude enzyme of *E. coli* expressing empty vector was used as control.

(E) T1OH couldn’t catalyze the hydroxylation at C1 of 13-deacetyl-1*β*-dehydroxybaccatin VI (**21**).

(F) LC-MS analysis of *in vitro* reaction using crude enzyme of *E. coli* expressing T1OH with 13-deacetyl-1*β*-dehydroxybaccatin VI (**21**) as substrate. Crude enzyme of *E. coli* expressing empty vector was used as control, EIC = 690.3120.

(G) HPLC analysis of *in vitro* reaction using crude enzyme of *E. coli* expressing T1OH with 1*β*-dehydroxybaccatin VI (**20**) as substrate. Crude enzyme of *E. coli* expressing empty vector was used as control.

(H) T1OH couldn’t catalyze the hydroxylation at C1 of 1*β*-dehydroxybaccatin VI (**20**).

(I) LC-MS analysis of *in vitro* reaction using crude enzyme of *E. coli* expressing T1OH with 1*β*-dehydroxybaccatin VI (**20**) as substrate. Crude enzyme of *E. coli* expressing empty vector was used as control, EIC = 732.3226.

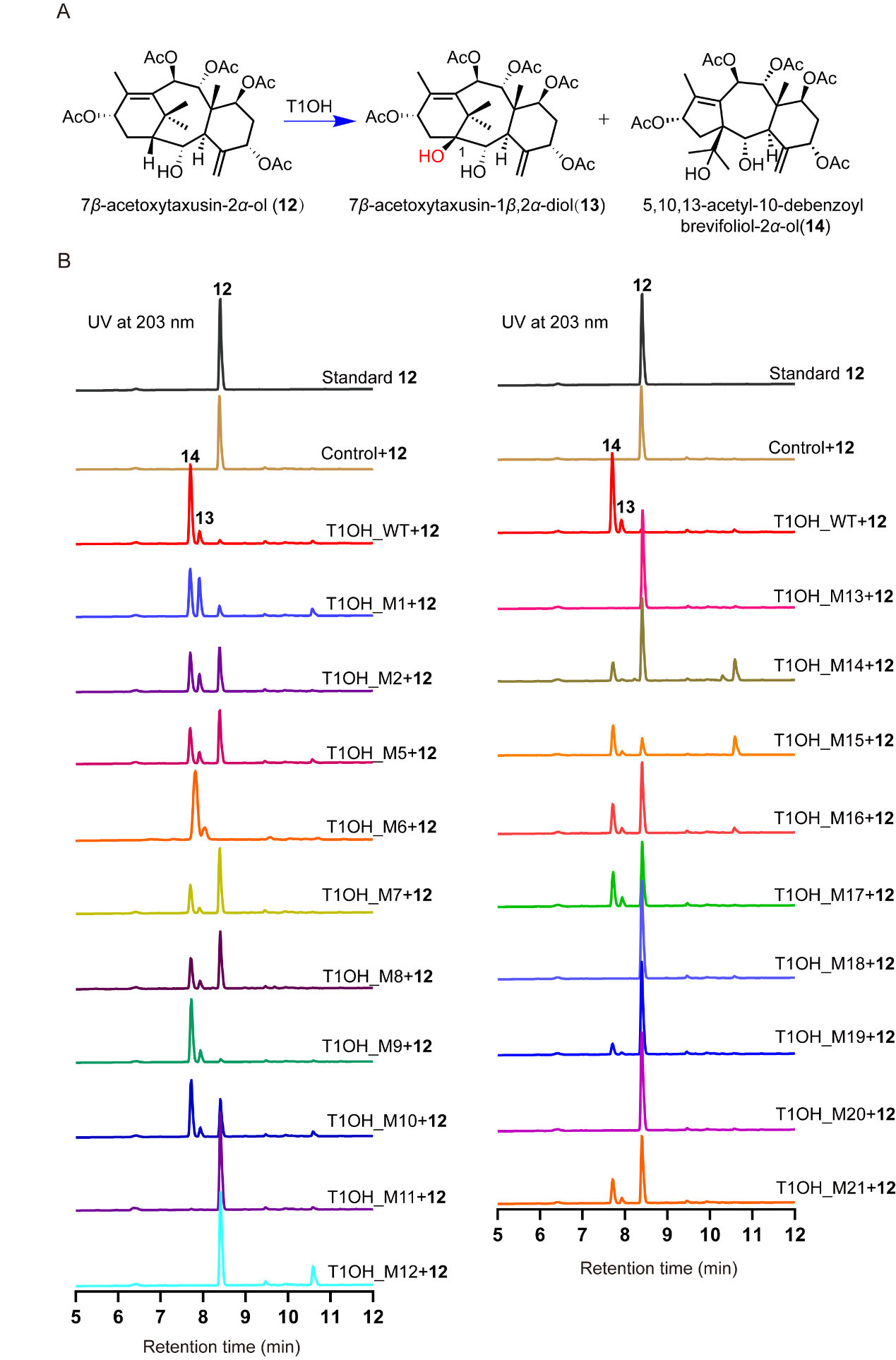

**Figure S6. Alanine scanning for candidate active sites of T1OH, related to Figure 4.**

(A) Hydroxylation reaction catalyzed by T1OH.

(B) HPLC analysis of *in vitro* reactions using crude enzyme of *E. coli* expressing each T1OH mutants with 7*β*-acetoxytaxusin-2*α*-ol (**12**) as substrate. Crude enzyme of *E. coli* expressing empty vector was used as control and crude enzyme of *E. coli* expressing wild type (WT) T1OH was used as positive control.

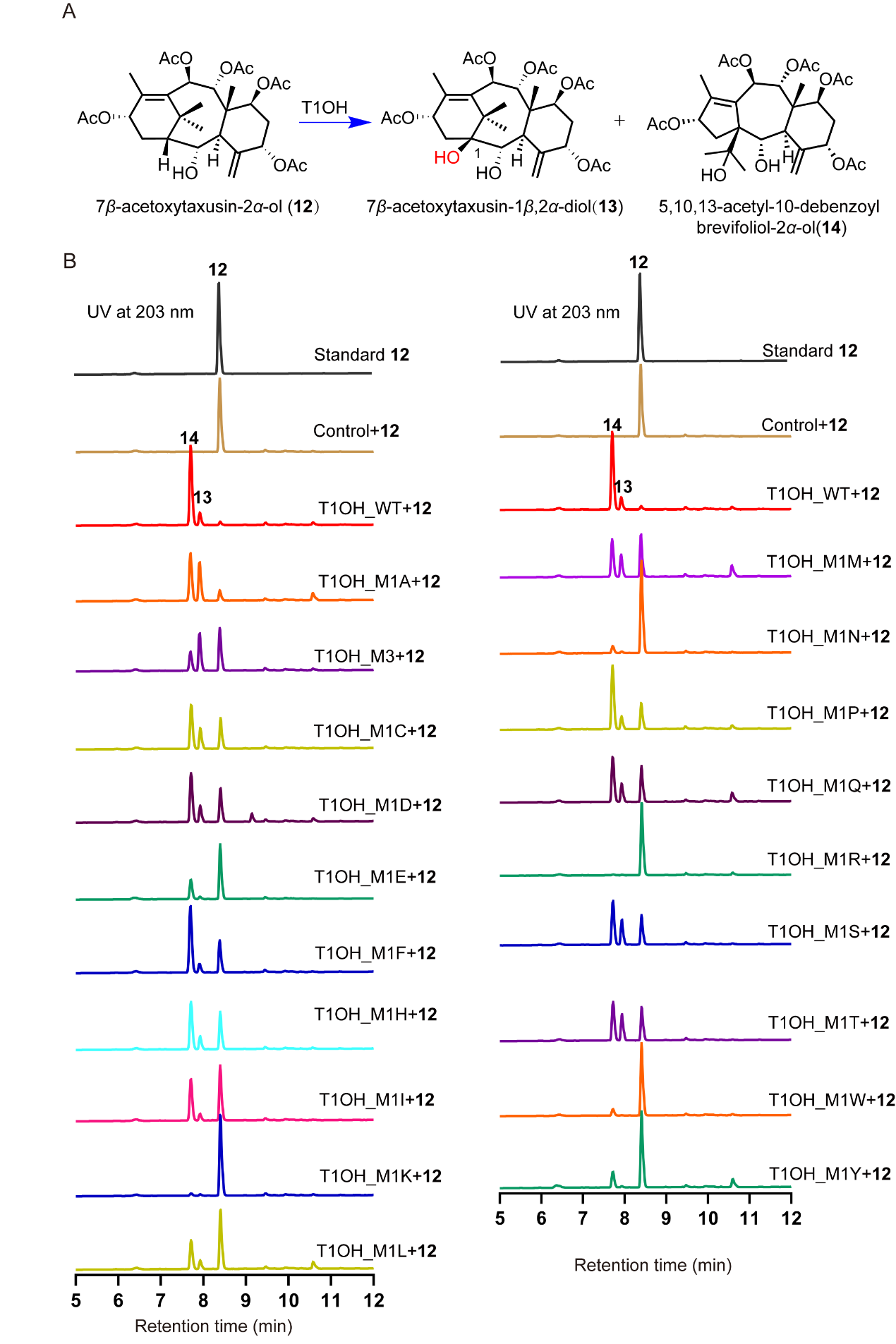

**Figure S7. Saturation mutagenesis of T1OH, related to Figure 4.**

(A) Hydroxylation reaction catalyzed by T1OH.

(B) HPLC analysis of *in vitro* reactions using crude enzyme of *E. coli* expressing different T1OH mutants with 7*β*-acetoxytaxusin-2*α*-ol (**12**) as substrate. Crude enzyme of *E. coli* expressing empty vector was used as control and crude enzyme of *E. coli* expressing wild type (WT) T1OH was used as positive control.

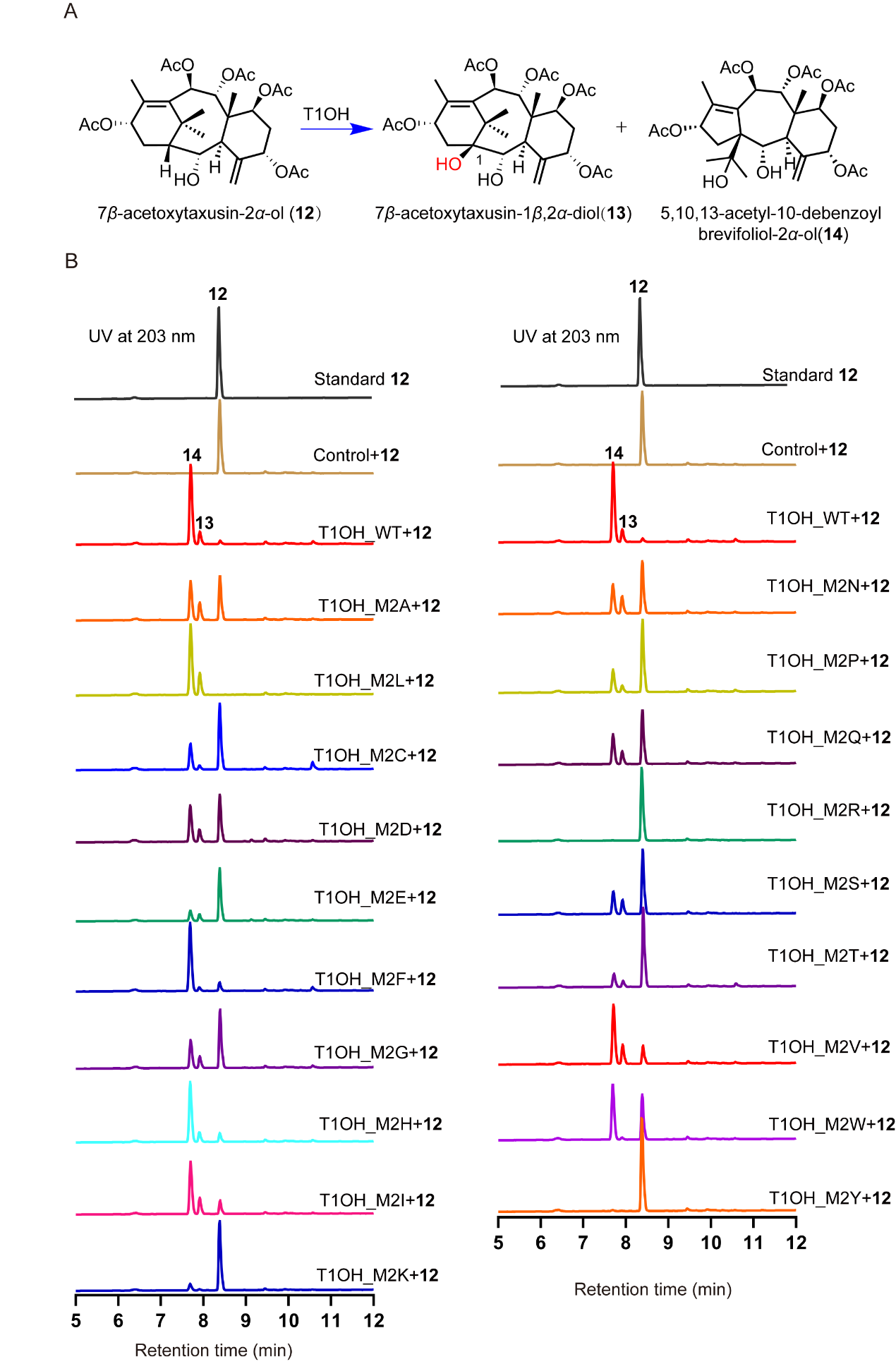

**Figure S8. Saturation mutagenesis of T1OH, related to Figure 4.**

(A) Hydroxylation reaction catalyzed by T1OH.

(B) HPLC analysis of *in vitro* reactions using crude enzyme of *E. coli* expressing different T1OH mutants with 7*β*-acetoxytaxusin-2*α*-ol (**12**) as substrate. Crude enzyme of *E. coli* expressing empty vector was used as control and crude enzyme of *E. coli* expressing wild type (WT) T1OH was used as positive control.

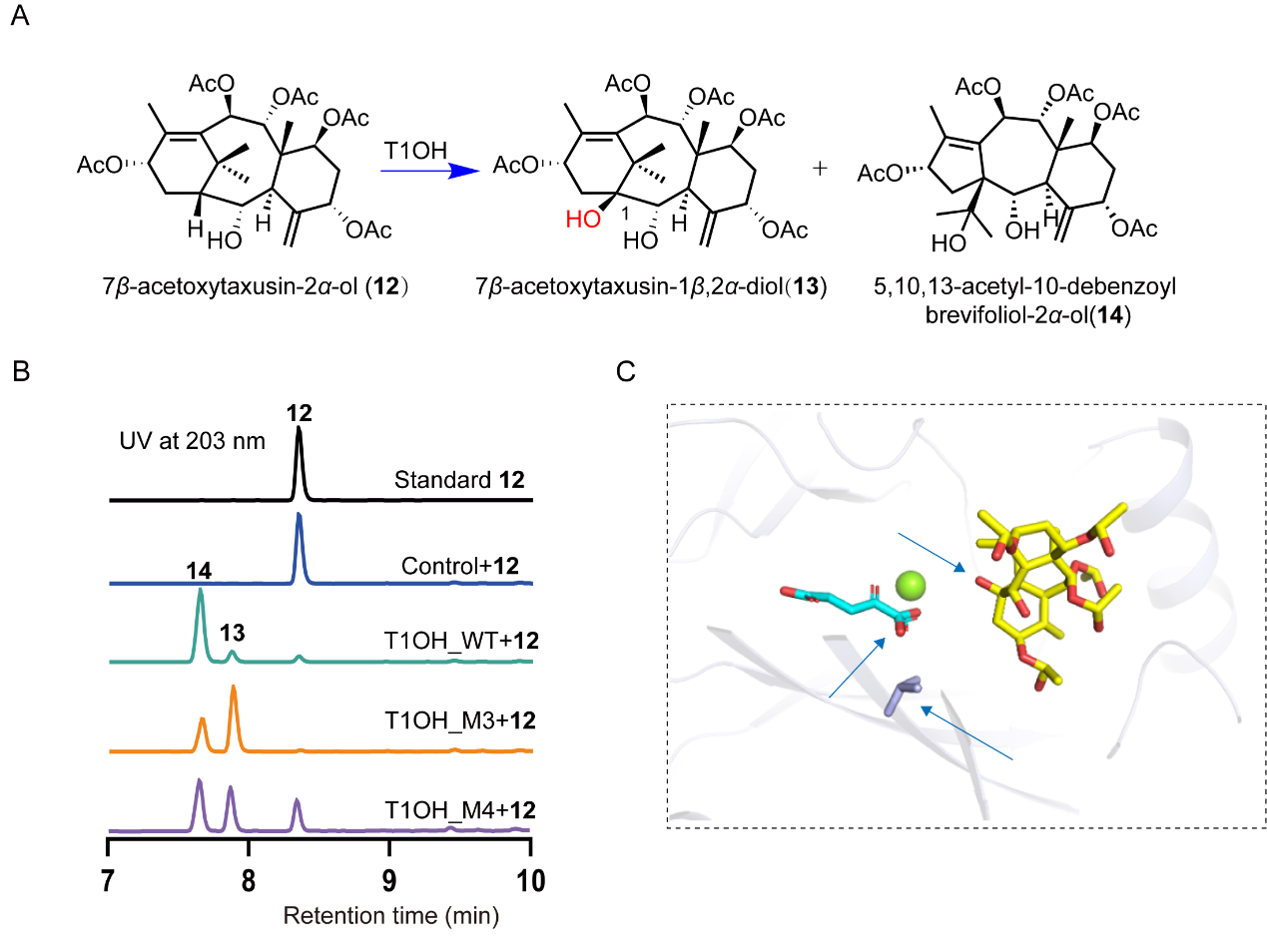

**Figure S9. Functional characterization of T1OH mutants, related to Figure 4.**

(A) Hydroxylation reaction catalyzed by T1OH.

(B) HPLC analysis of *in vitro* reactions using crude enzyme of *E. coli* expressing different T1OH mutants with 7*β*-acetoxytaxusin-2*α*-ol (**12**) as substrate. Crude enzyme of *E. coli* expressing empty vector was used as control and crude enzyme of *E. coli* expressing wild type (WT) T1OH was used as positive control.

(C) Zoomed-in view of the intermolecular interactions of T1OH with substrate 7*β*-acetoxytaxusin-2*α*-ol (**12**). The key interacting residues are identified in stick representation. The stick structures of substrate and *α*-ketoglutarate were marked with yellow and cyan, respectively.

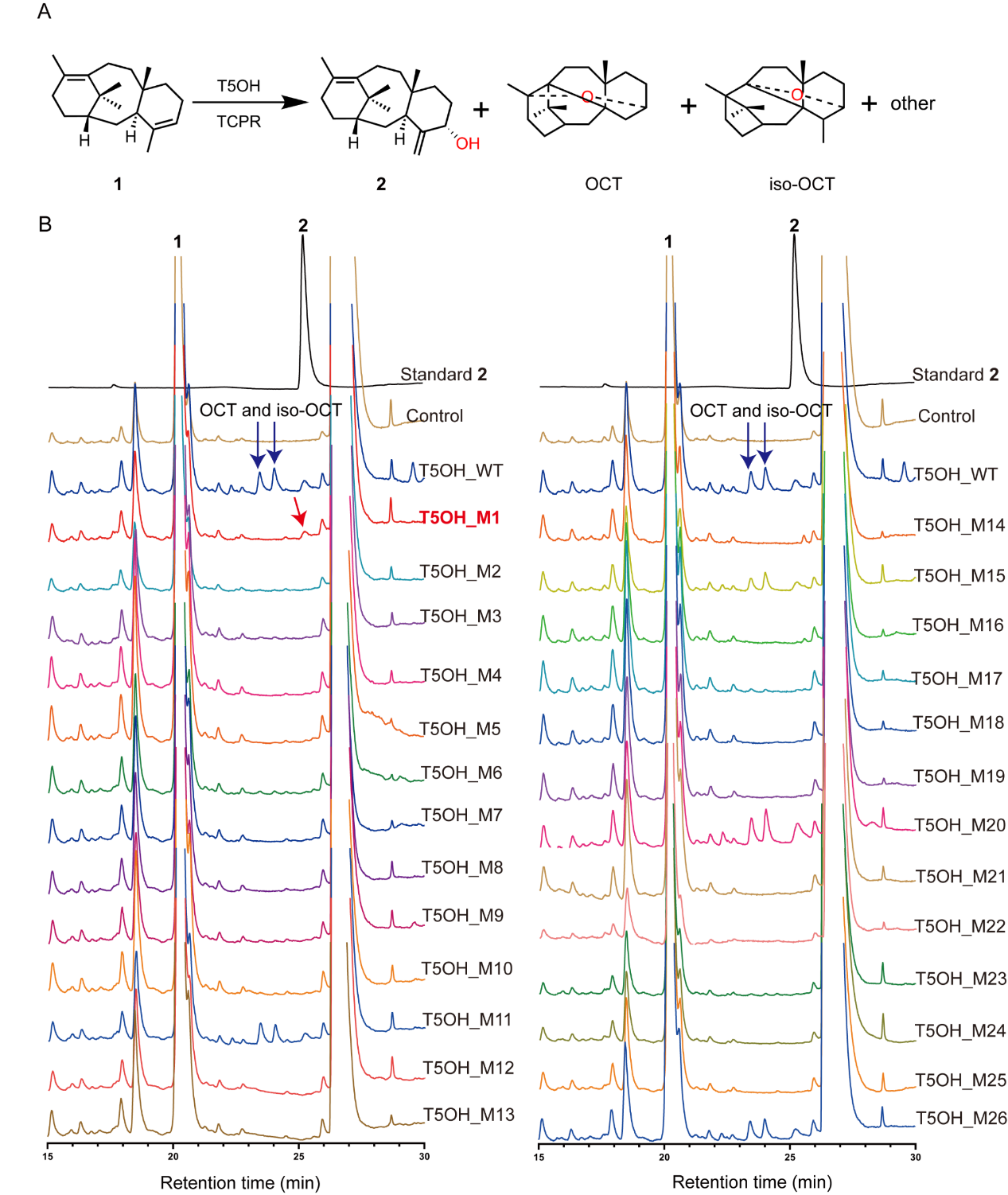

**Figure S10. Functional characterization of T5OH mutants, related to Figure 5.**

(A) Oxidation reaction catalyzed by T5OH.

(B) GC-FID analysis of products of *S. cerevisiae* cell expressing T5OH and its mutants.

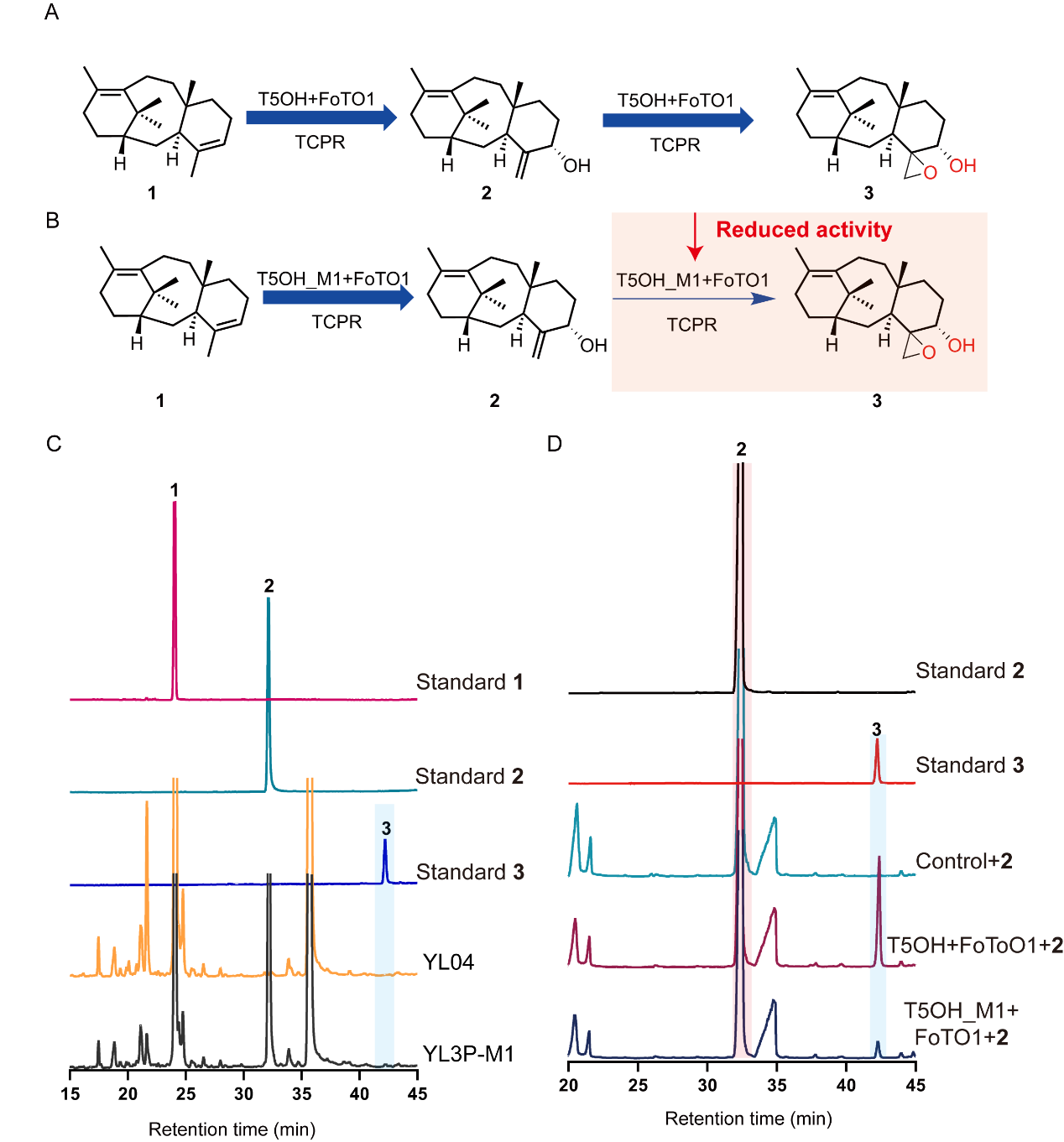

**Figure S11. Functional characterization of T5OH mutants, related to Figure 5.**

(A) Proposed process for the formation of 4*α*,20-epoxy-taxadiene-5*α*-ol (**3**).

(B) Proposed process for the reduced activity for formation of 4*α*,20-epoxy-taxadiene-5*α*-ol (**3**) with T5OH_M1.

(C) GC-FID analysis of product of *S. cerevisiae* strain YL3P-M1 expressing FoTO1 with T5OH_M1 in *S. cerevisiae* strain YL04 producing taxadiene (**1**).

(D) GC-FID analysis of products of *S. cerevisiae* strains YFT5OH and YFT5OH_M1 respectively expressing FoTO1 with T5OH or its mutation fed with taxadiene-5*α*-ol (**2**). *S. cerevisiae* strain YTCPR was used as the control.

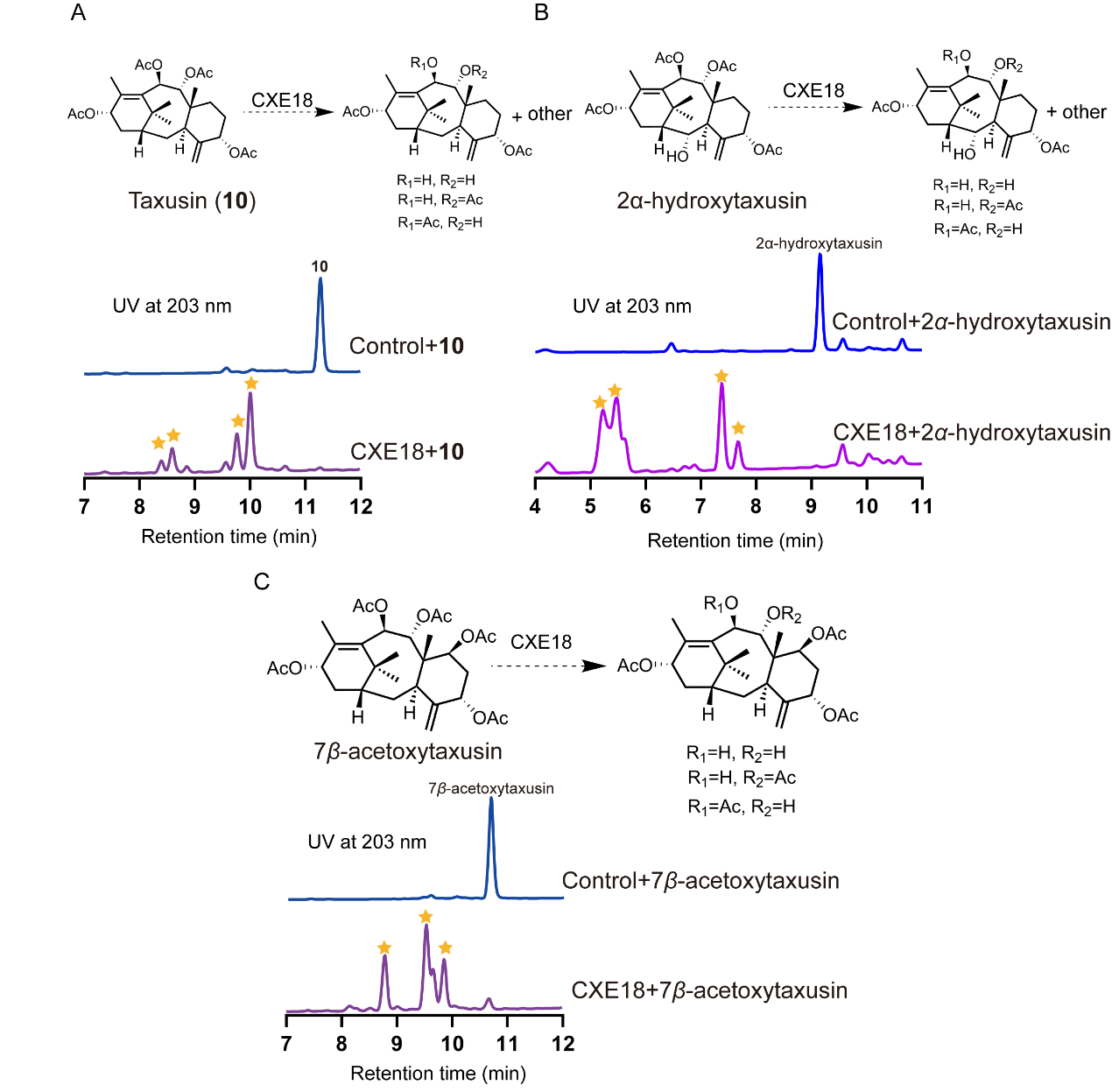

**Figure S12.** **Substrate promiscuity of CXE18, related to Figure 6.**

(A) HPLC analysis of *in vitro* reaction using crude enzyme of *E. coli* expressing CXE18 using taxusin (**10**) as substrate. Crude enzyme of *E. coli* expressing empty vector was used as control. Products were marked with yellow stars.

(B) HPLC analysis of *in vitro* reaction using crude enzyme of *E. coli* expressing CXE18 using 2*α*-hydroxytaxusin as substrate. Crude enzyme of *E. coli* expressing empty vector was used as control. Products were marked with yellow stars.

(C) HPLC analysis of *in vitro* reaction using crude enzyme of *E. coli* expressing CXE18 using 7*β*-acetoxytaxusin as substrate. Crude enzyme of *E. coli* expressing empty vector was used as control. Products were marked with yellow stars.

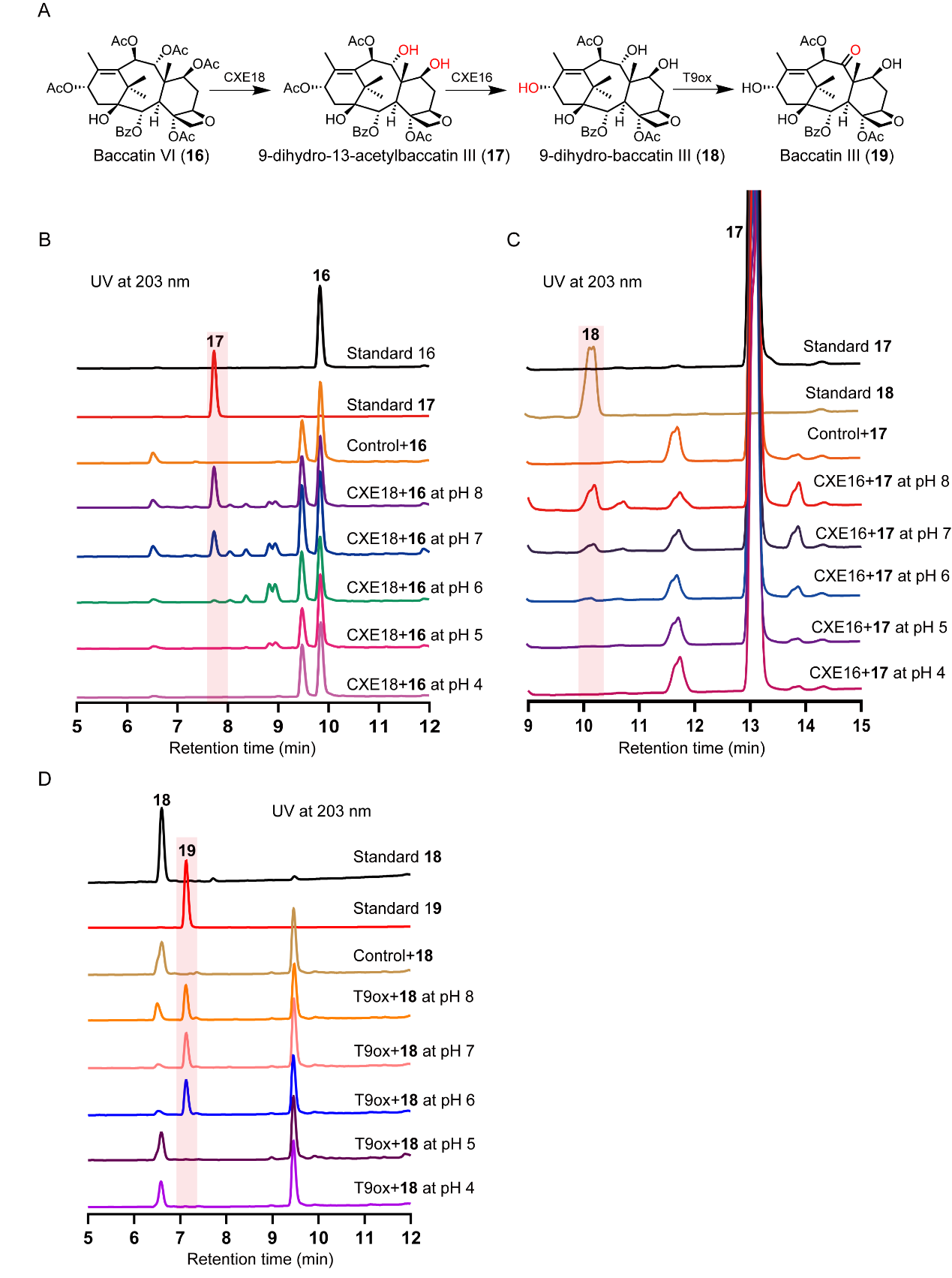

**Figure S13.** **Impact of pH on the catalytic activities of CXE18, CXE16 and T9ox, related to Figure 6.**

(A) Biotransformation of baccatin VI (**16**) to baccatin III (**19**) *via* subsequent deacetylation and oxidation catalyzed by CXE18, CXE16 and T9ox.

(B) HPLC analysis of *in vitro* reactions using crude enzyme of *E. coli* expressing CXE18 with baccatin VI (**16**) as substrate under different pH conditions. Crude enzyme of *E. coli* expressing empty vector was used as control.

(C) HPLC analysis of *in vitro* reactions using crude enzyme of *E. coli* expressing CXE16 with 9-dihydro-13-acetylbaccatin III (**17**) as substrate under different pH conditions. Crude enzyme of *E. coli* expressing empty vector was used as control.

(D) HPLC analysis of *in vitro* reactions using crude enzyme of *E. coli* expressing T9ox with 9-dihydro-baccatin III (**18**) as substrate at different pH conditions. Crude enzyme of *E. coli* expressing empty vector was used as control.

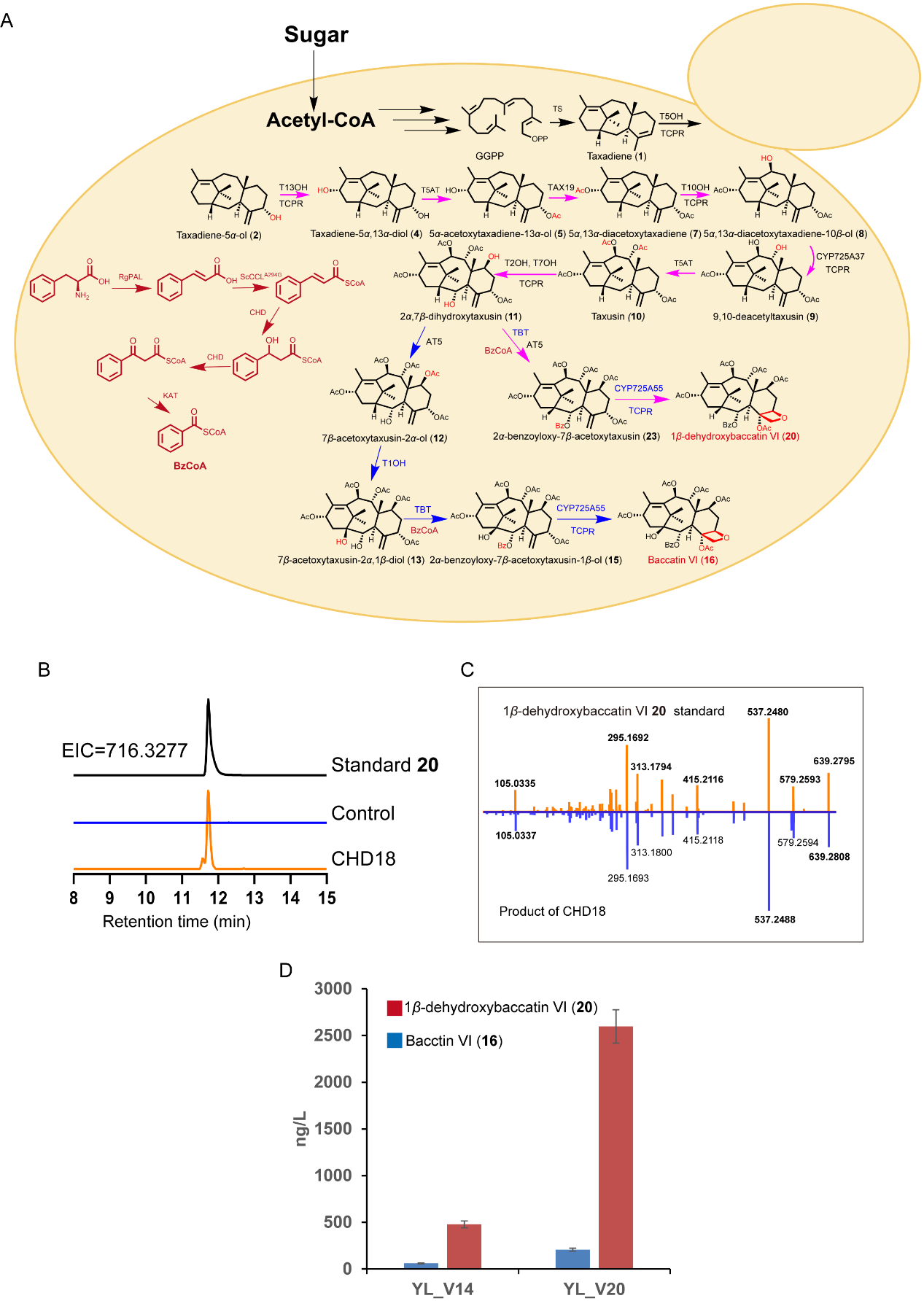

**Figure S14.** ***De novo* biosynthesis of baccatin VI in *S. cerevisiae* cell factories, related to Figure 6.**

(A) The biosynthetic pathway of baccatin VI (**16**) and byproduct 1*β*-dehydroxybaccatin VI (**20**). Reactions marked with magenta arrows were characterized and confirmed in our previous study. Reactions marked with blue arrows were characterized in this study.

(B) LC-MS analysis of product of *S. cerevisiae* cell factory CHD18 for *de novo* biosynthesis of 1*β*-dehydroxybaccatin VI (**20**). Extracted ion chromatogram (EIC) = 716.3277. *S. cerevisiae* strain CS2-Cas9 without biosynthesis of precursor taxadiene-5*α*-ol (**2**) was used as control.

(C) MS/MS fragmentation patterns of product from *S. cerevisiae* cell factory CHD18 compared to that of 1*β*-dehydroxybaccatin VI (**20**) standard.

(D) The yield of baccatin VI (**16**) and byproduct 1*β*-dehydroxybaccatin VI (**20**) in *S. cerevisiae* cell factories YL_V14 and YL_V20. Overexpression of T5OH_M1 and T1OH_M3 mutants, together with reinforcement of the biosynthesis of taxadiene (**1**) in *S. cerevisiae* cell factory YL_V14 generated *S. cerevisiae* cell factory YL_V20.

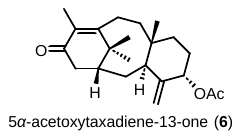

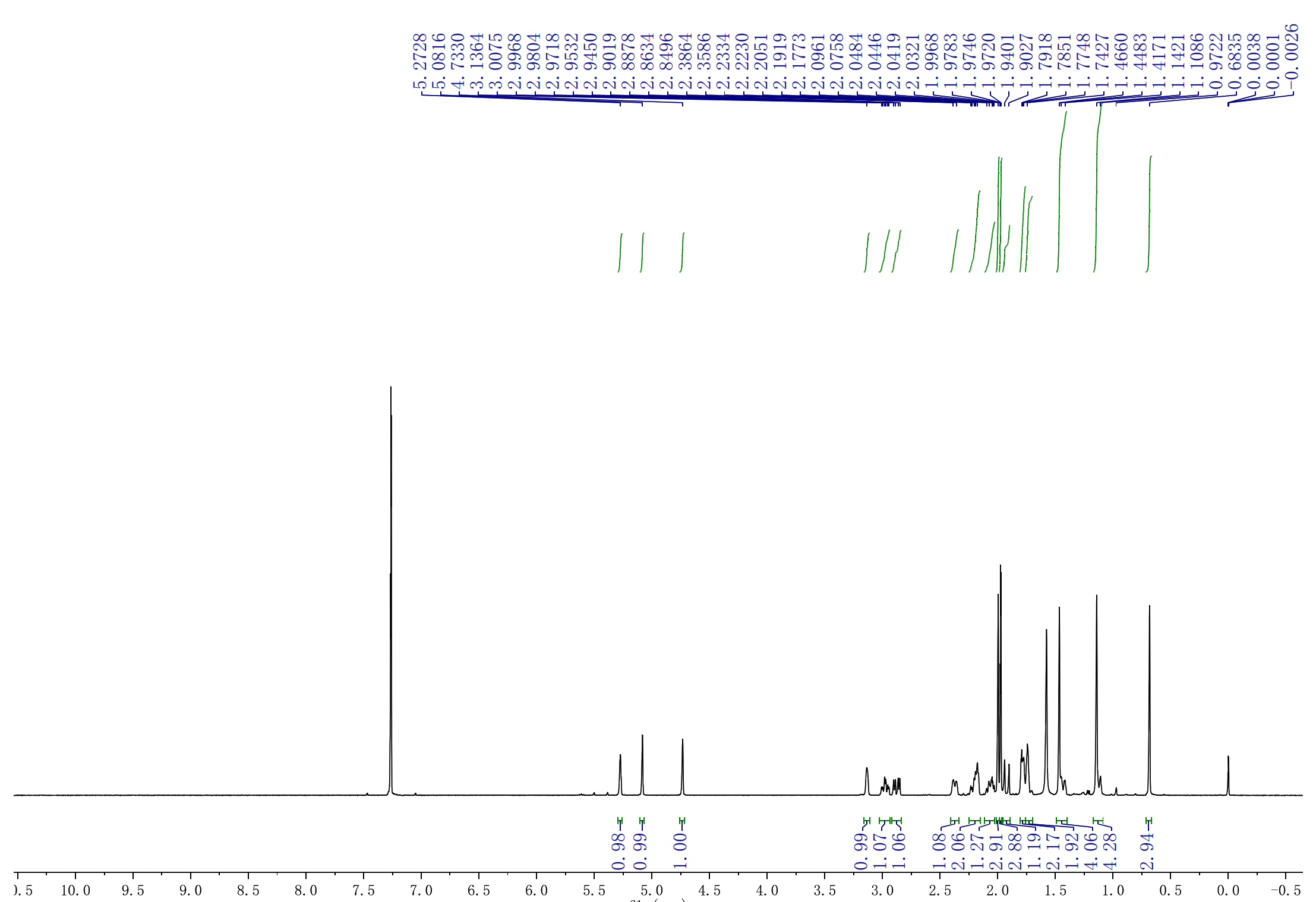

**Figure S15. ^1^H NMR spectrum of 5*α*-acetoxytaxadiene-13-one** **(6).**

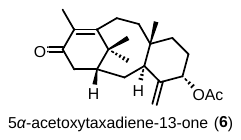

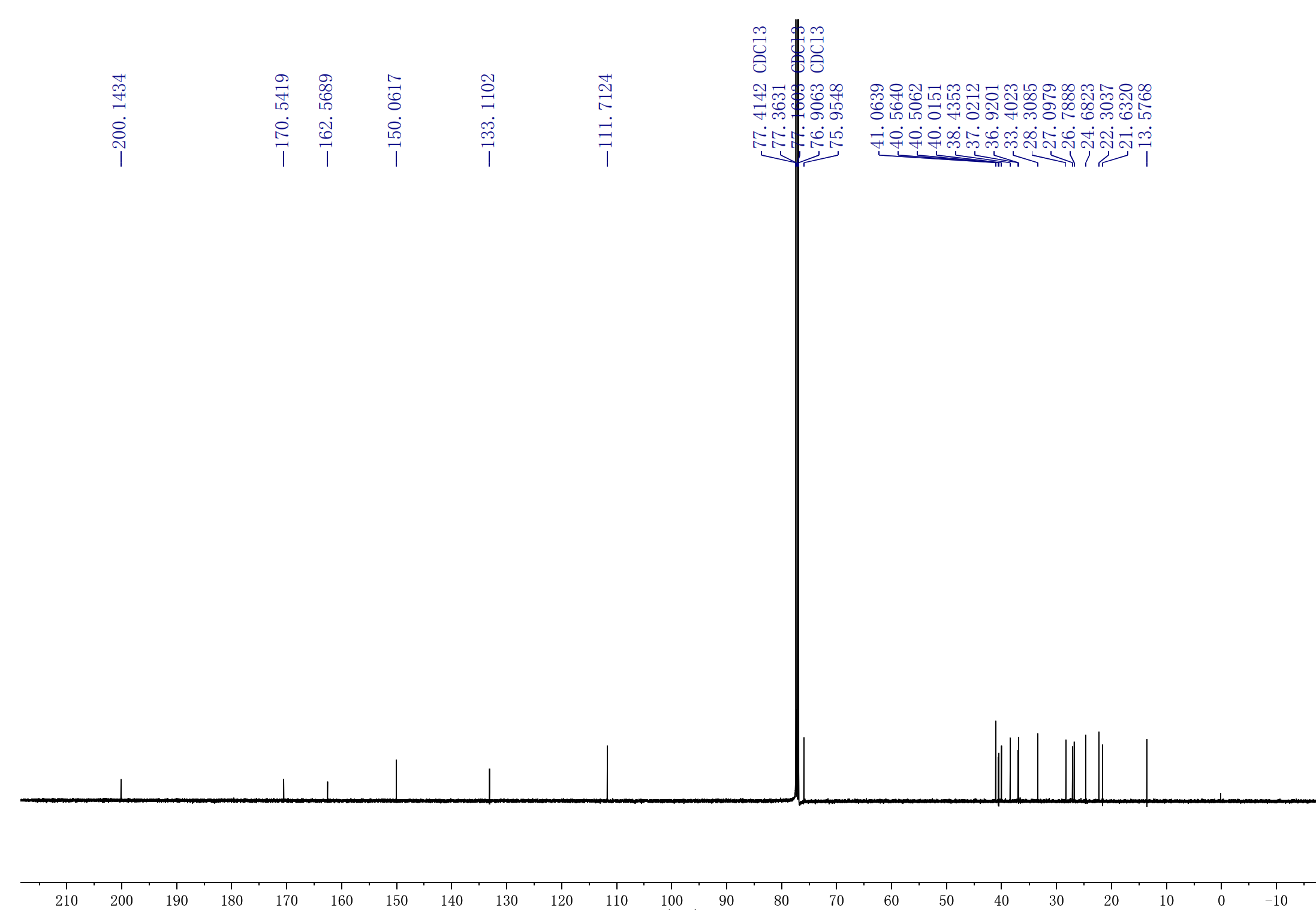

**Figure S16. ^13^C NMR spectrum of 5*α*-acetoxytaxadiene-13-one (6).**

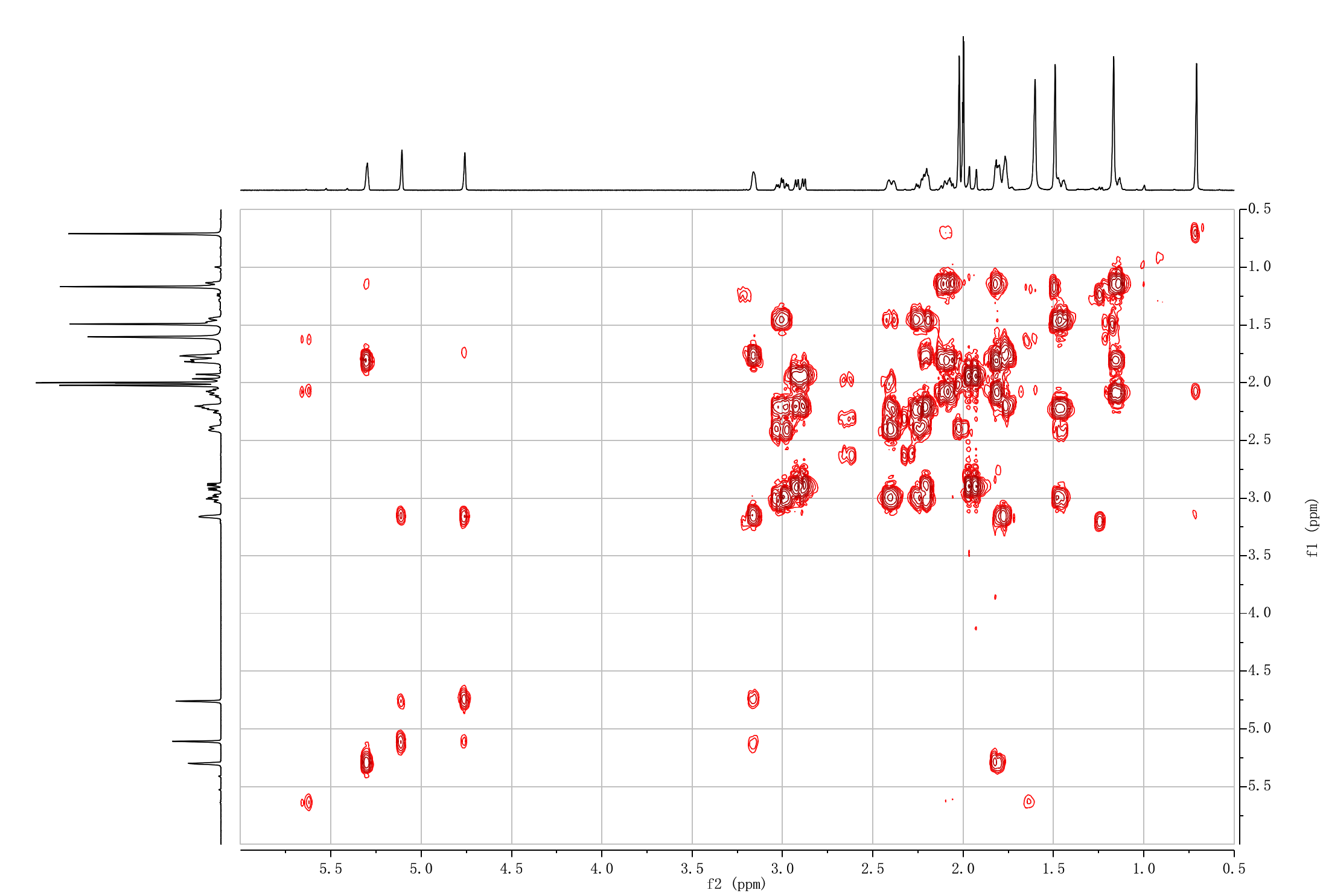

**Figure S17. ^1^H-^1^H COSY NMR spectrum of 5*α*-acetoxytaxadiene-13-one (6).**

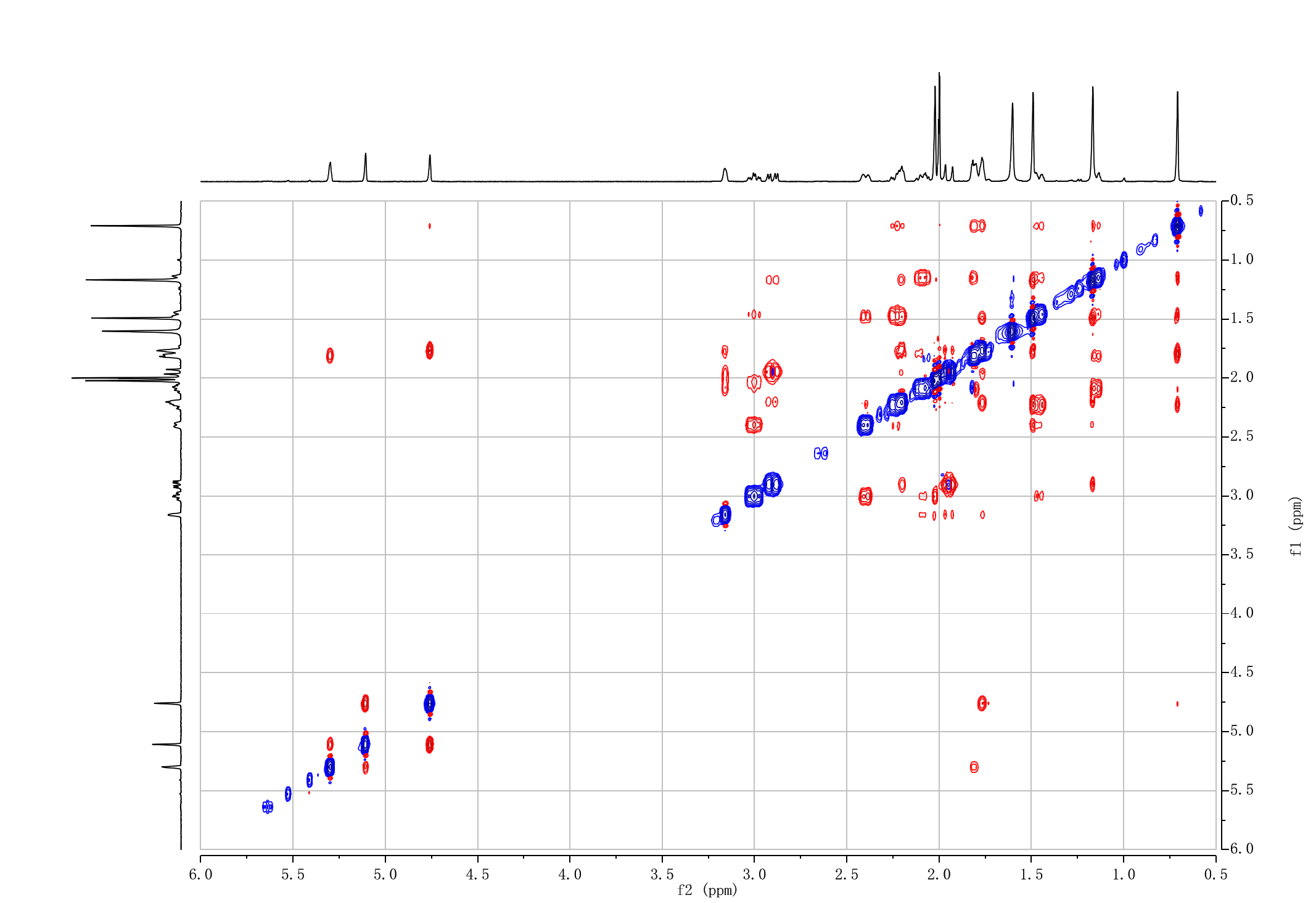

**Figure S18.^1^H-^1^H NOE NMR spectrum of 5*α*-acetoxytaxadiene-13-one (6).**

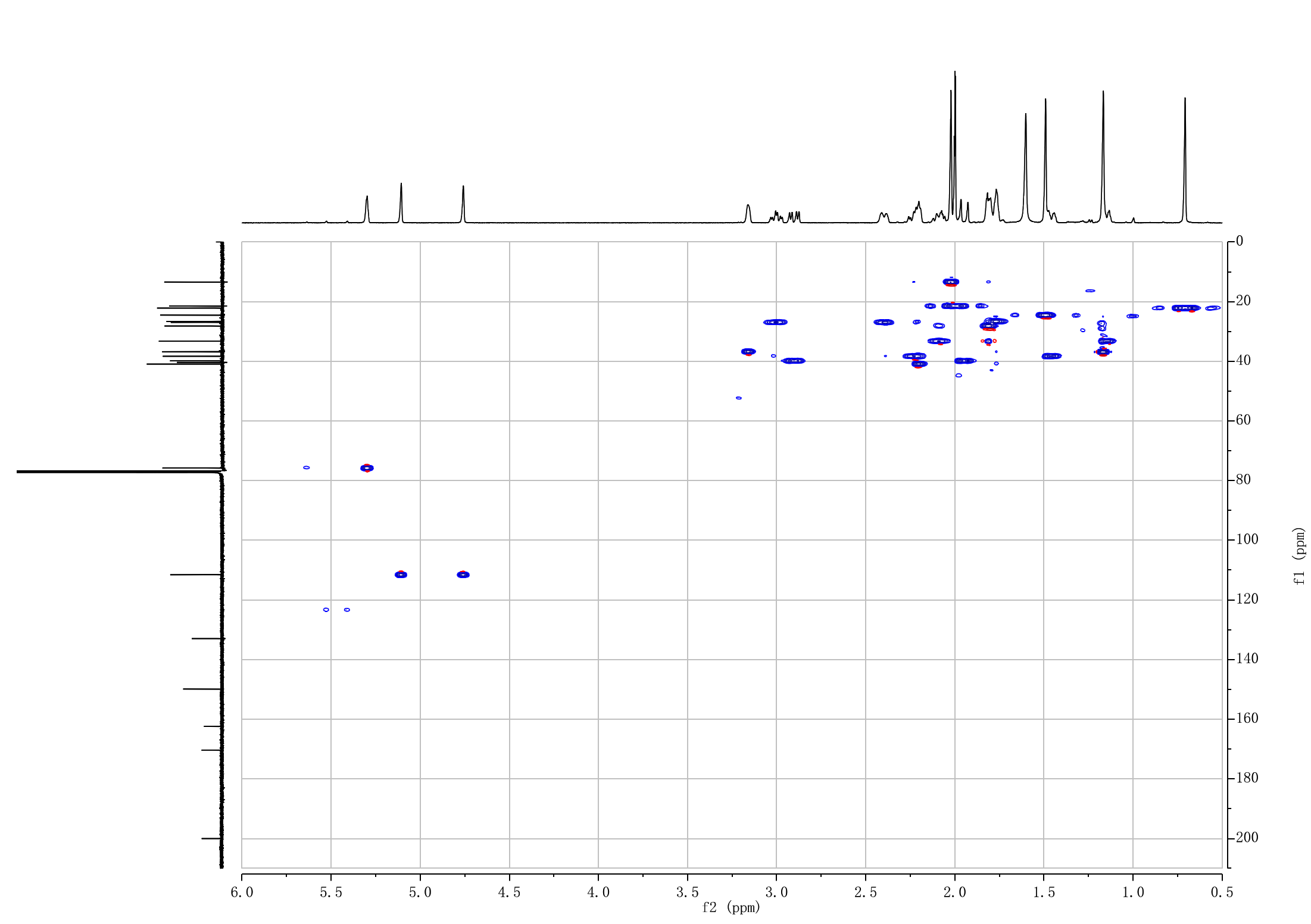

**Figure S19. ^1^H-^13^C HSQC NMR spectrum of 5*α*-acetoxytaxadiene-13-one (6).**

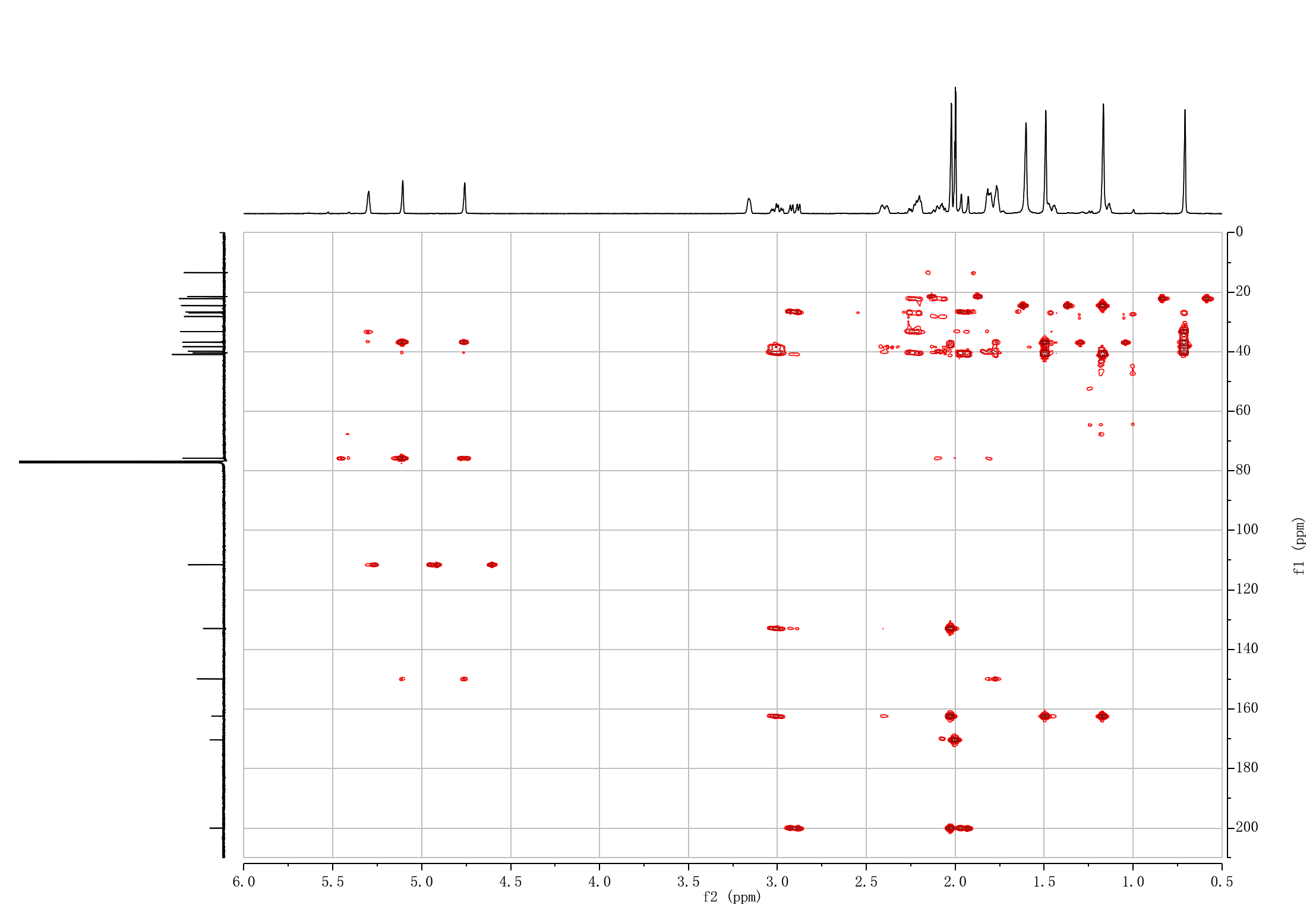

**Figure S20. ^1^H-^13^C HMBC NMR spectrum of 5*α*-acetoxytaxadiene-13-one (6).**
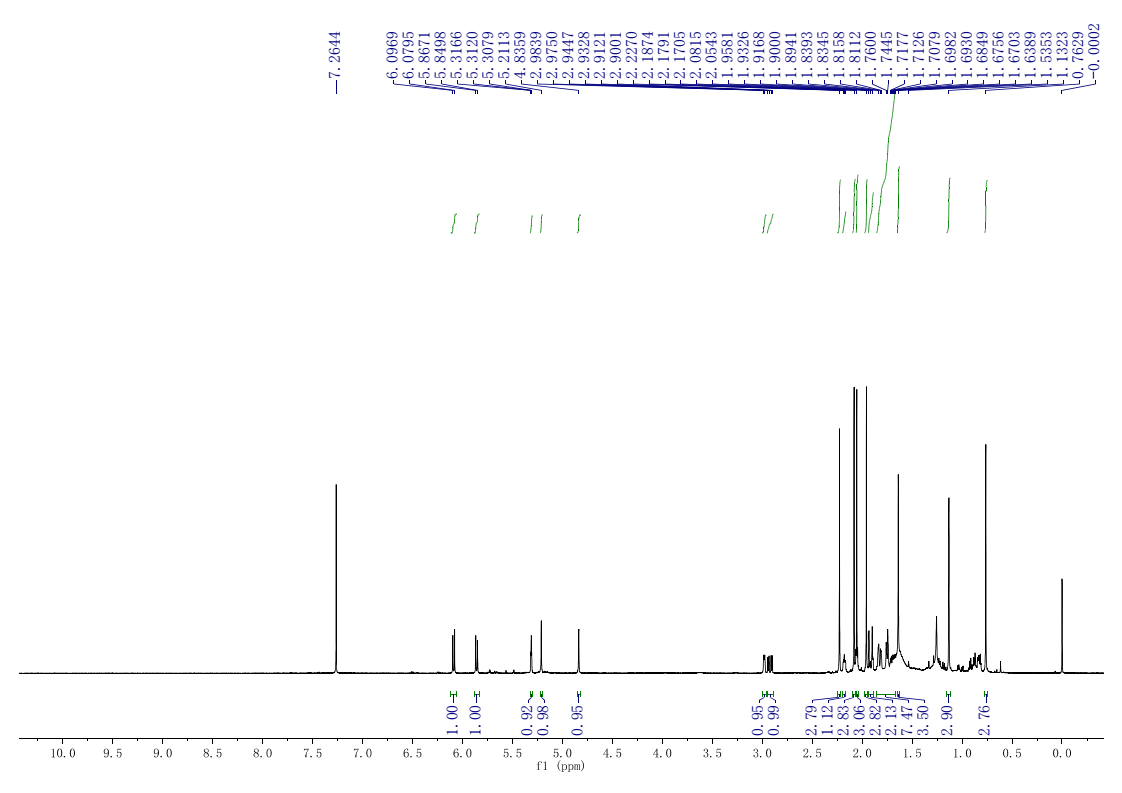

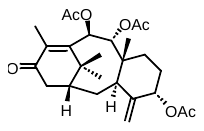

**Figure S21. ^1^H spectrum of taxusin-13-one.**

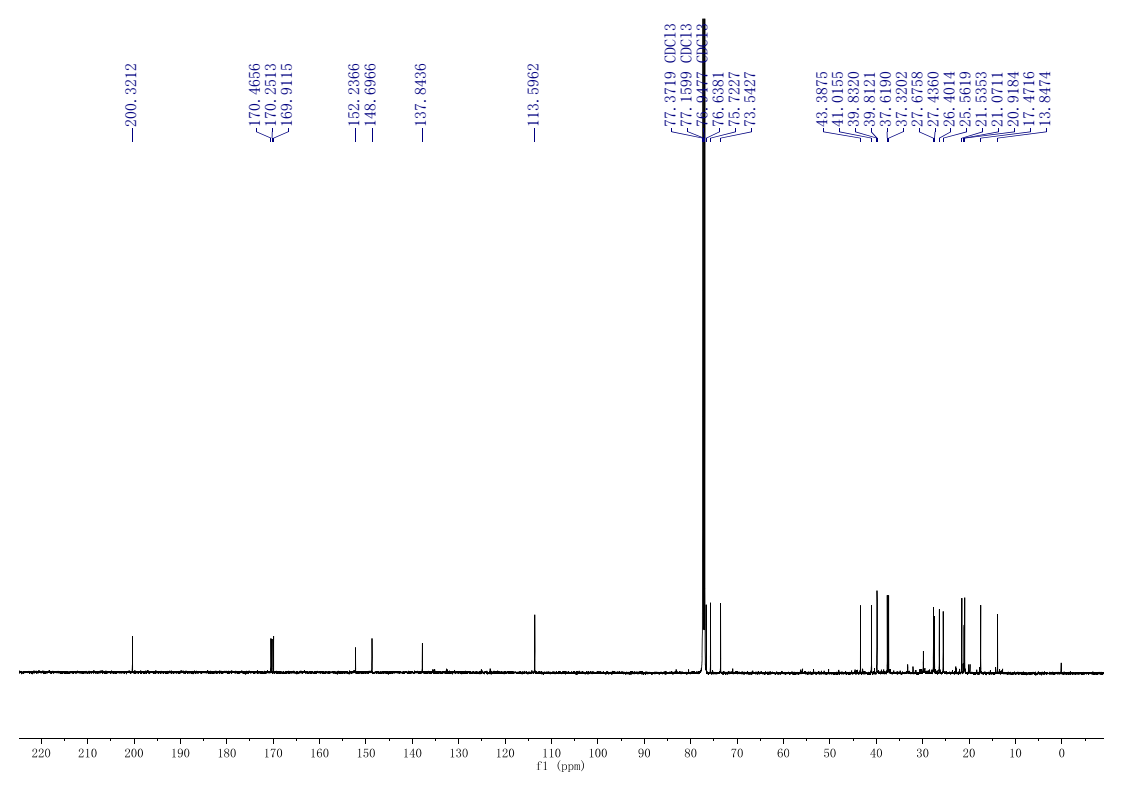

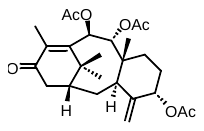

**Figure S22. ^13^C spectrum of taxusin-13-one.**

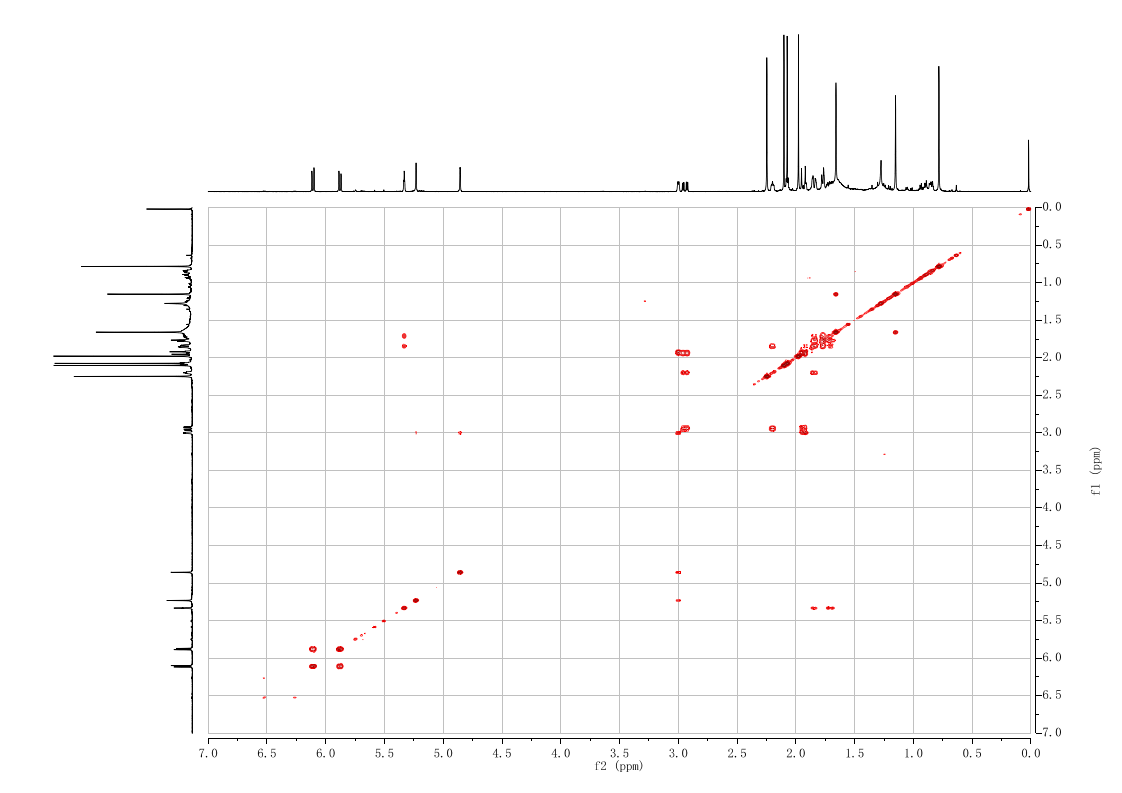

**Figure S23. ^1^H-^1^H COSY spectrum of taxusin-13-one.**

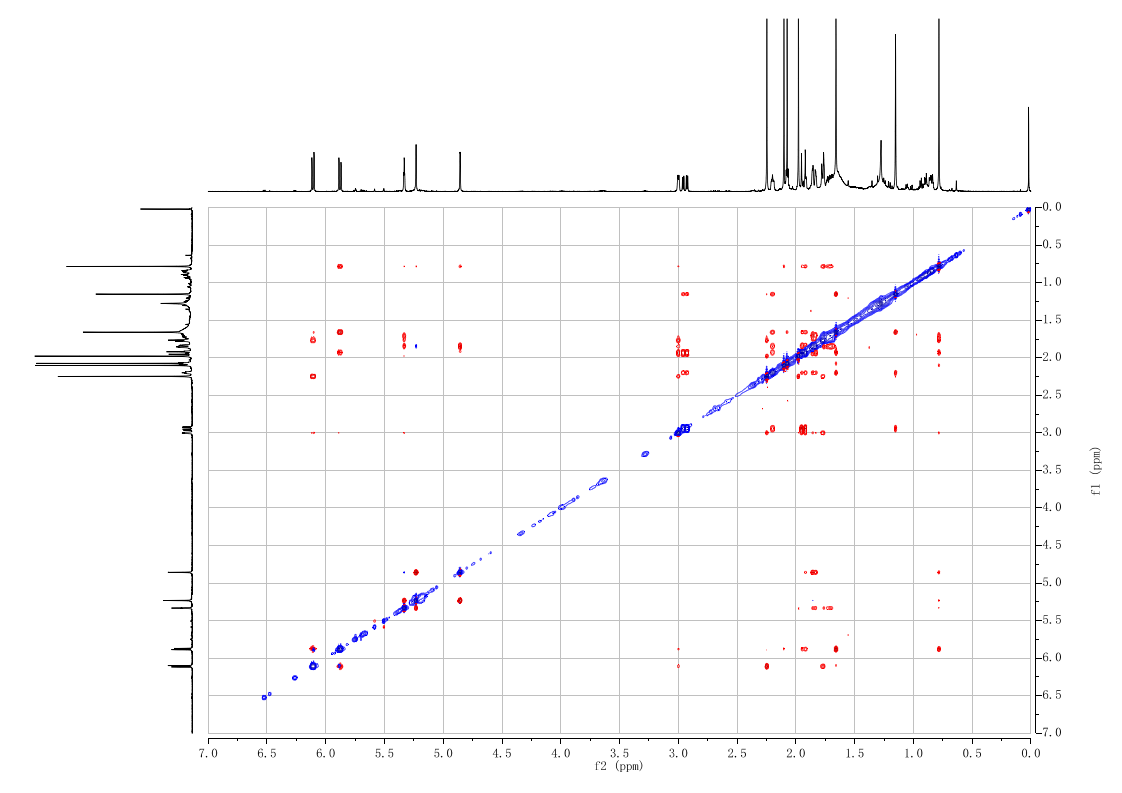

**Figure S24. ^1^H-^1^H NOE spectrum of taxusin-13-one.**

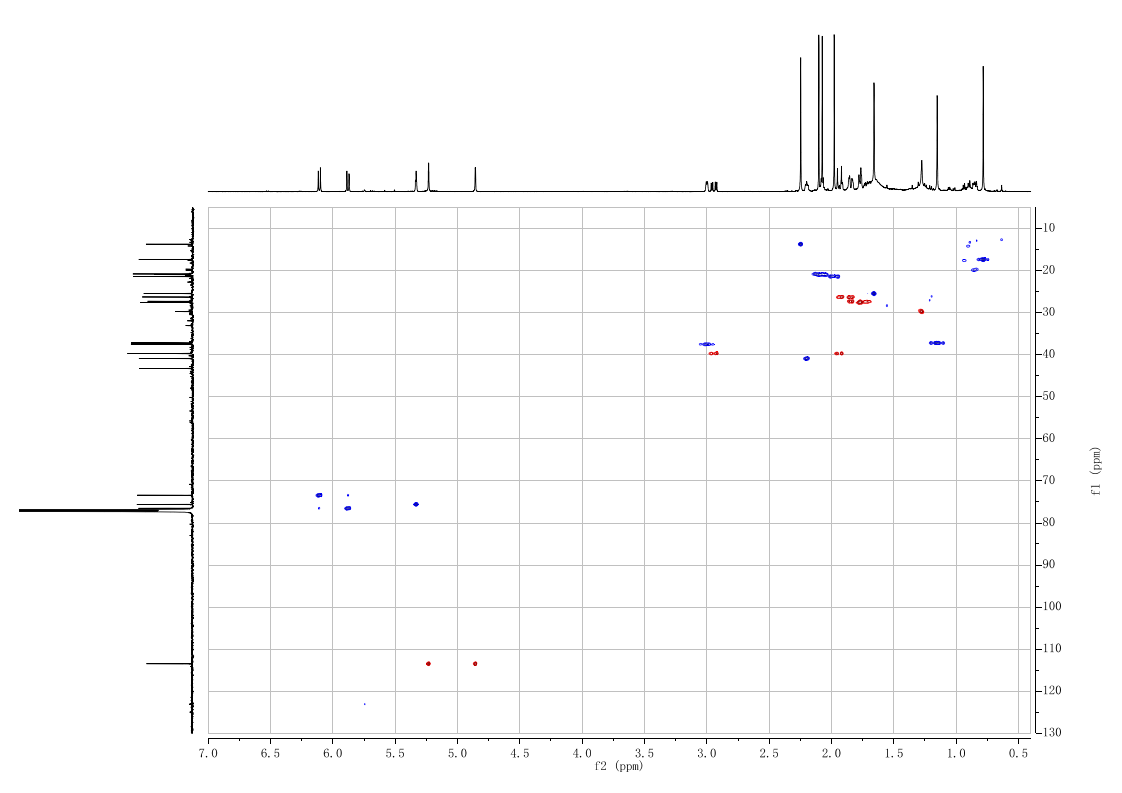

**Figure S25. ^1^H-^13^C HSQC spectrum of taxusin-13-one.**

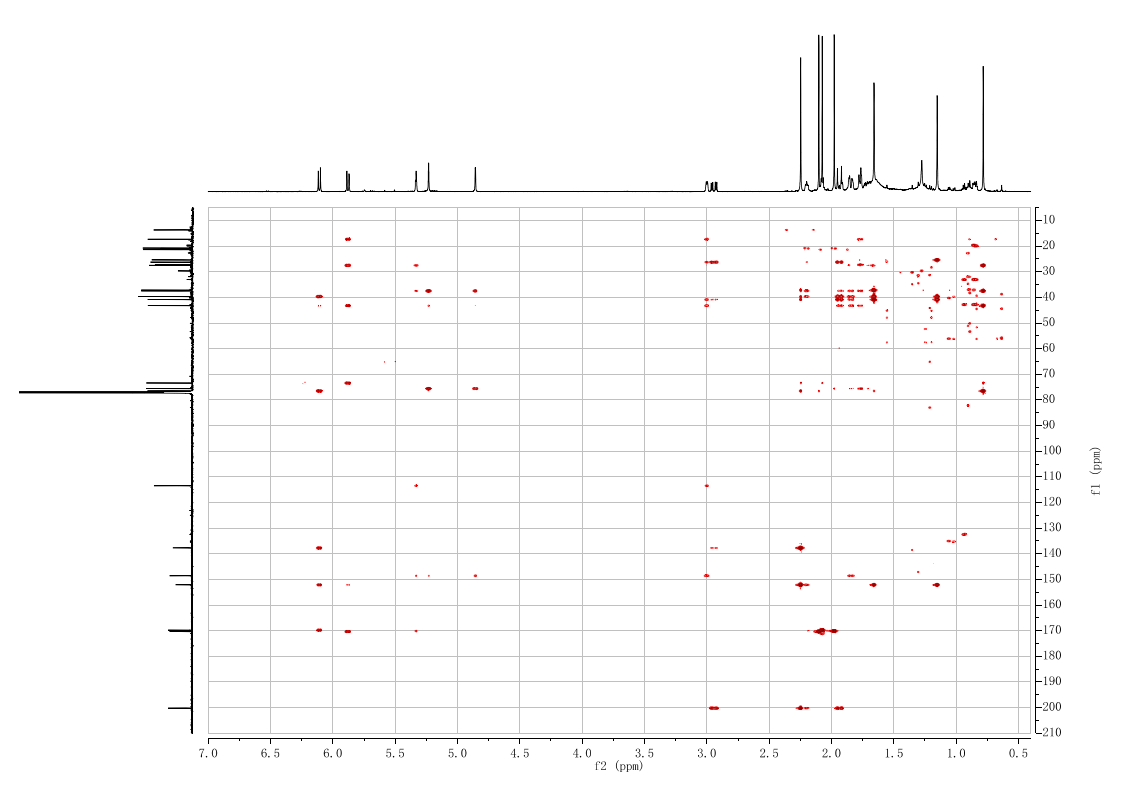

**Figure S26. ^1^H-^13^C HMBC spectrum of taxusin-13-one.**

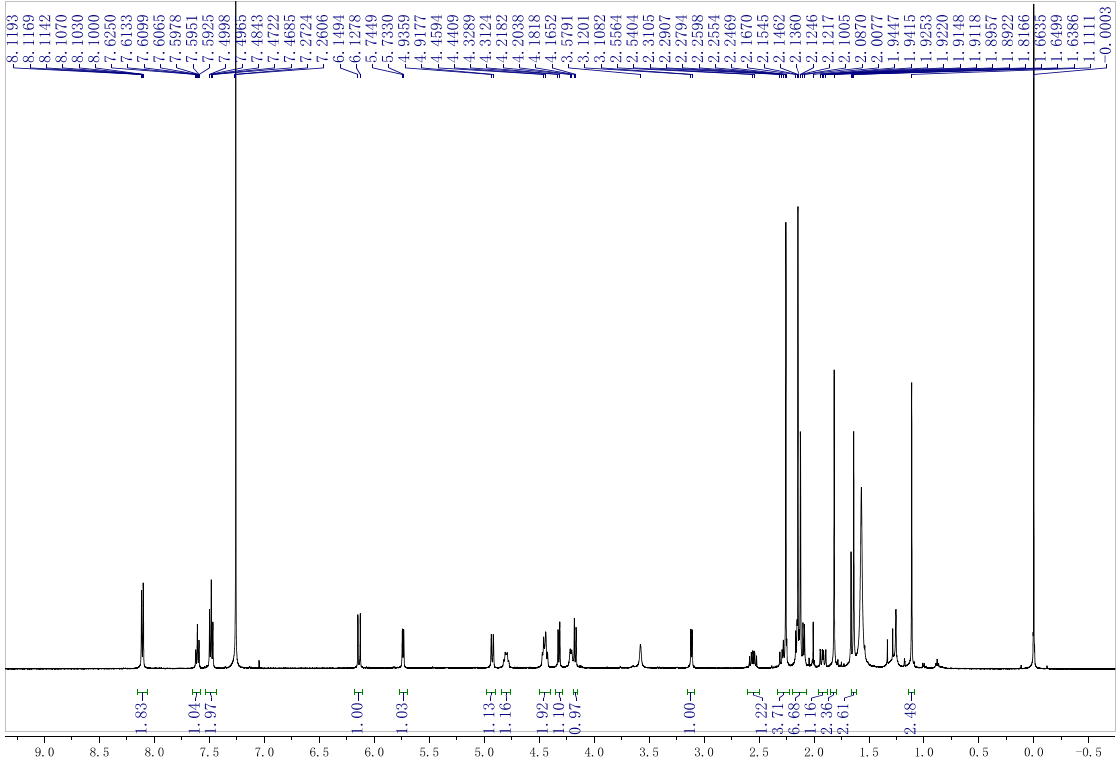

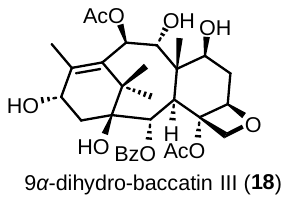

**Figure S27. ^1^H spectrum of 9*α*-hydroxyl-baccatin III (18).**

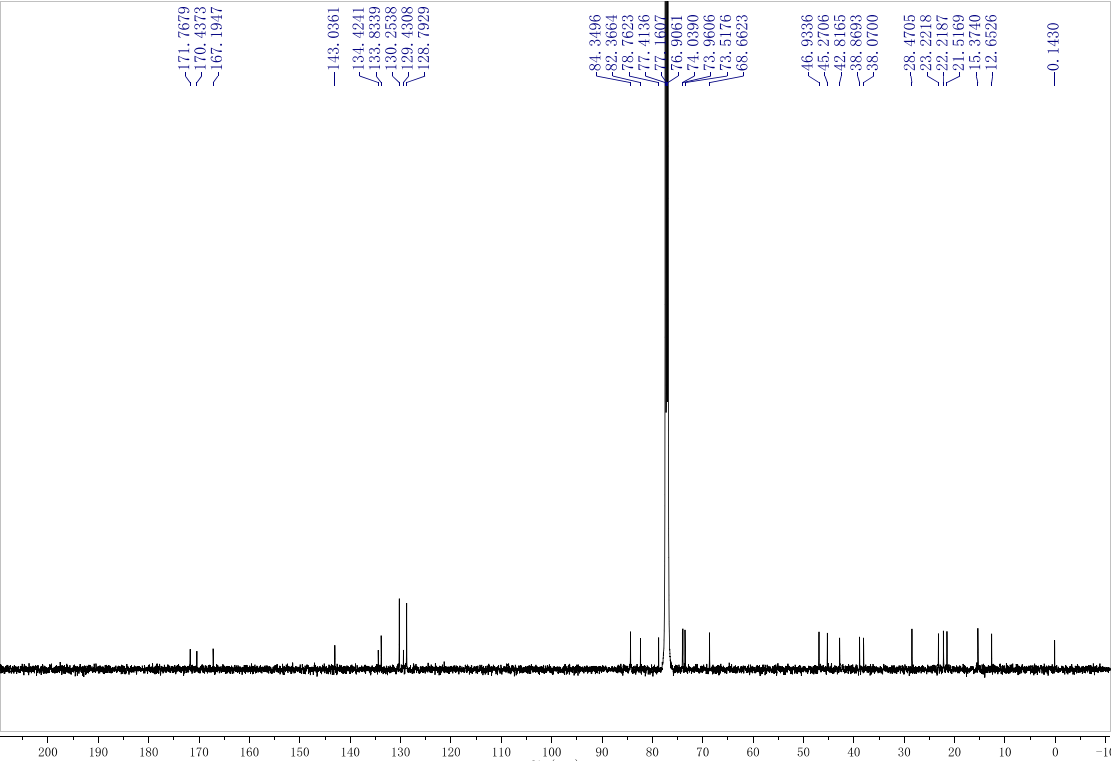

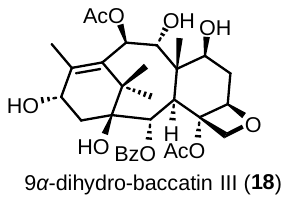

**Figure S28. ^13^C spectrum of 9*α*-dihydro-baccatin III (18).**

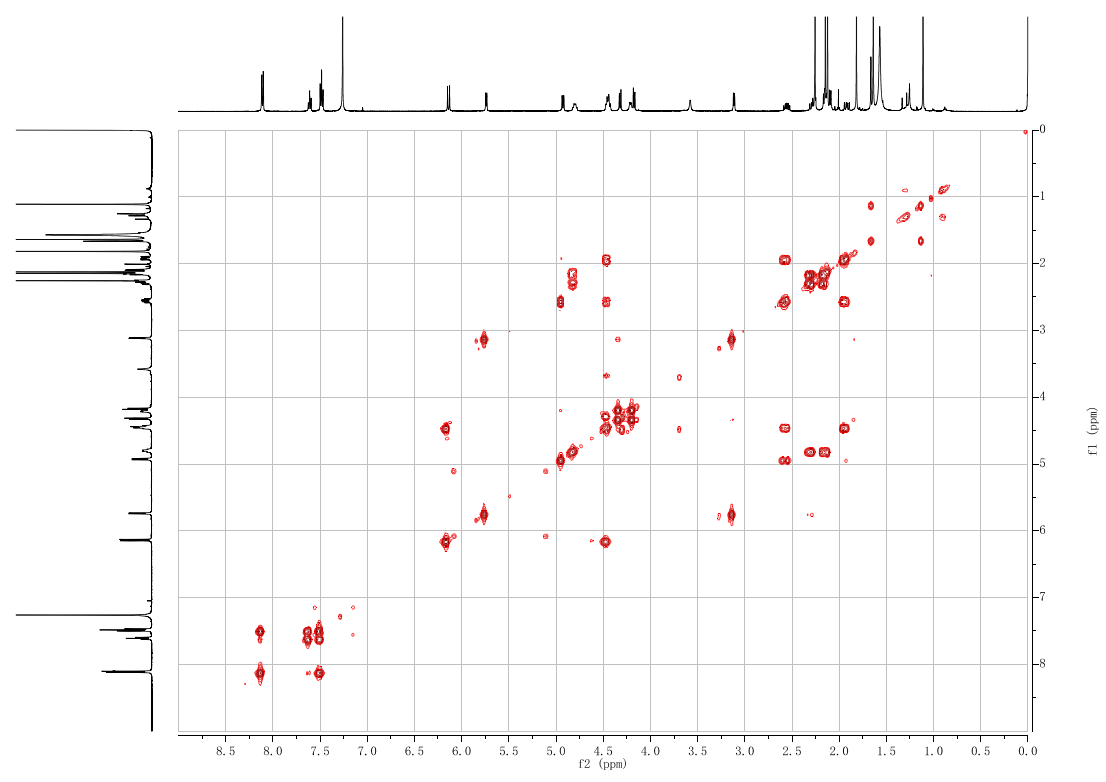

**Figure S29. ^1^H-^1^H COSY spectrum of 9*α*-dihydro-baccatin III (18).**

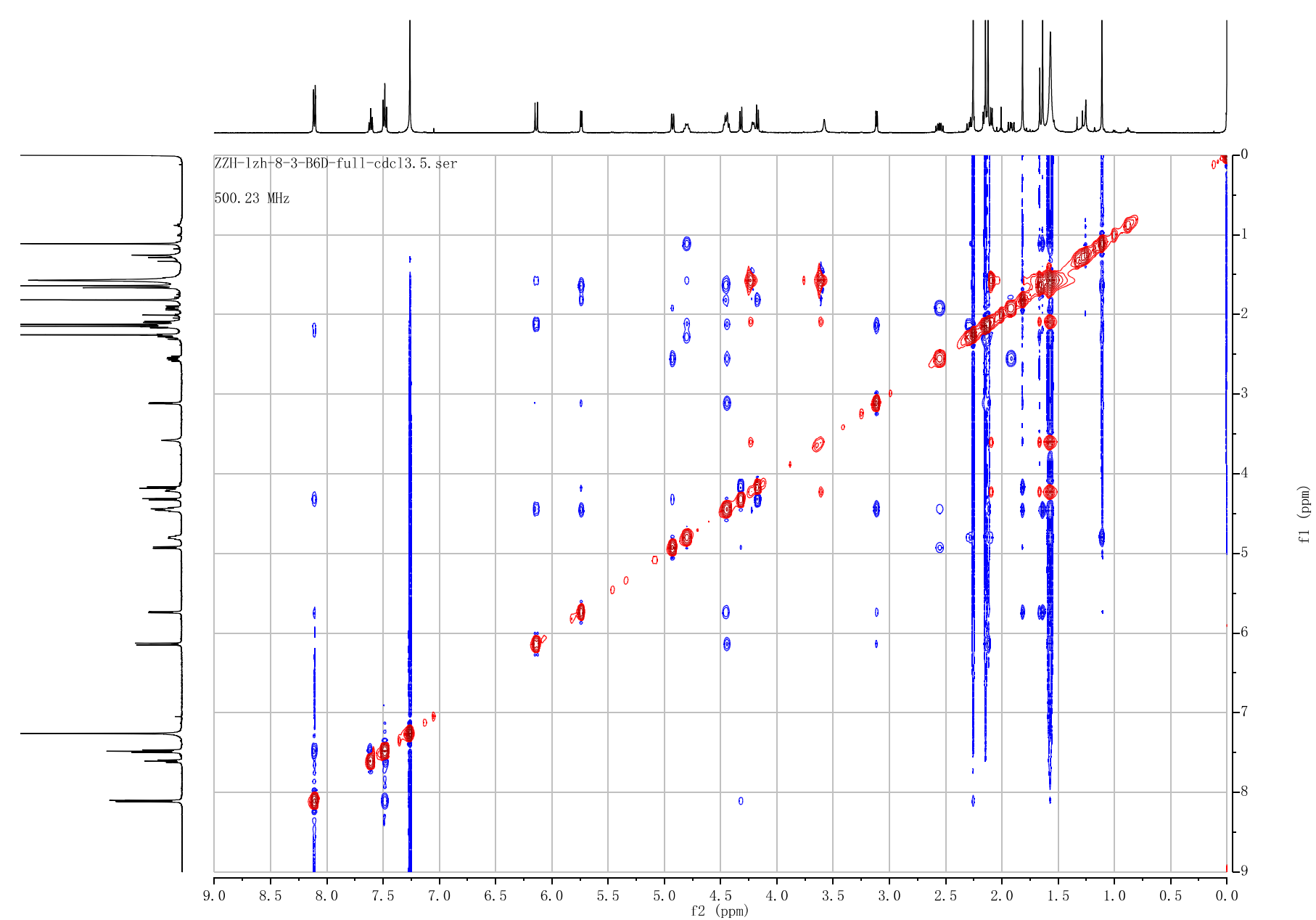

**Figure S30. ^1^H-^1^H NOE spectrum of 9*α*-dihydro-baccatin III (18).**

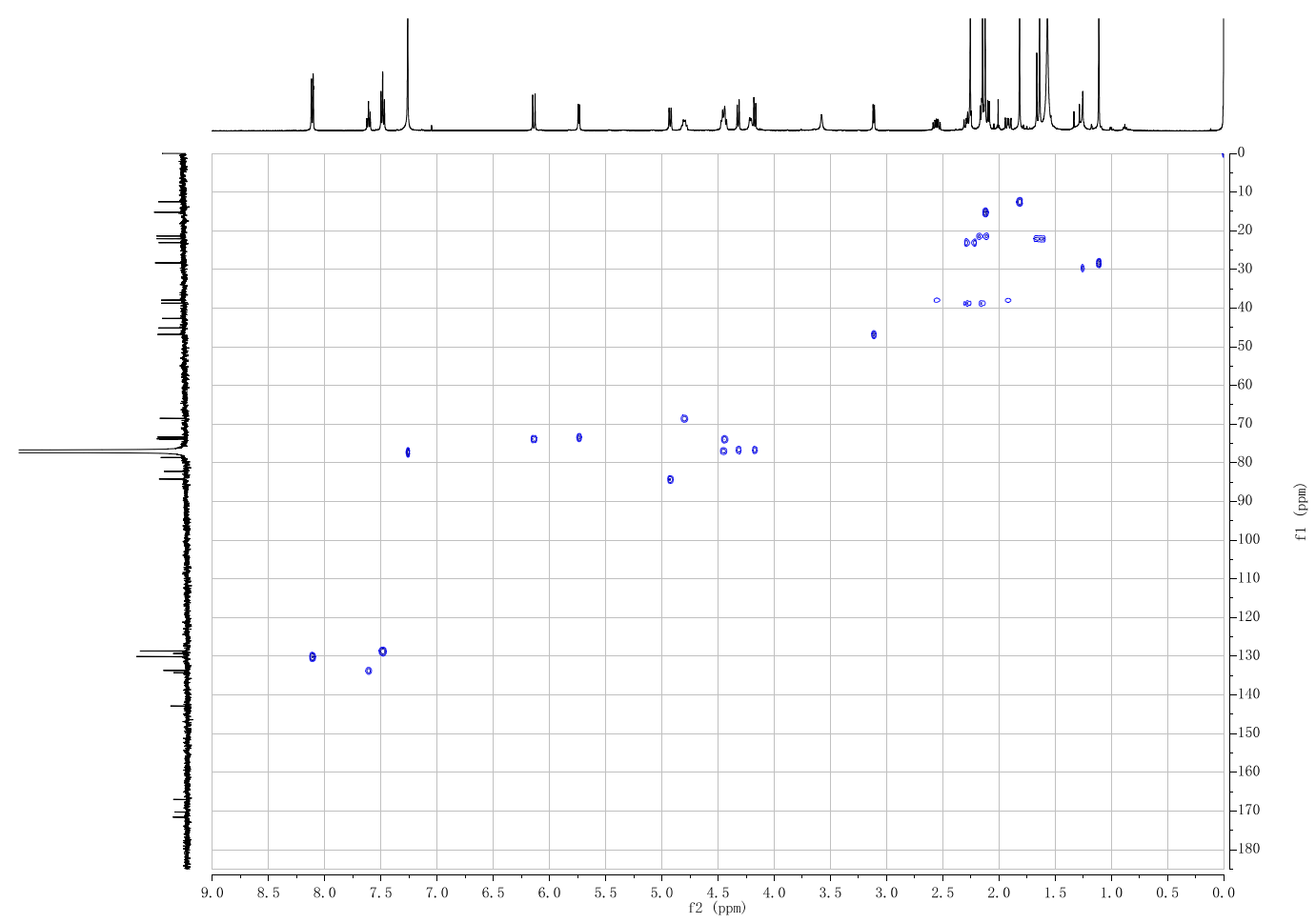

**Figure S31. ^1^H-^13^C HSQC spectrum of 9*α*-dihydro-baccatin III (18).**

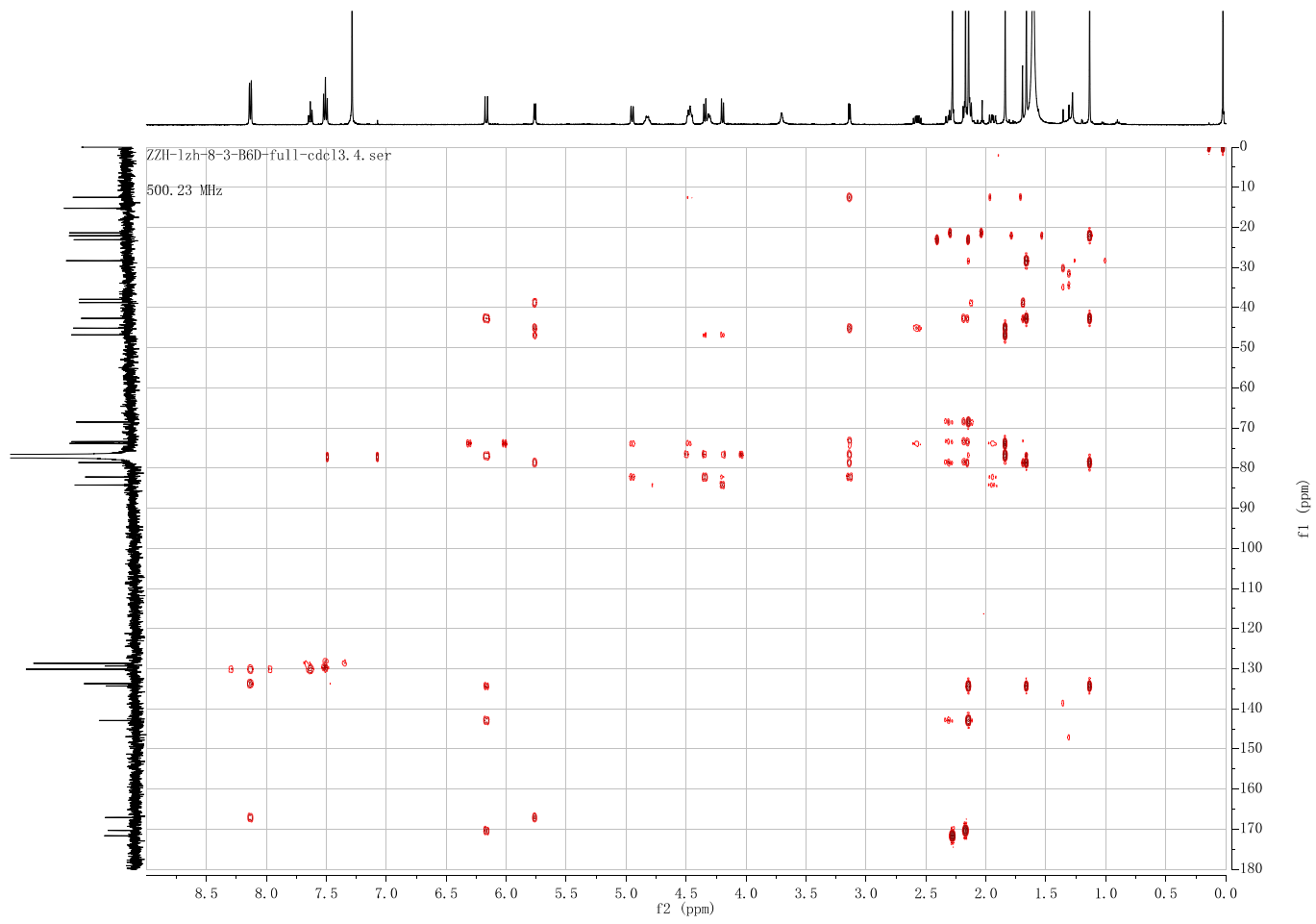

**Figure S32. ^1^H-^13^C HMBC spectrum of 9*α*-dihydro-baccatin III (18).**

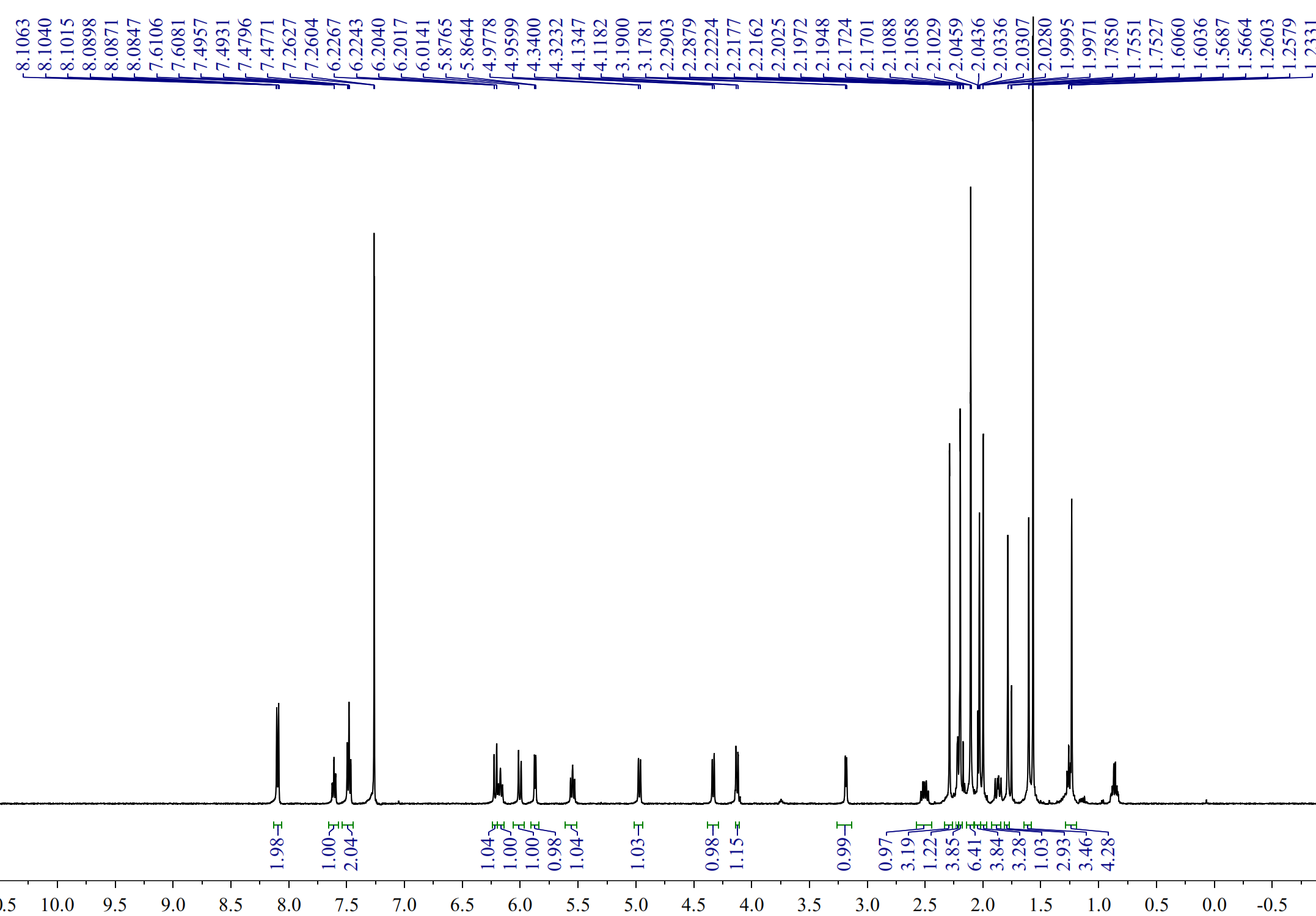

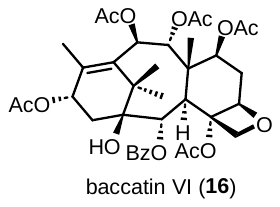

**Figure S33. ^1^H NMR spectrum of baccatin VI (16).**

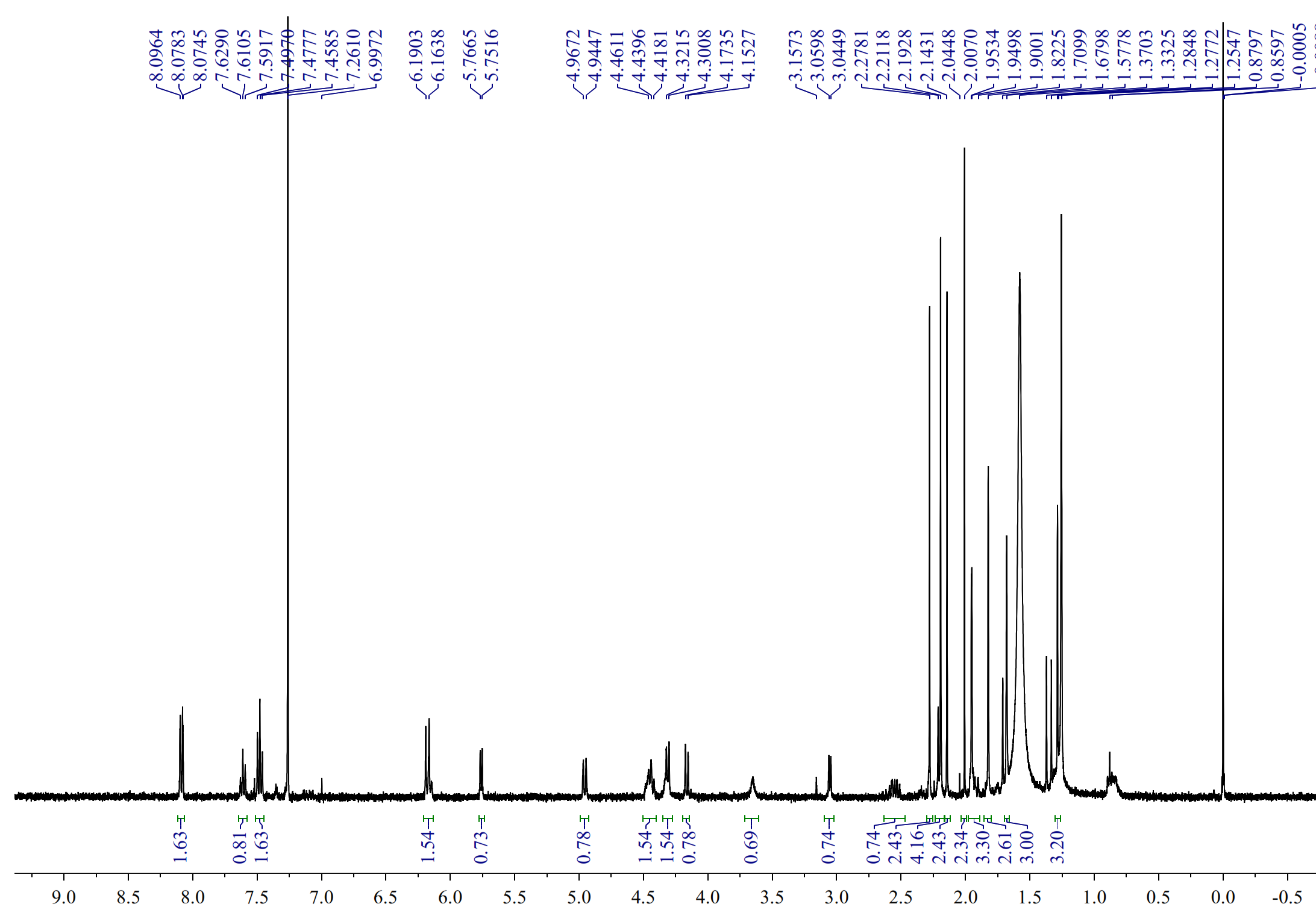

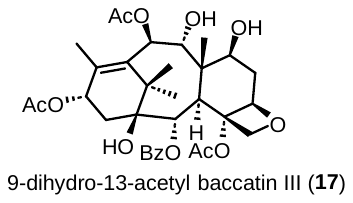

**Figure S34. ^1^H NMR spectrum of 9-dihydro-13-acetyl baccatin III (17).**

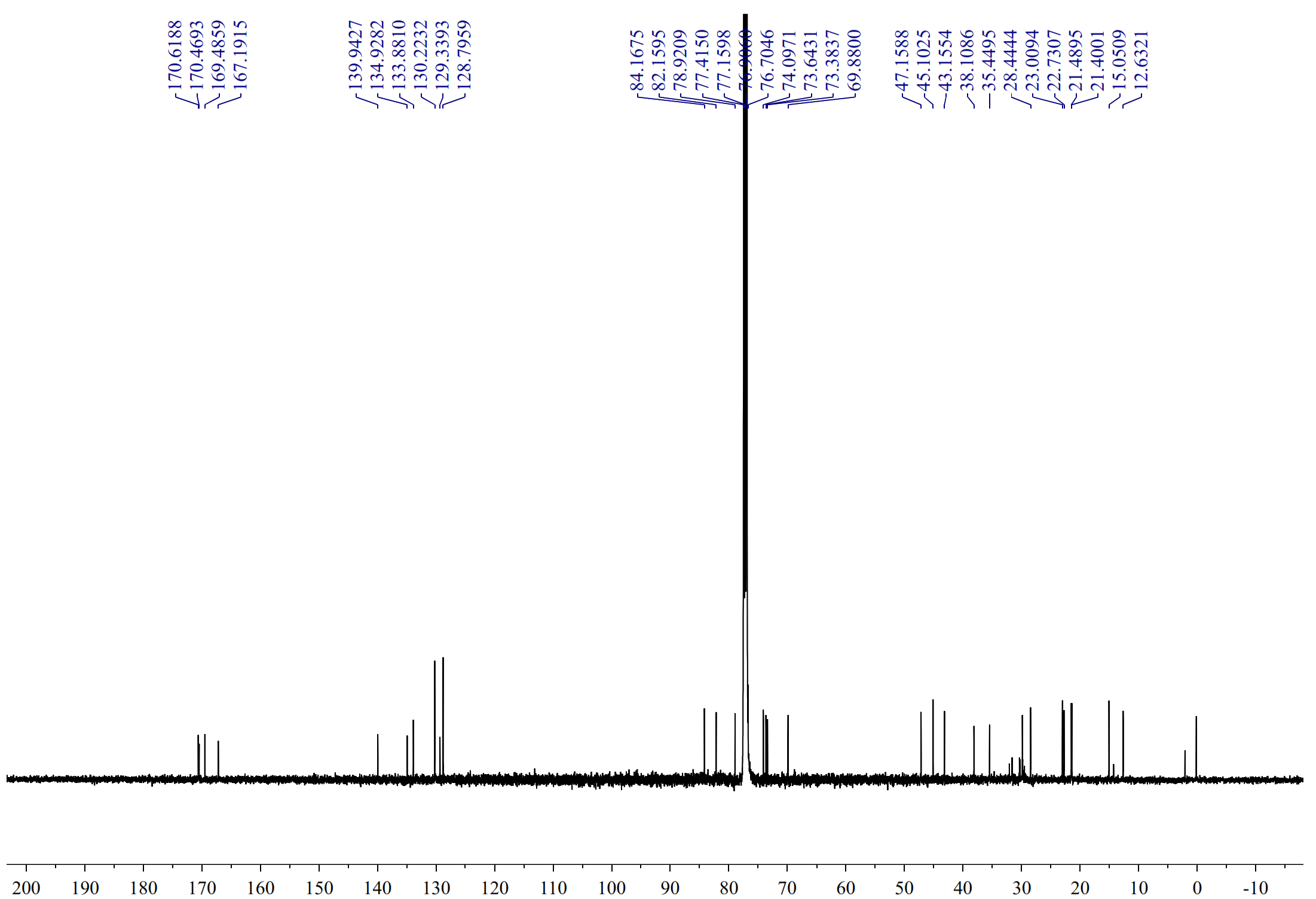

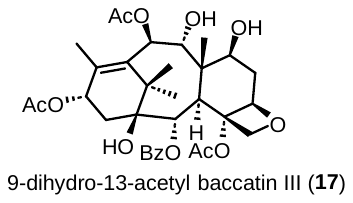

**Figure S35: ^13^C NMR spectrum of 9-dihydro-13-acetyl baccatin III (17).**

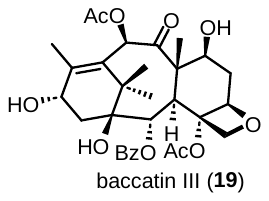

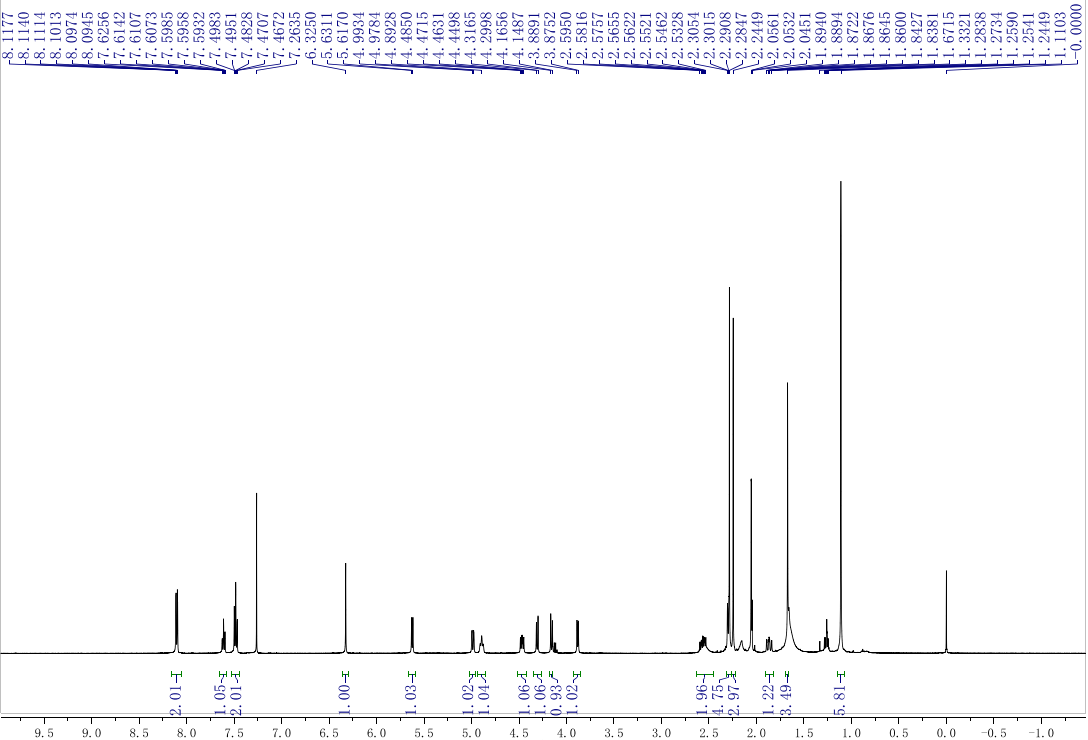

**Figure S36. ^1^H spectrum of biosynthesized baccatin III (19).**

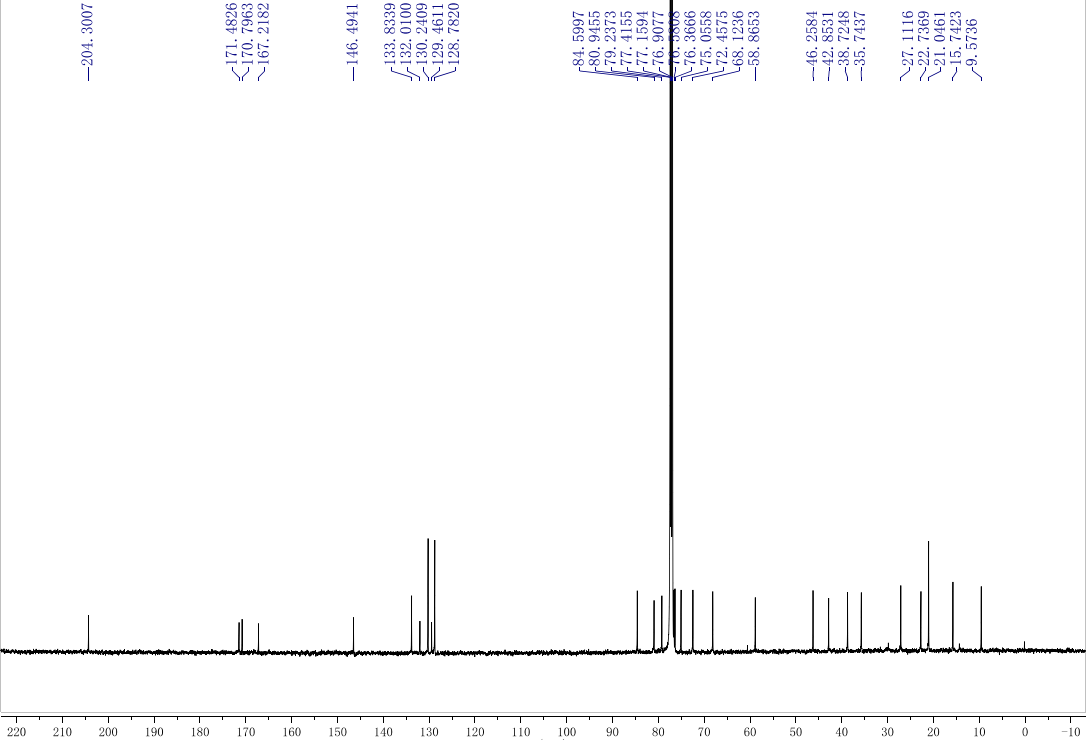

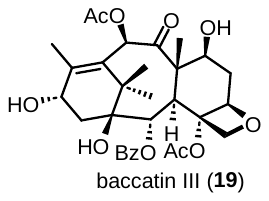

**Figure S37. ^13^C spectrum of** **biosynthesized baccatin III (19).**

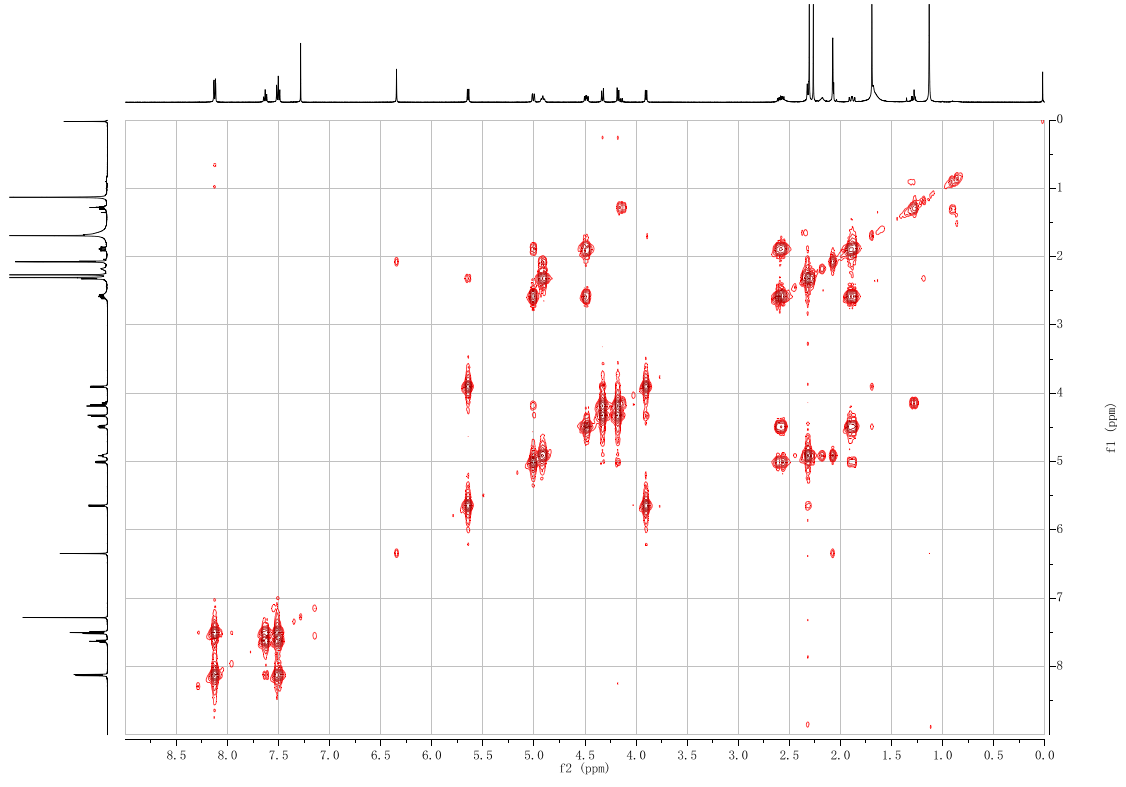

**Figure S38. ^1^H-^1^H COSY spectrum of baccatin III (19)**

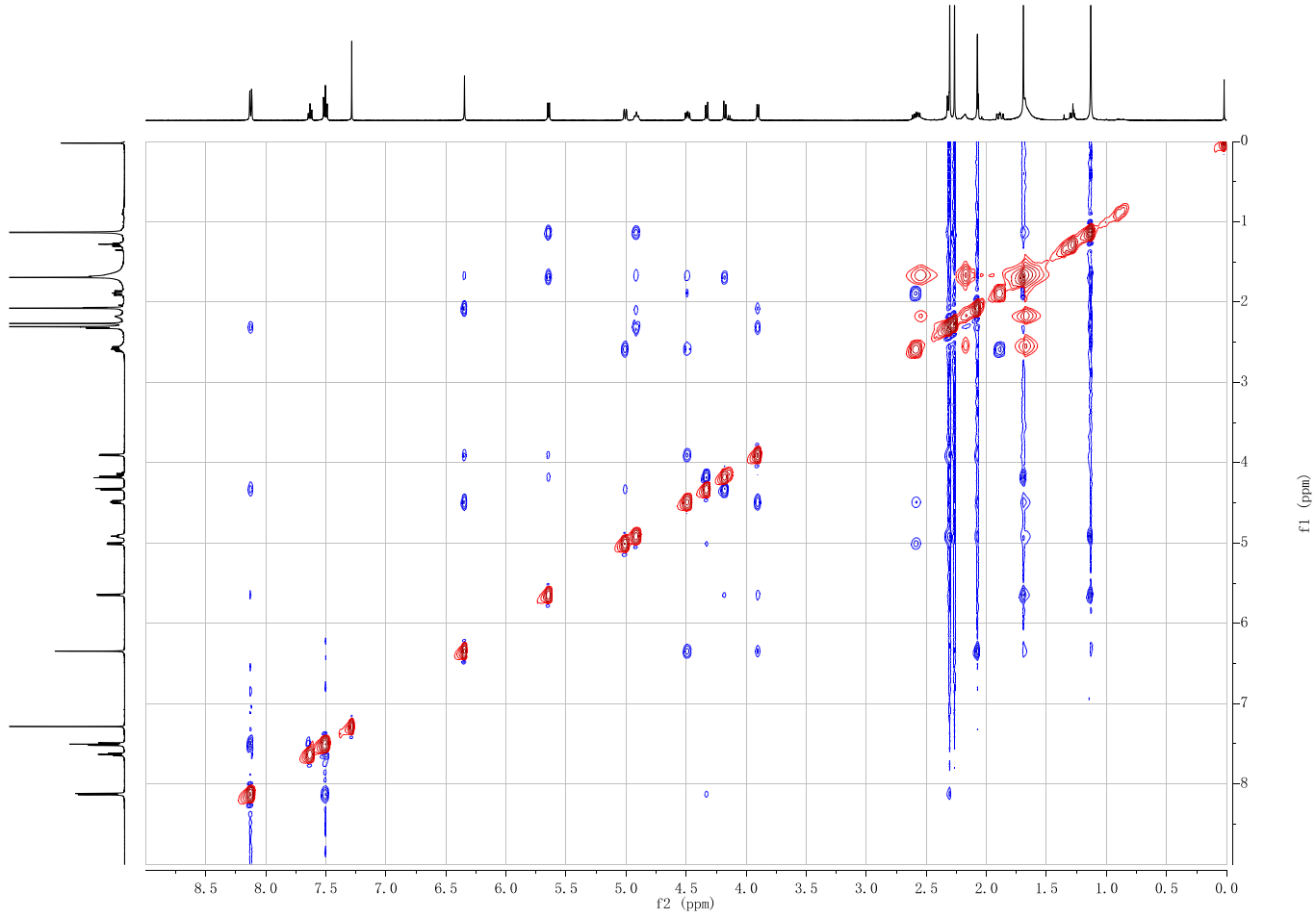

**Figure S39. ^1^H-^1^H NOE spectrum of baccatin III (19).**

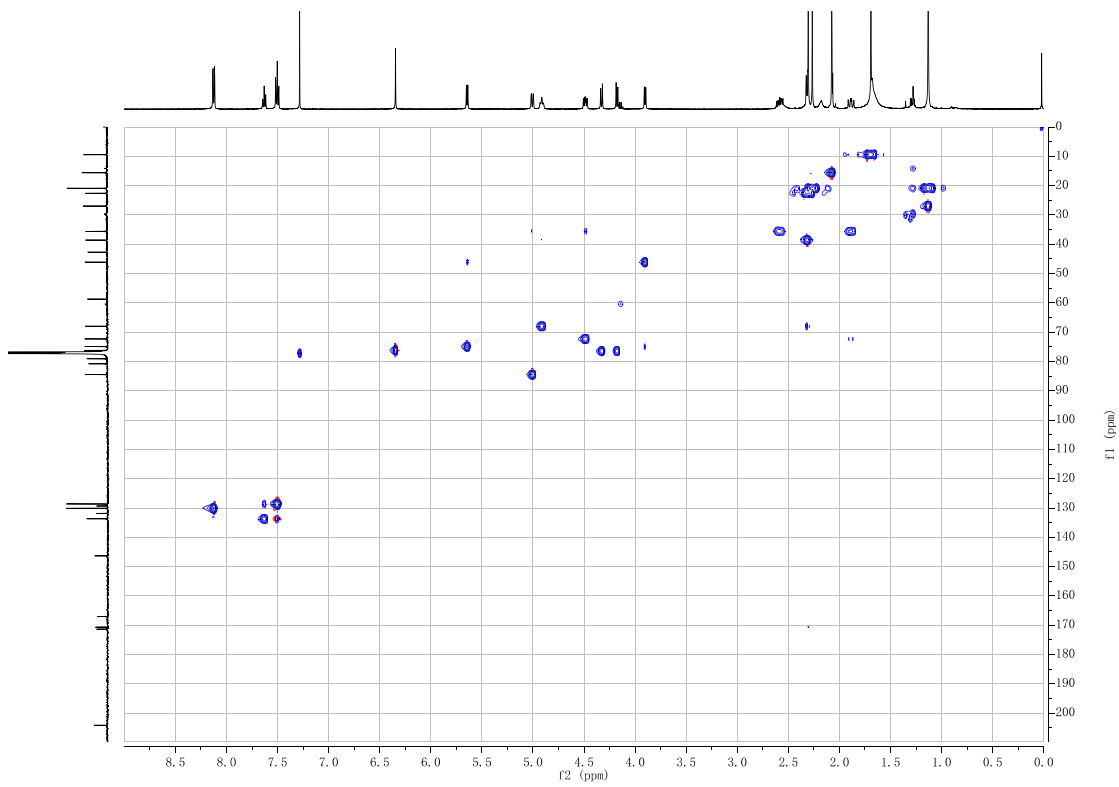

**Figure S40. ^1^H-^13^C HSQC spectrum of baccatin III (19).**

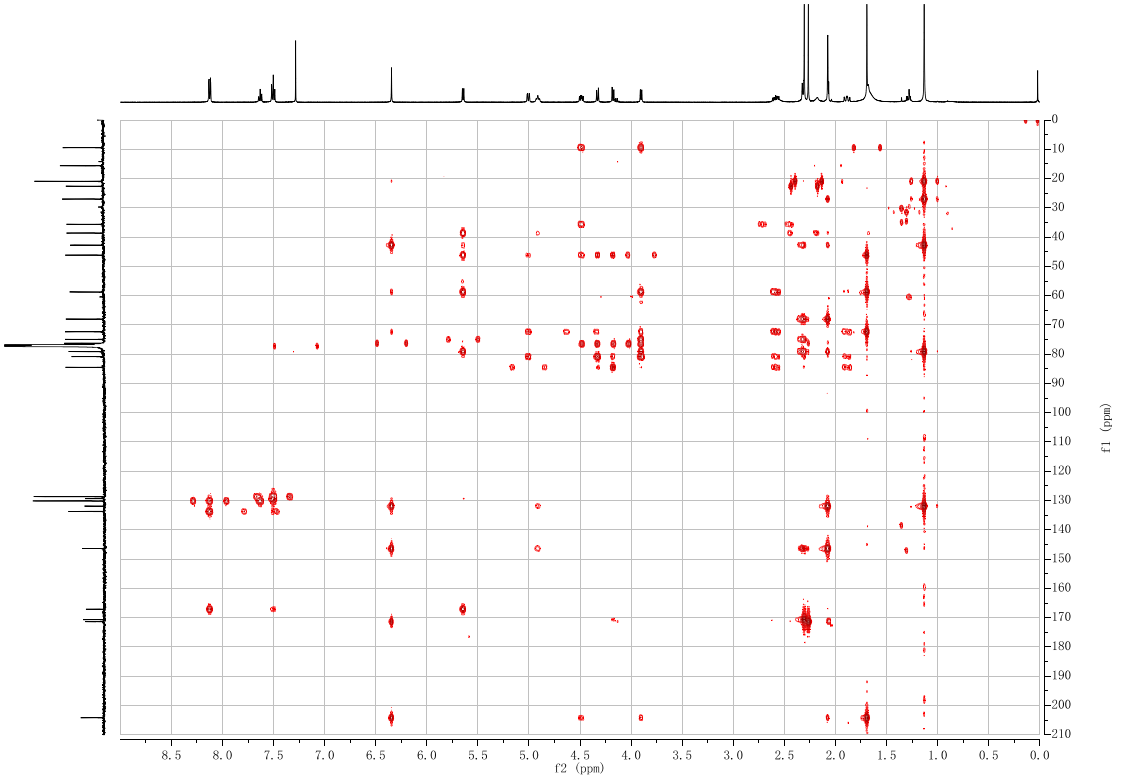

**Figure S41. ^1^H-^13^C HMBC spectrum of baccatin III (19)**

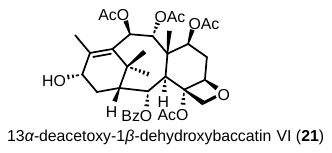

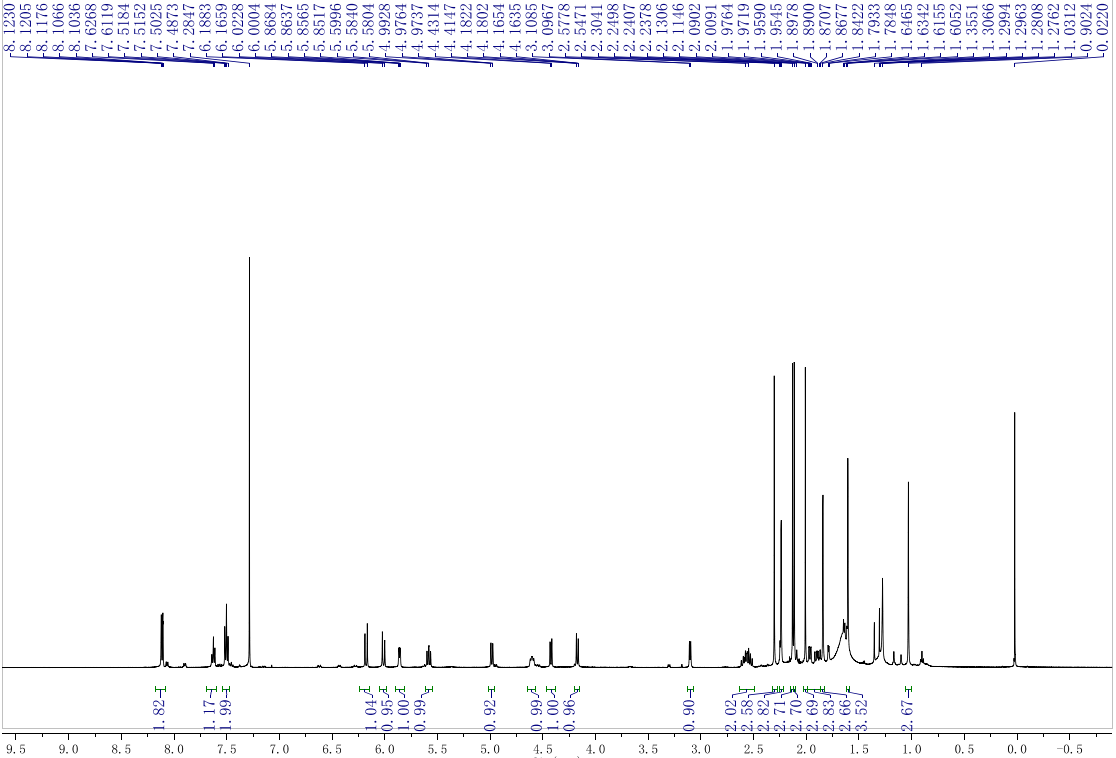

**Figure S42. ^1^H spectrum of 13*α*-deacetoxy-1*β*-dehydroxybaccatin VI (21)**

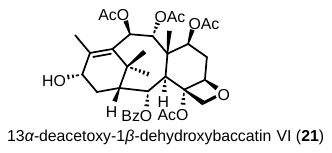

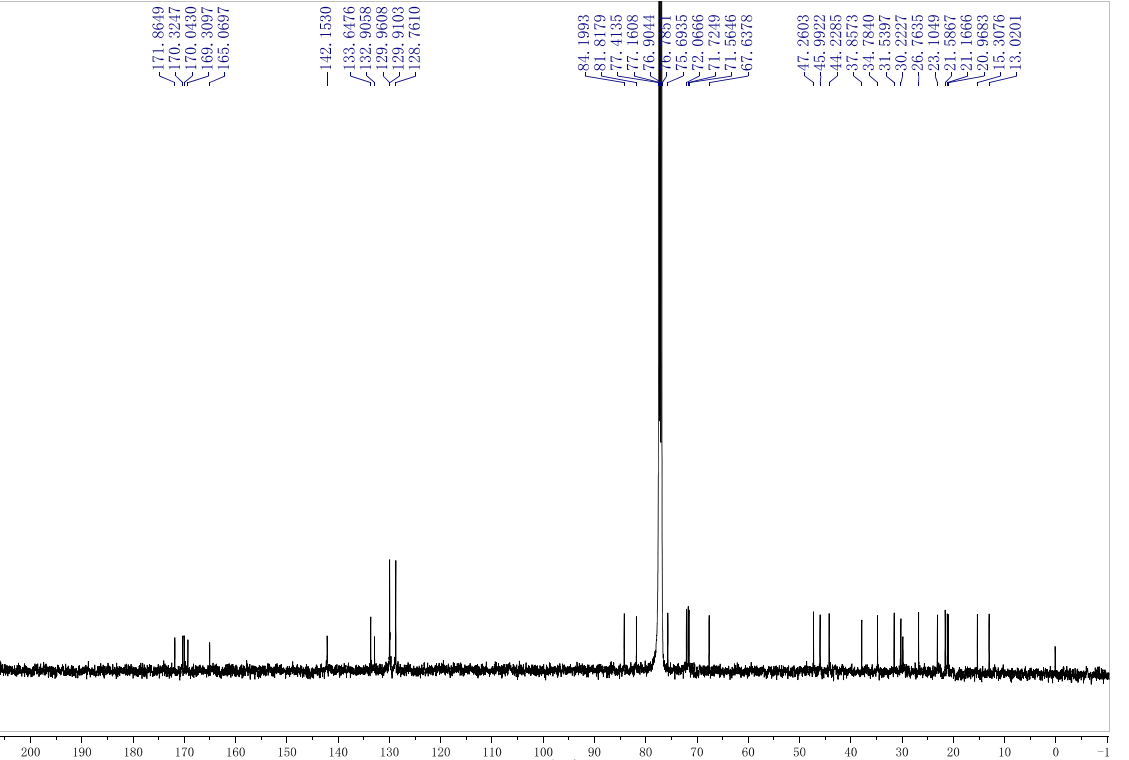

**Figure S43. ^13^C spectrum of 13*α*-deacetoxy-1*β*-dehydroxybaccatin VI (21).**

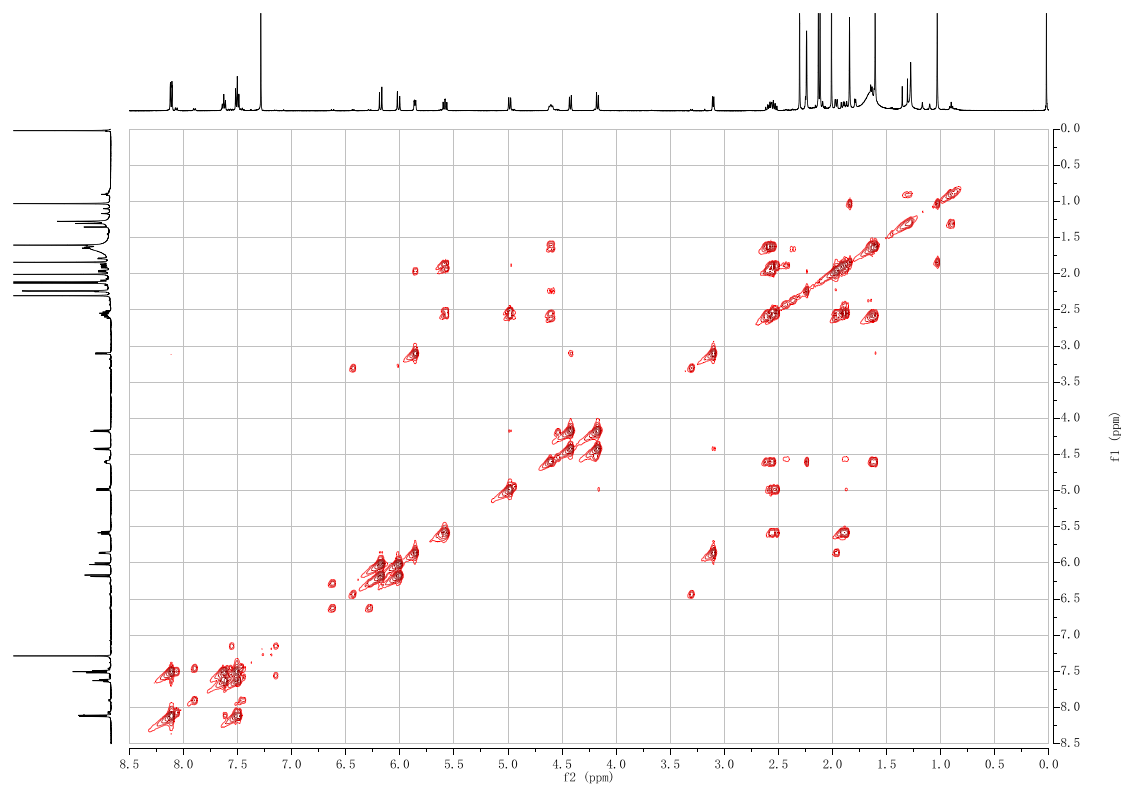

**Figure S44. ^1^H-^1^H COSY spectrum of 13*α*-deacetoxy-1*β*-dehydroxybaccatin VI (21).**

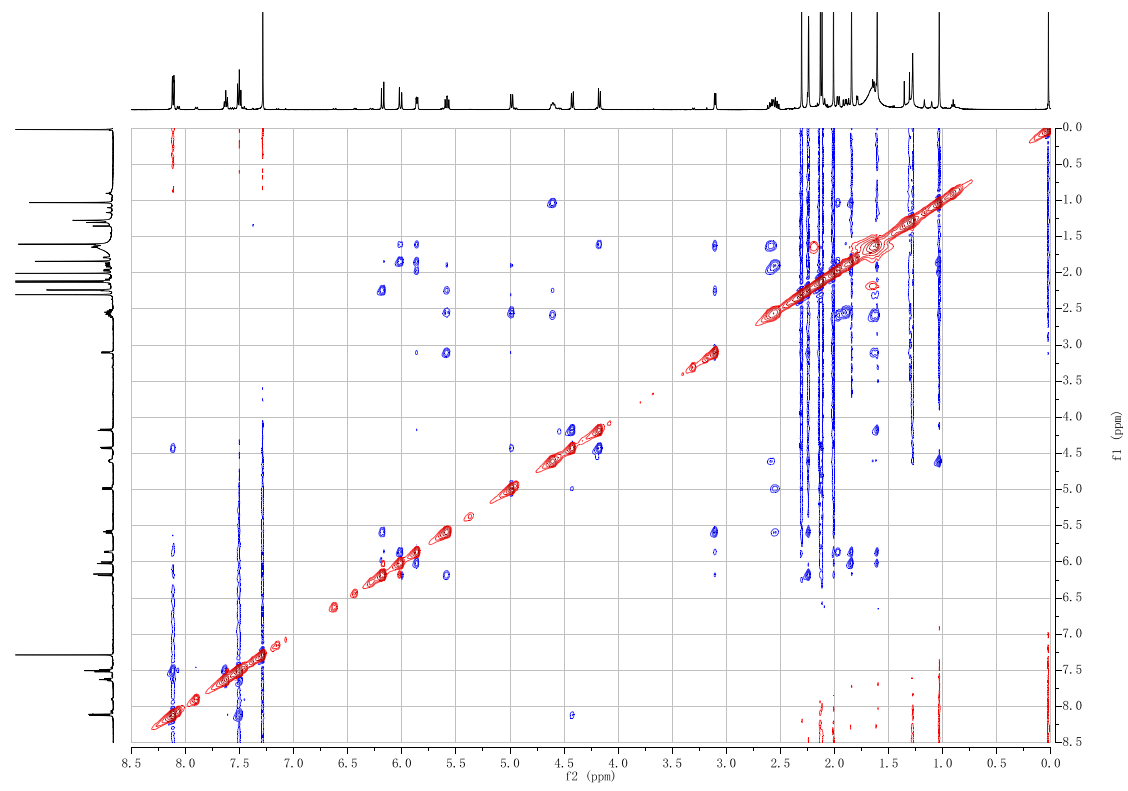

**Figure S45. ^1^H-^1^H NOE spectrum of 13*α*-deacetoxy-1*β*-dehydroxybaccatin VI (21).**

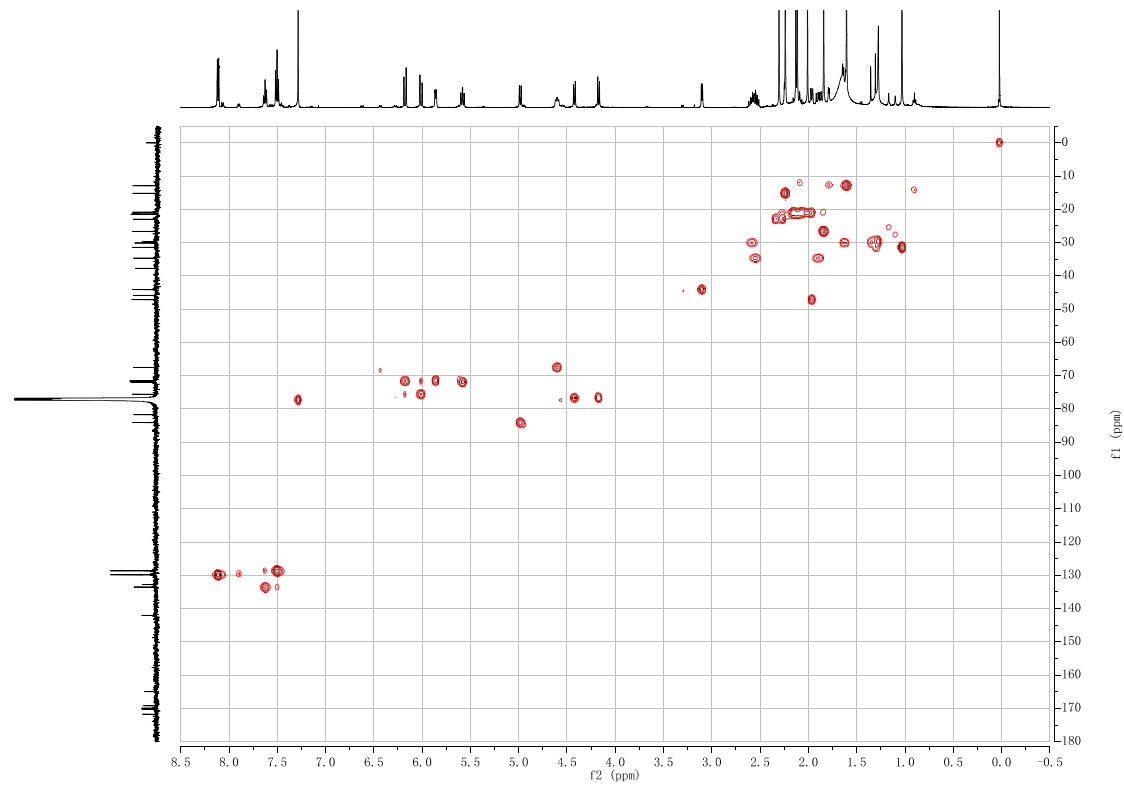

**Figure S46. ^1^H-^13^C HSQC spectrum of 13*α*-deacetoxy-1*β*-dehydroxybaccatin VI (21).**

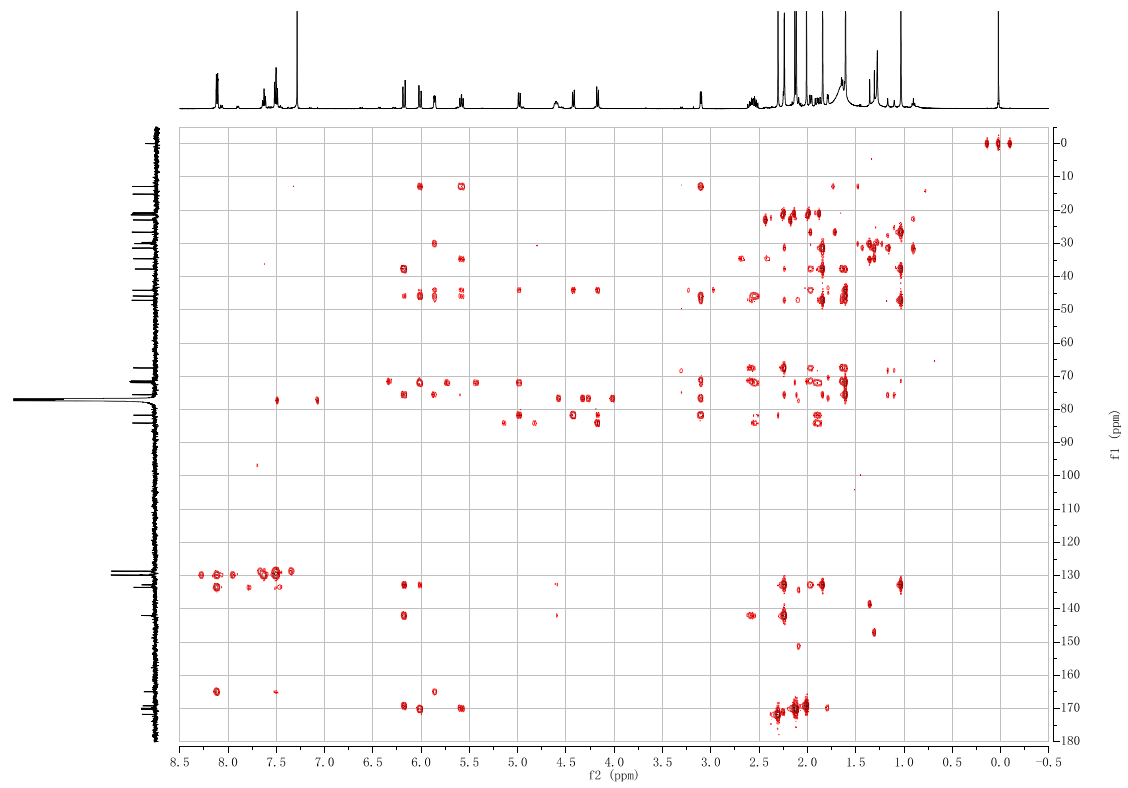

**Figure S47. ^1^H-^13^C HMBC spectrum of 13*α*-deacetoxy-1*β*-dehydroxybaccatin VI (21).**

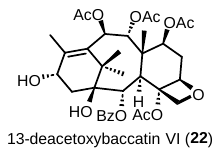

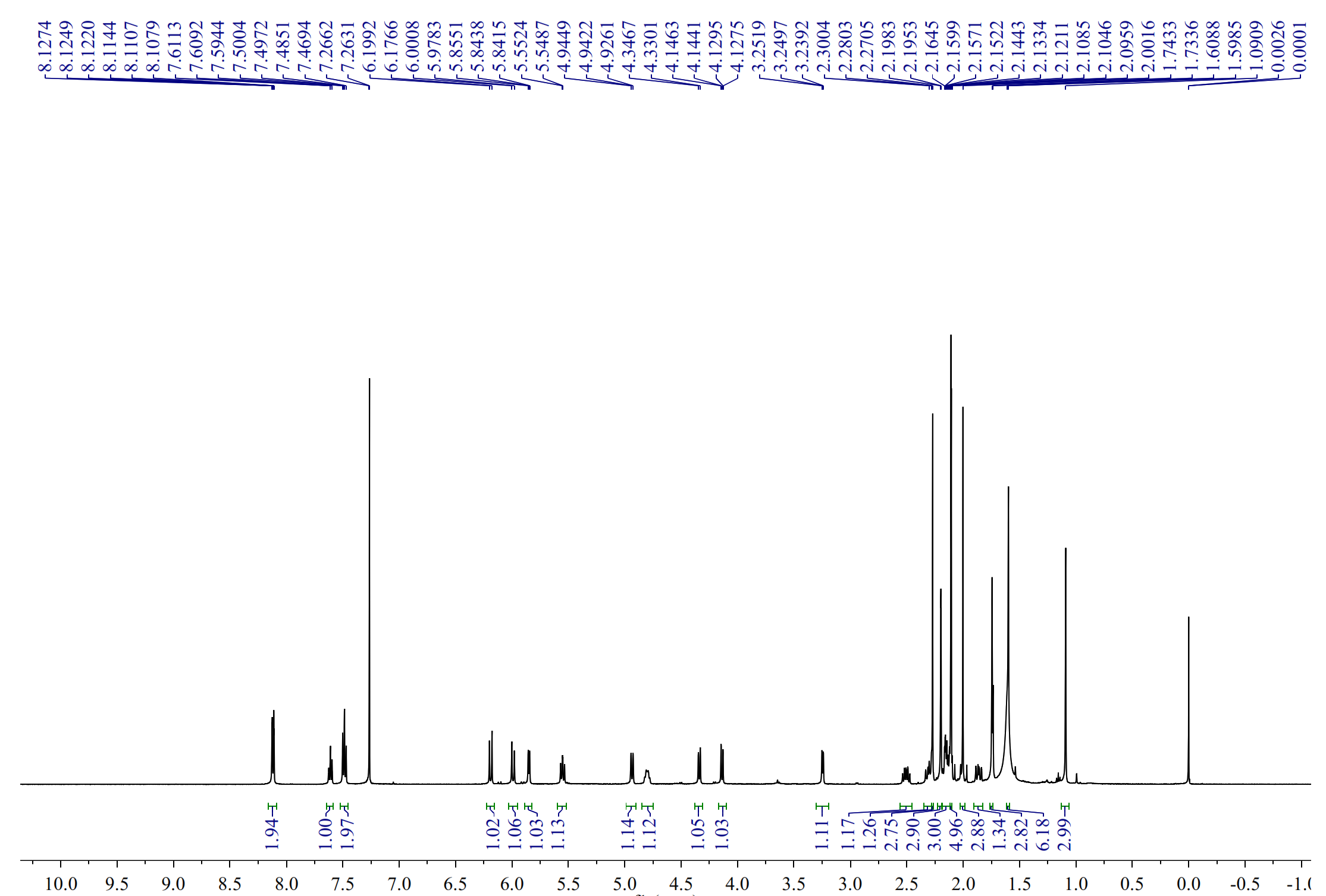

**Figure S48. ^1^H NMR spectrum of 13-deacetoxybaccatin VI (22).**

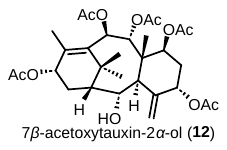

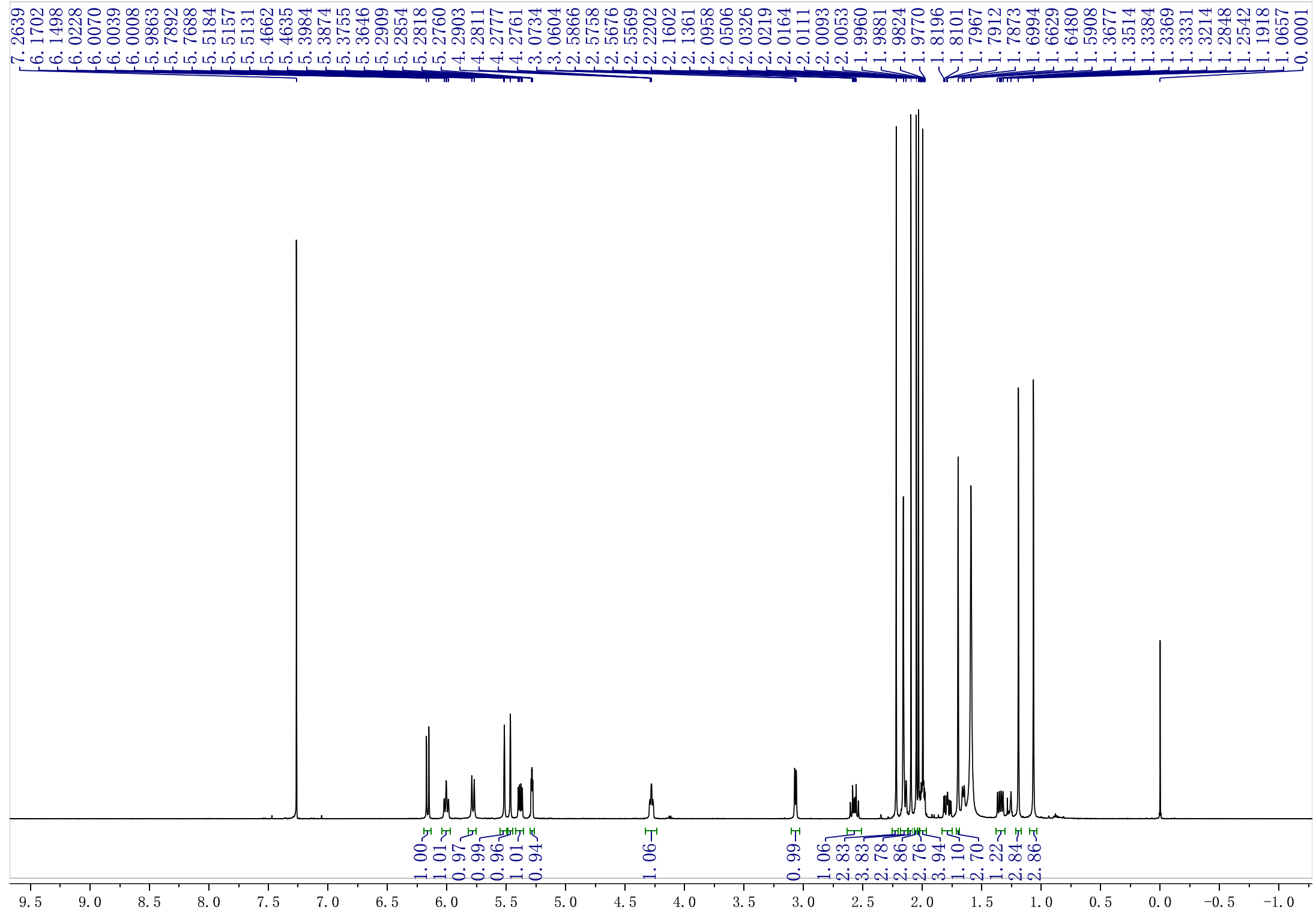

**Figure S49. ^1^H NMR spectrum of 7*β*-acetoxytauxin-2*α*-ol (12)**

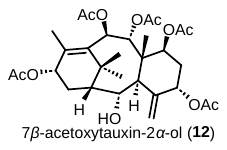

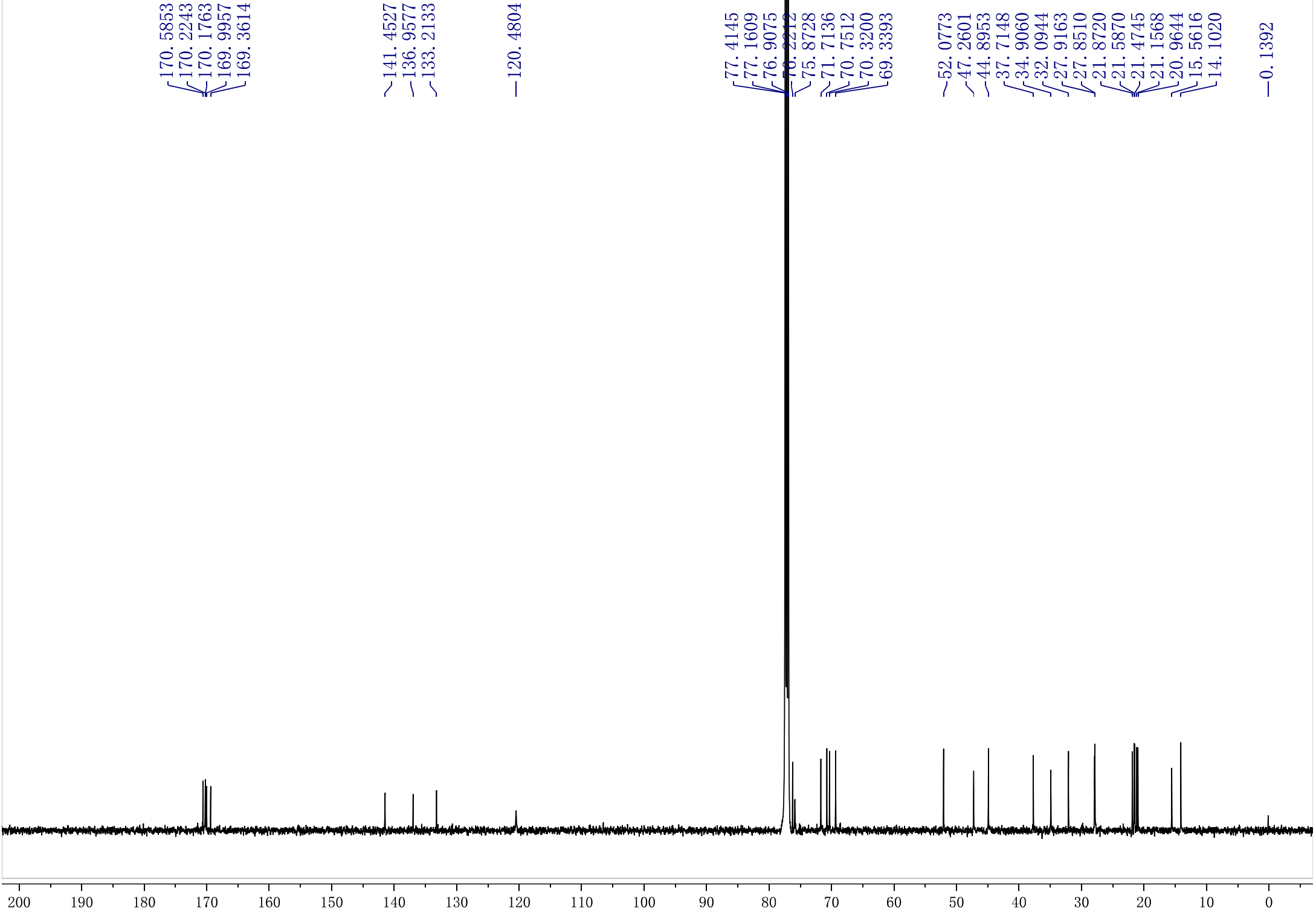

**Figure S50. ^13^C NMR spectrum of 7*β*-acetoxytauxin-2*α*-ol (12)**

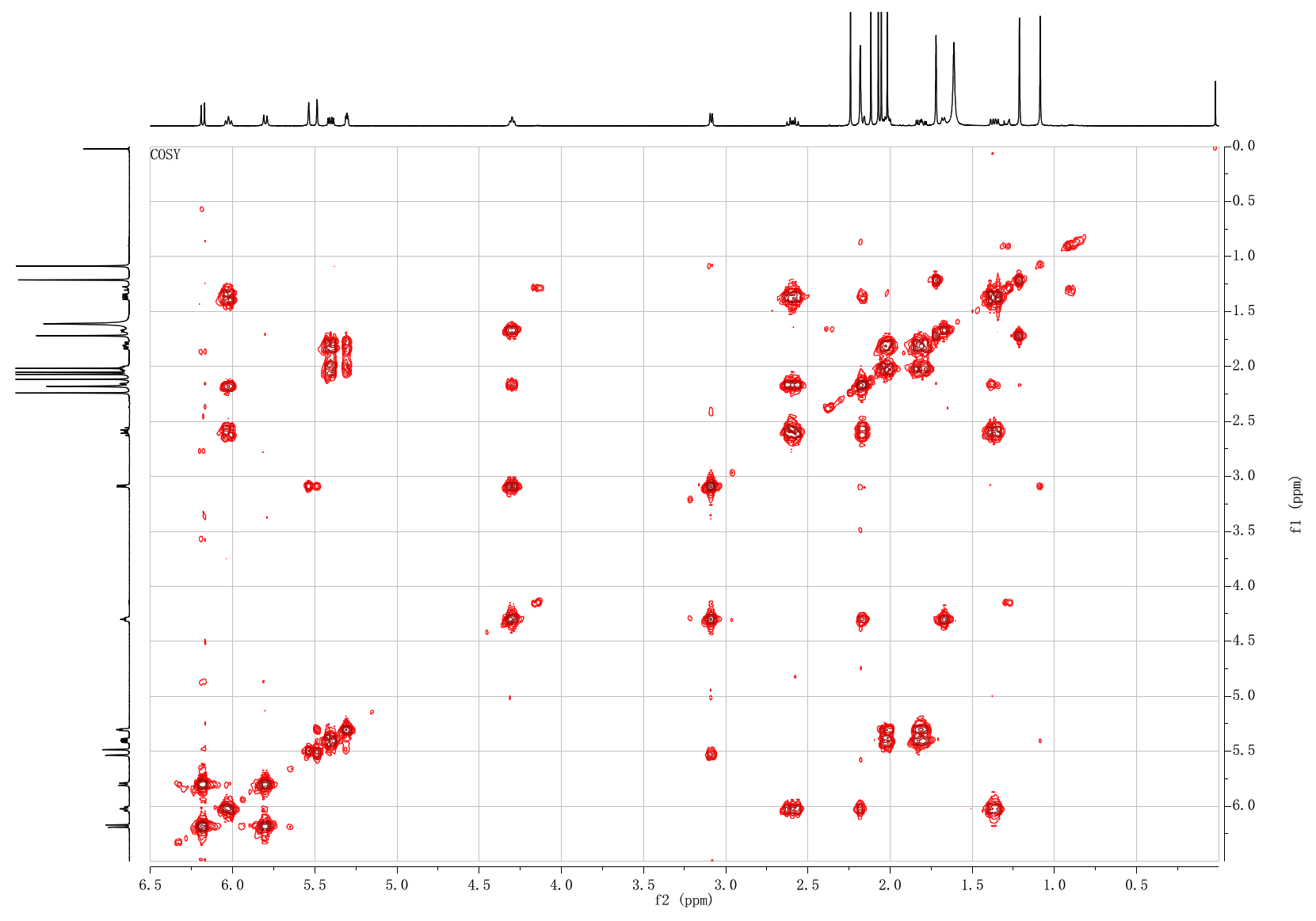

**Figure S51. ^1^H-^1^H COSY NMR spectrum of 7*β*-acetoxytauxin-2*α*-ol (12)**

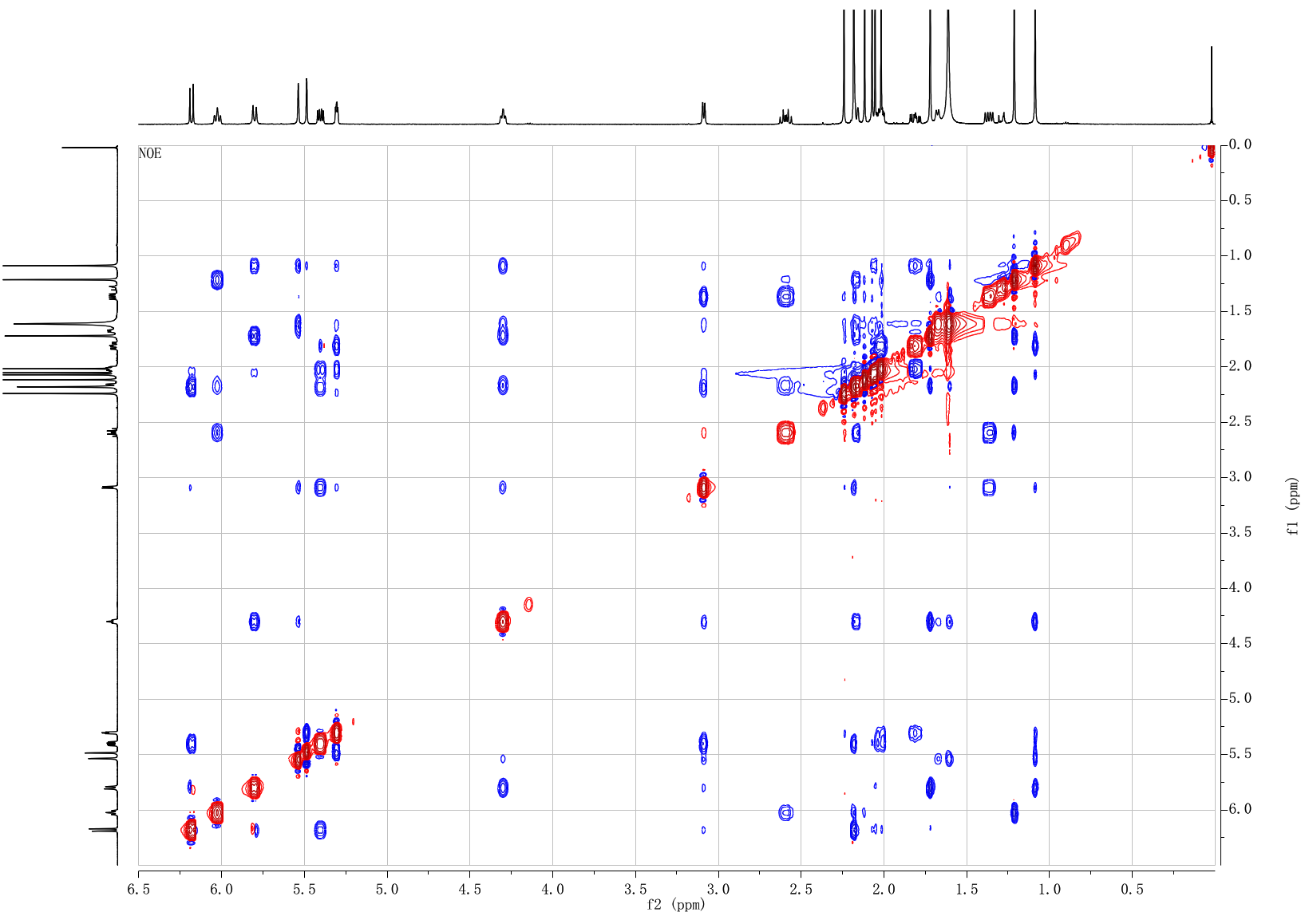

**Figure S52. ^1^H-^1^H NOESY NMR spectrum of 7*β*-acetoxytauxin-2*α*-ol (12).**

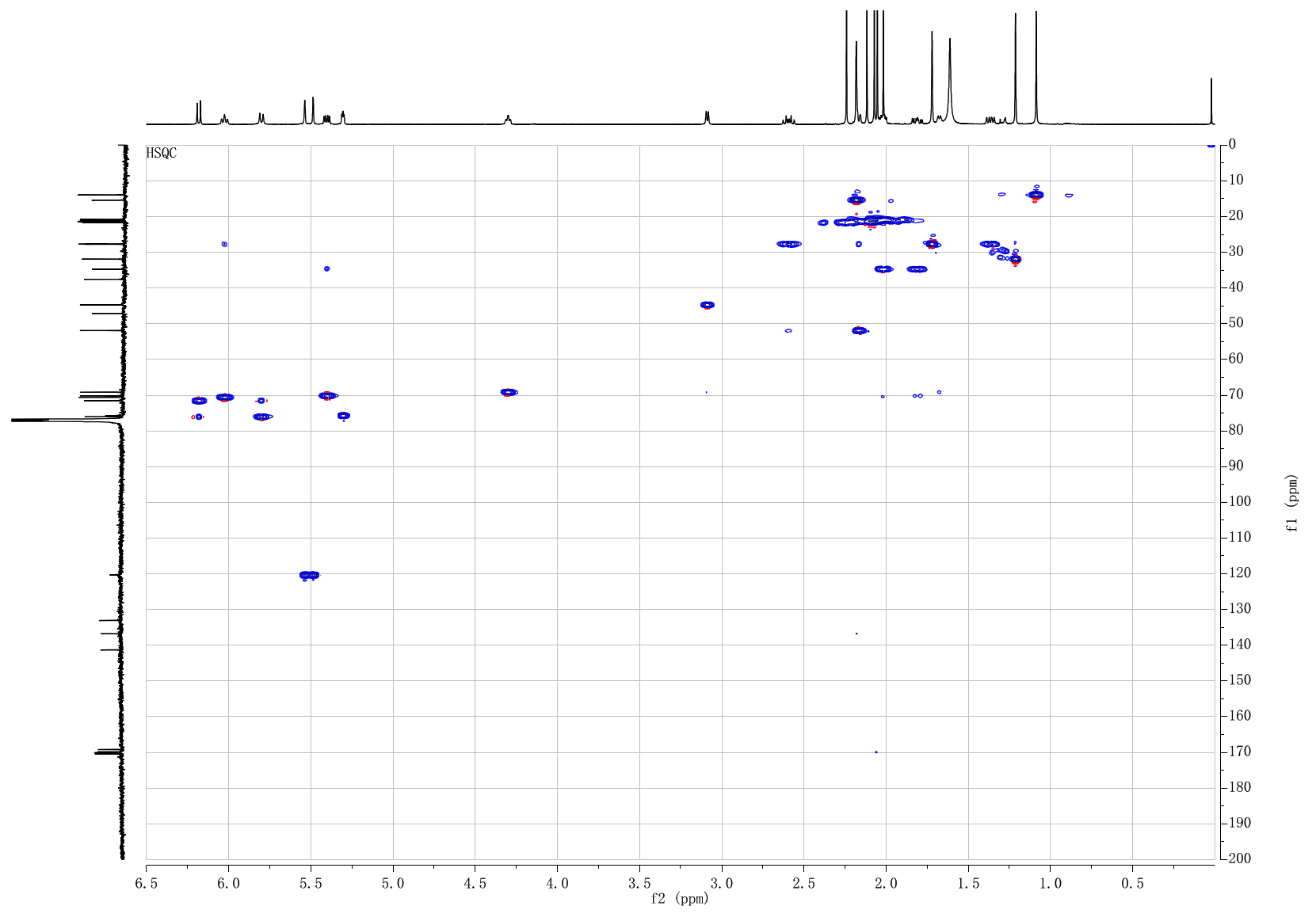

**Figure S53. ^1^H-^13^C HSQC NMR spectrum of 7*β*-acetoxytauxin-2*α*-ol (12)**

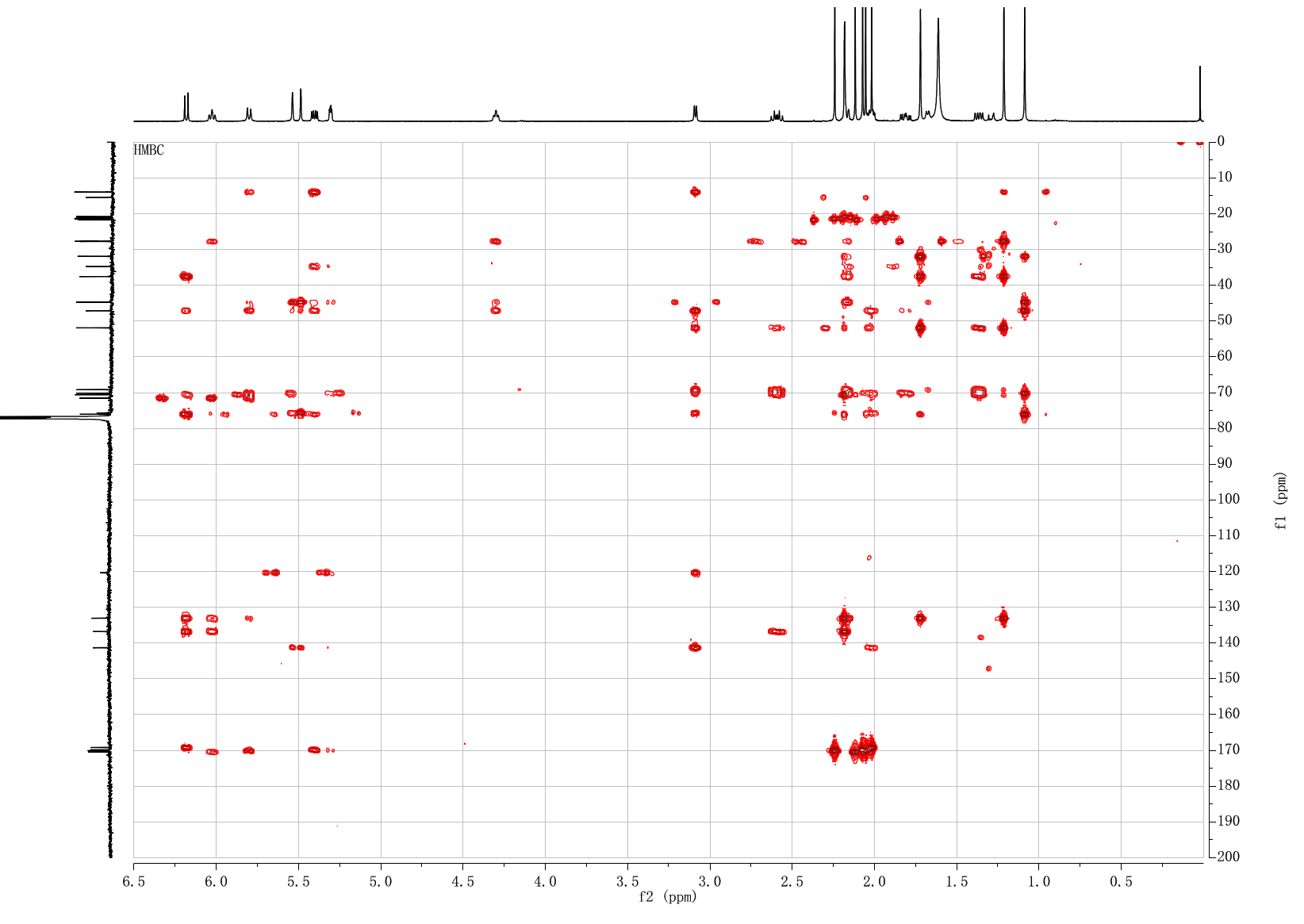

**Figure S54. ^1^H-^13^C HMBC NMR spectrum of 7*β*-acetoxytauxin-2*α*-ol (12).**

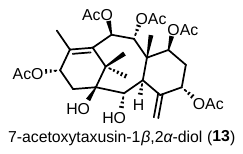

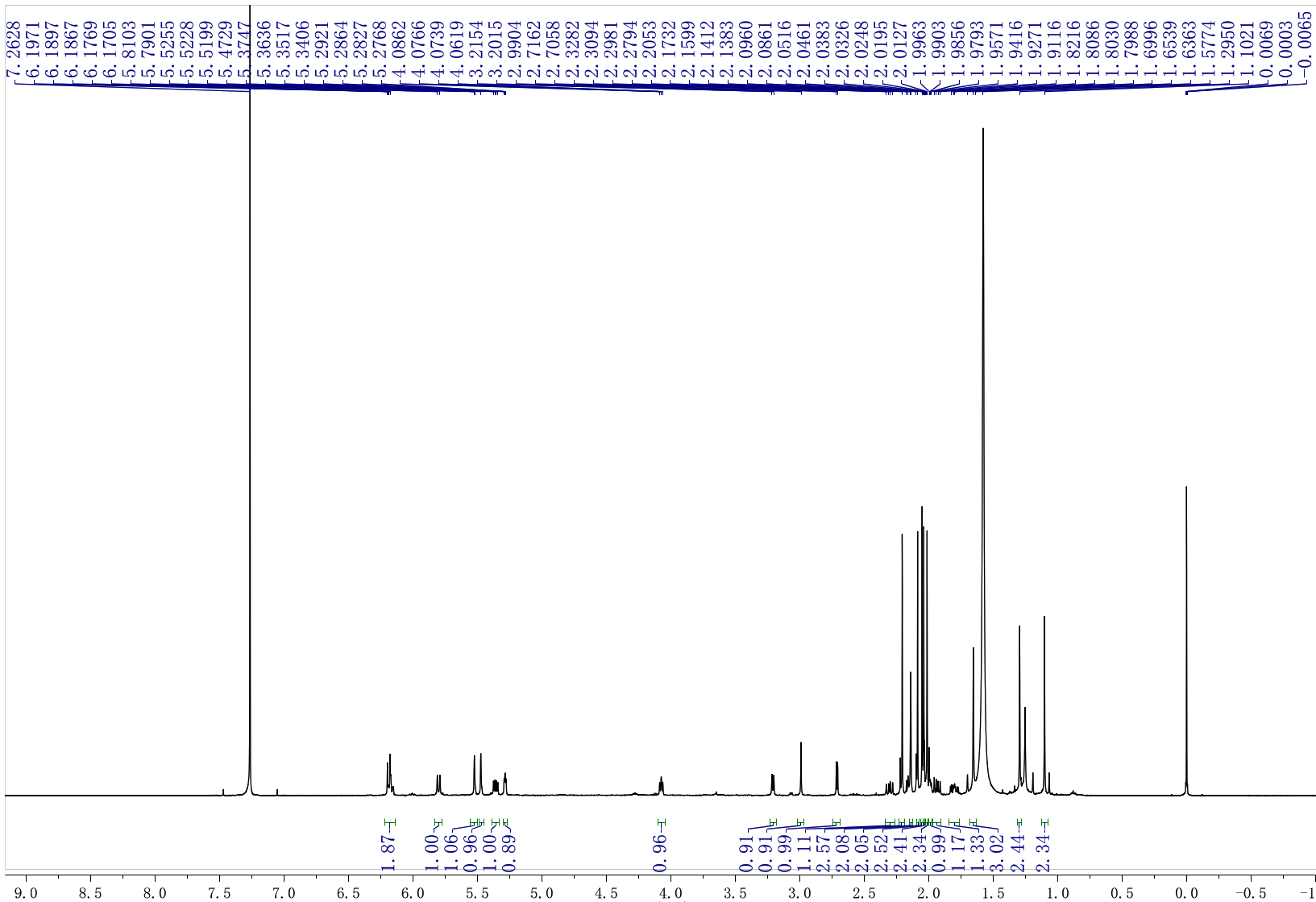

**Figure S55. ^1^H NMR spectrum of 7*β*-acetoxytauxin-1*β*, 2*α*-ol (13).**

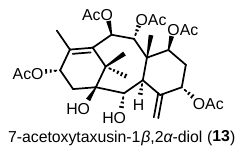

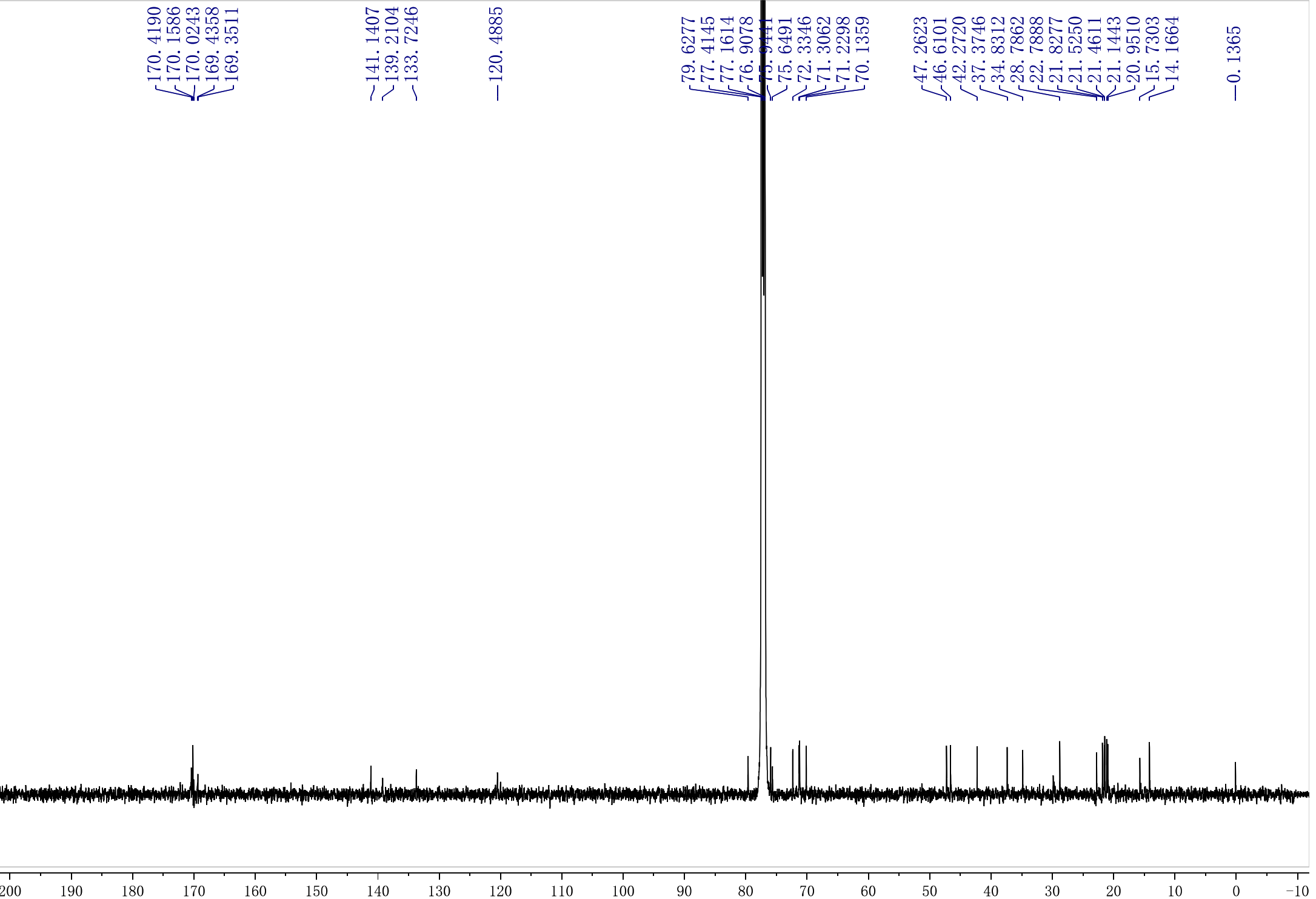

**Figure S56. ^13^C NMR spectrum of 7*β*-acetoxytauxin-1*β*, 2*α*-ol (13).**

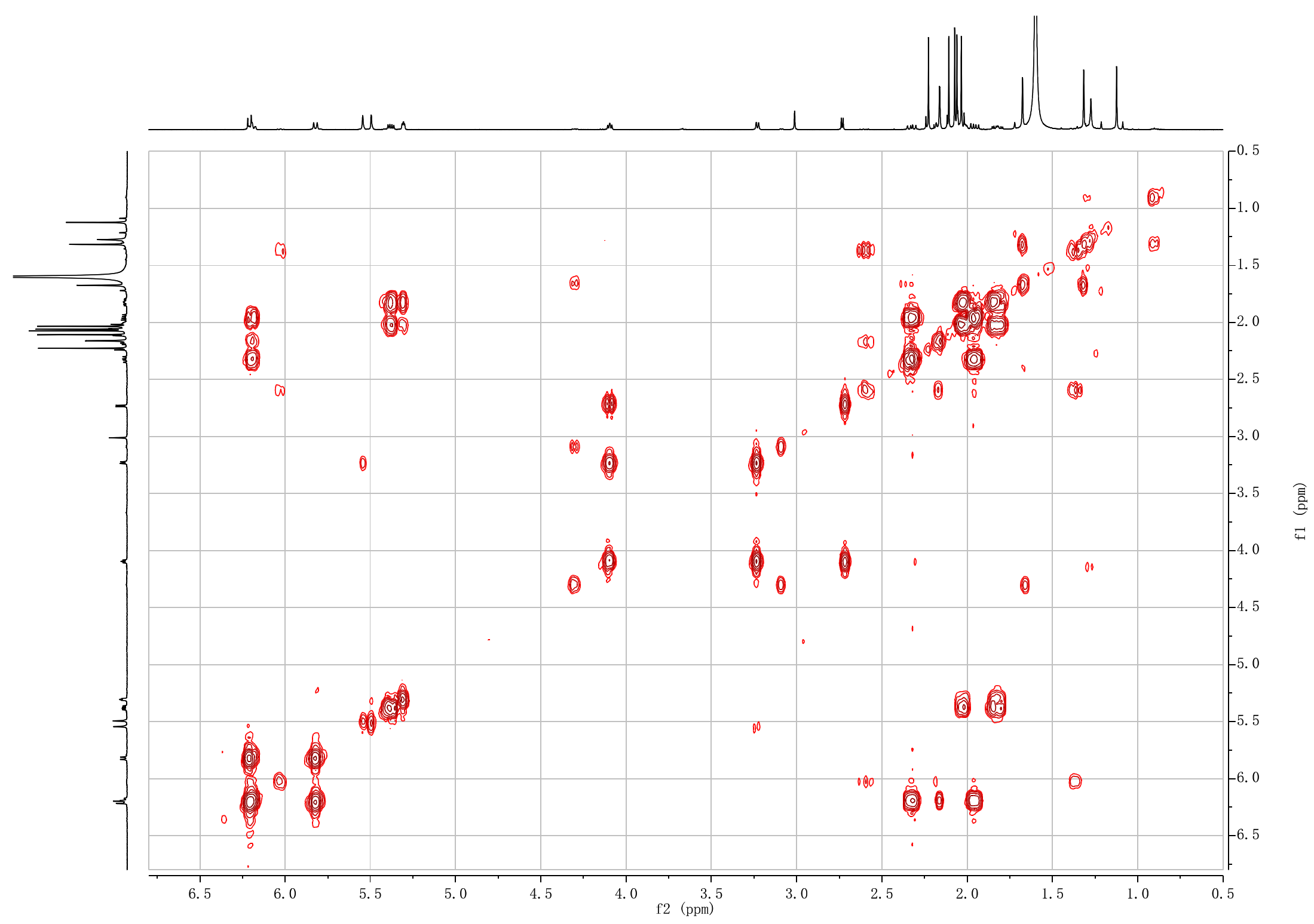

**Figure S57. ^1^H-^1^H COSY NMR spectrum of 7*β*-acetoxytauxin-1*β*, 2*α*-ol (13).**

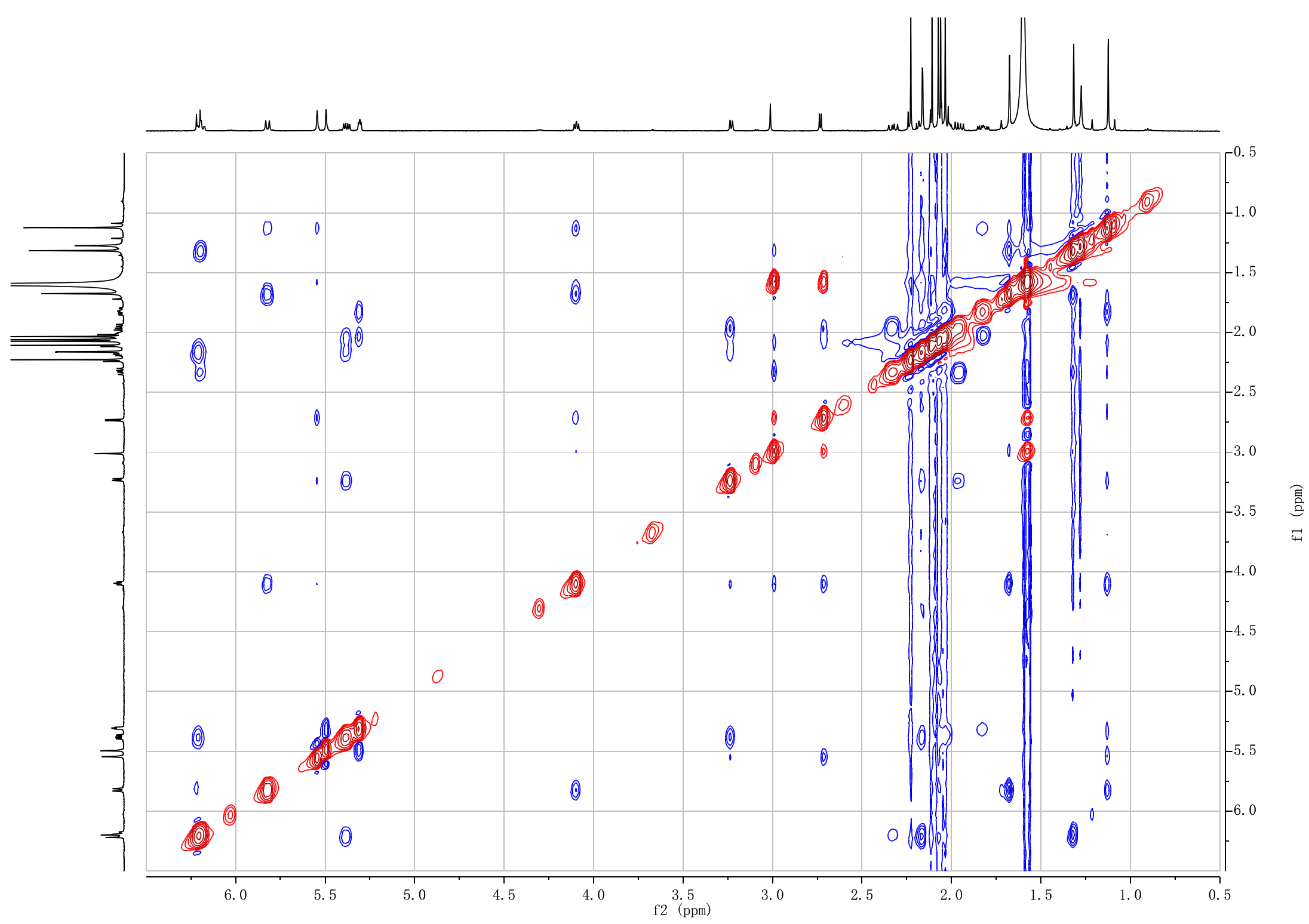

**Figure S58. ^1^H-^1^H NOESY NMR spectrum of 7*β*-acetoxytauxin-1*β*, 2*α*-ol (13).**

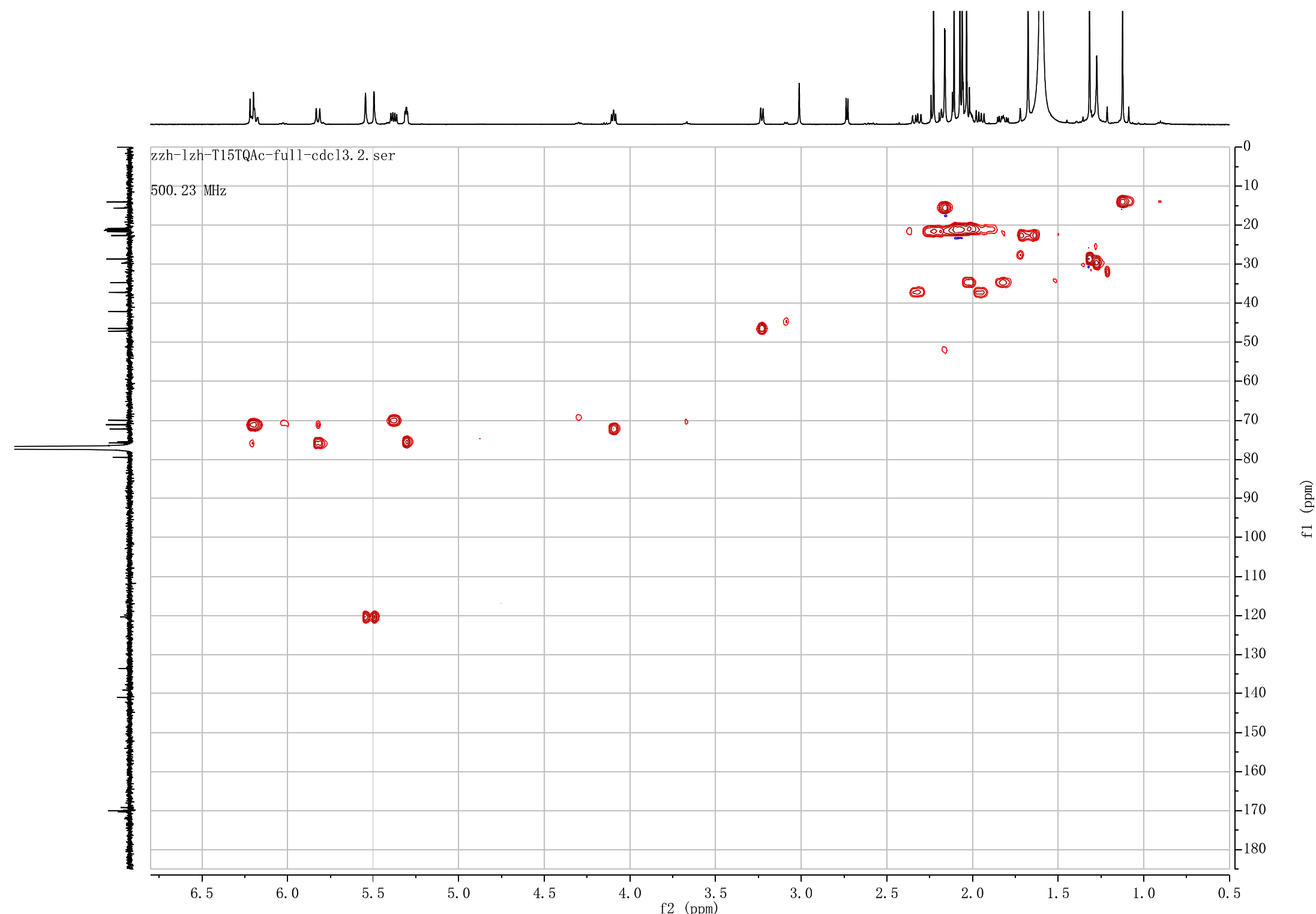

**Figure S59. ^1^H-^13^C HSQC NMR spectrum of 7*β*-acetoxytauxin-1*β*, 2*α*-ol (13).**

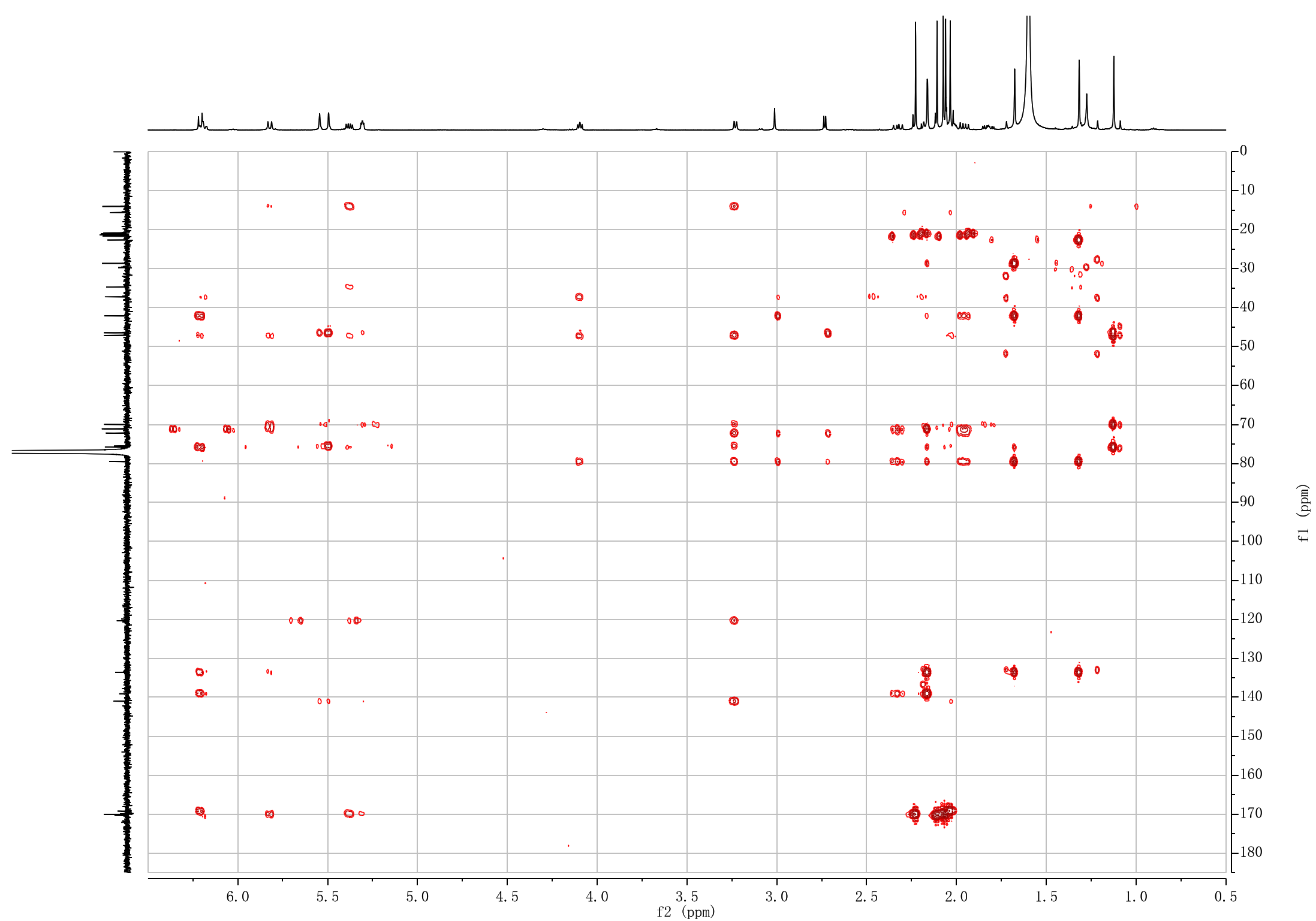

**Figure S60. ^1^H-^13^C HMBC NMR spectrum of 7*β*-acetoxytauxin-1*β*, 2*α*-ol (13).**

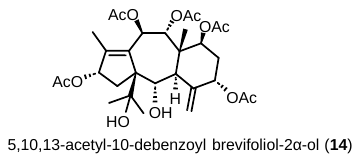

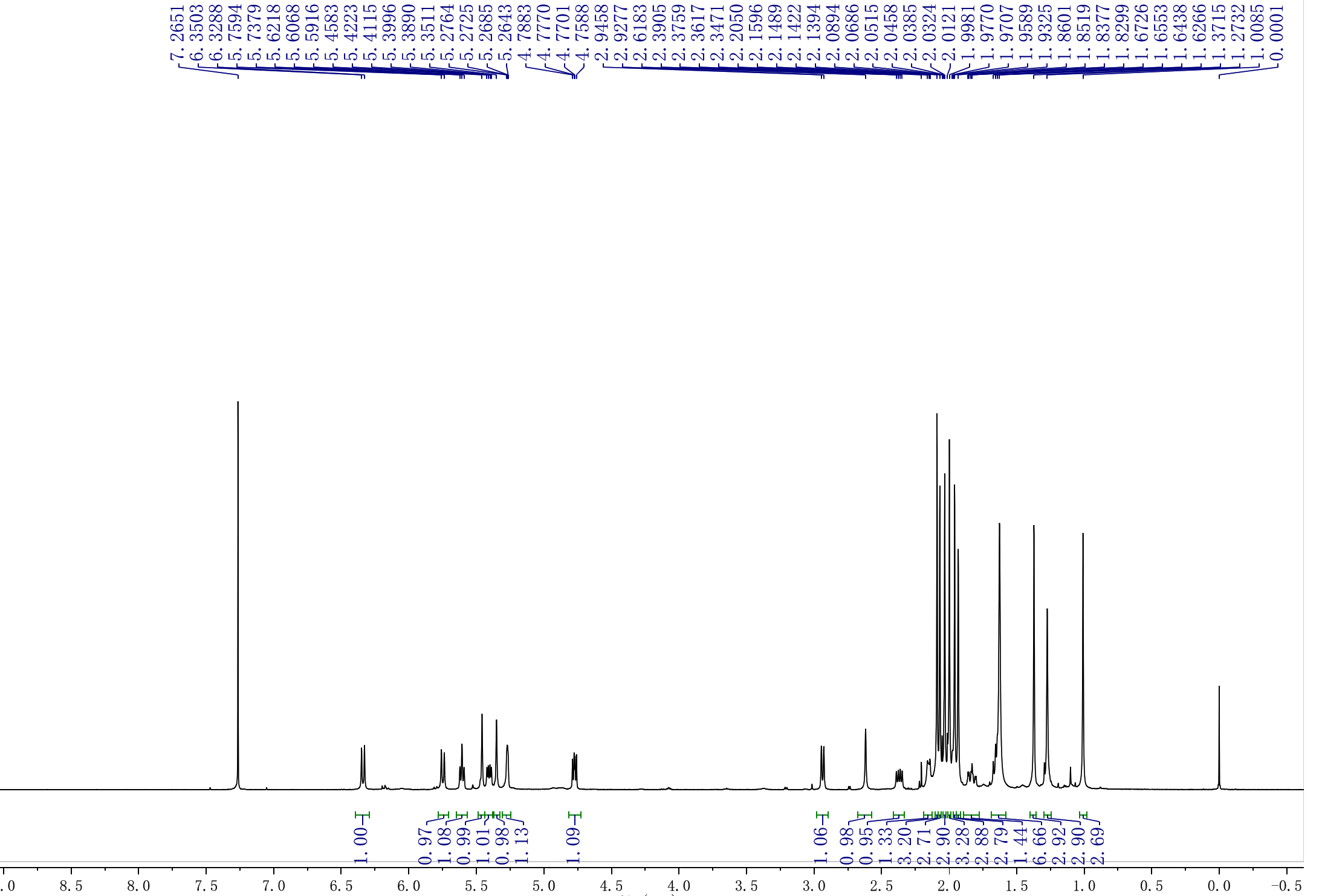

**Figure S61. ^1^H NMR spectrum of 5,10,13-acetyl-10-debenzoylbrevifoliol-2*α*-ol (14).**

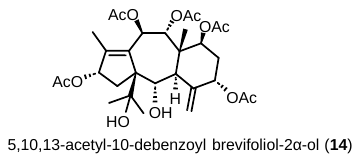

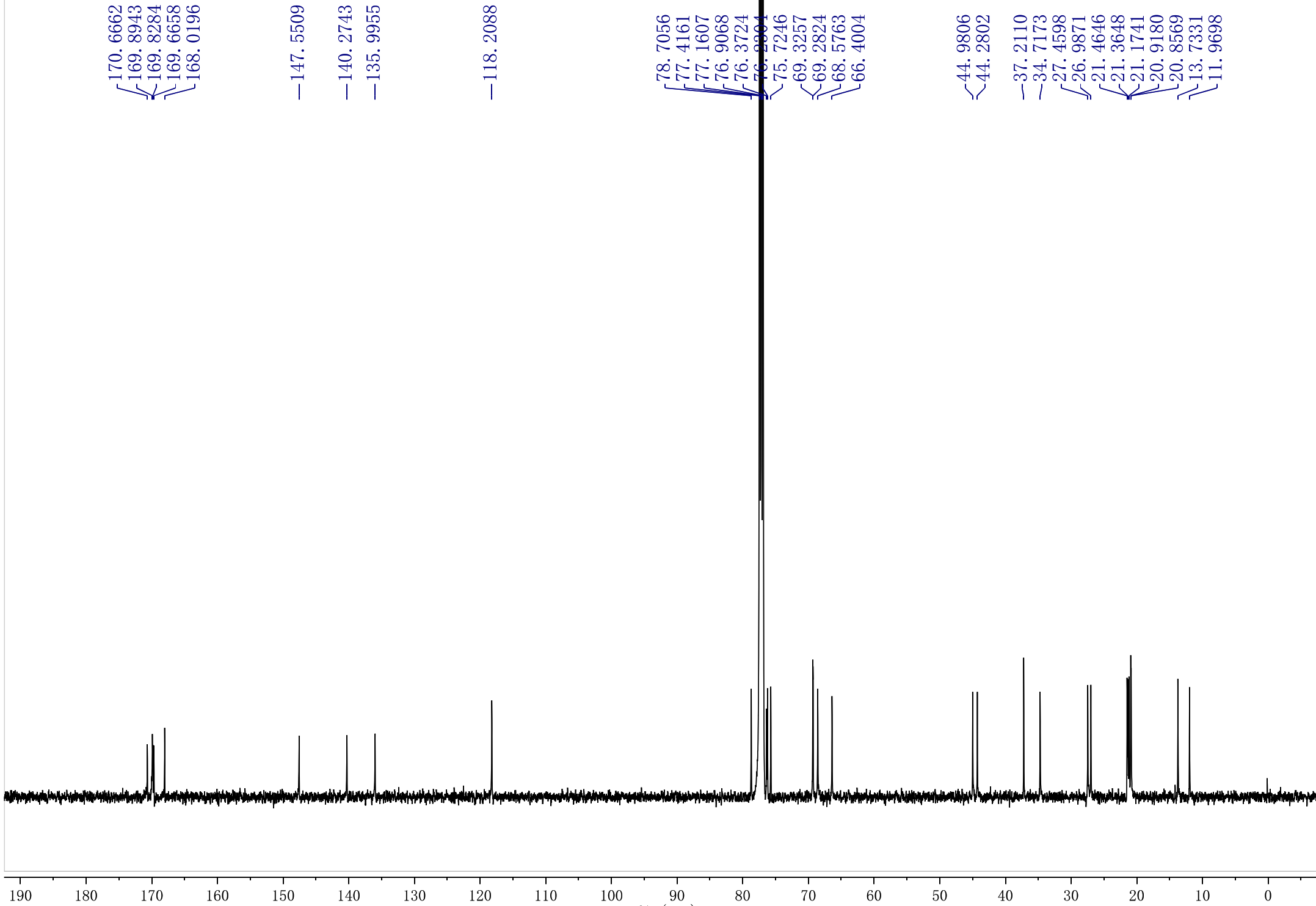

**Figure S62. ^13^C NMR spectrum of 5,10,13-acetyl-10-debenzoylbrevifoliol-2*α*-ol (14).**

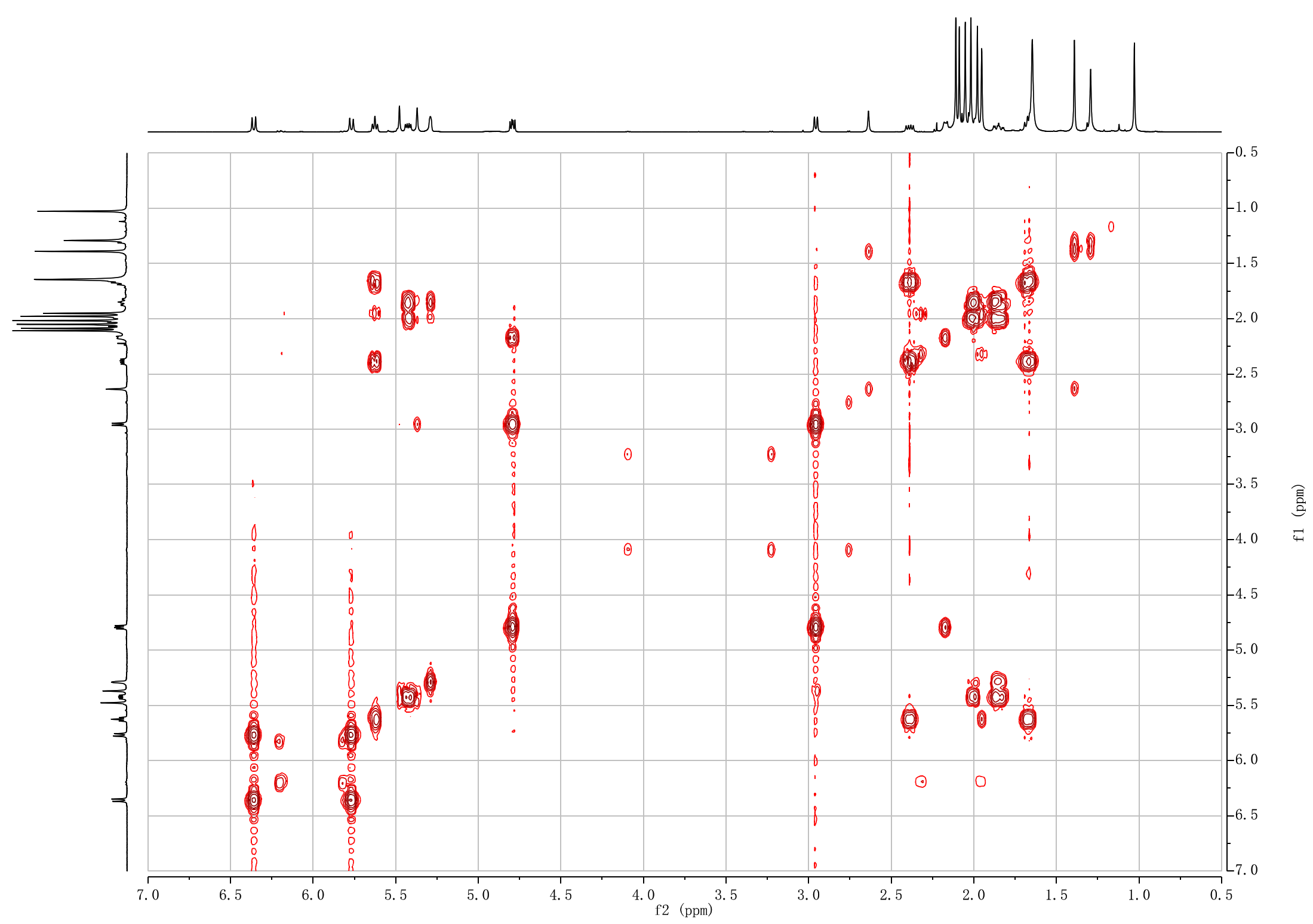

**Figure S63. ^1^H-^1^H COSY NMR spectrum of 5,10,13-acetyl-10-debenzoyl brevifoliol-2*α*-ol (14).**

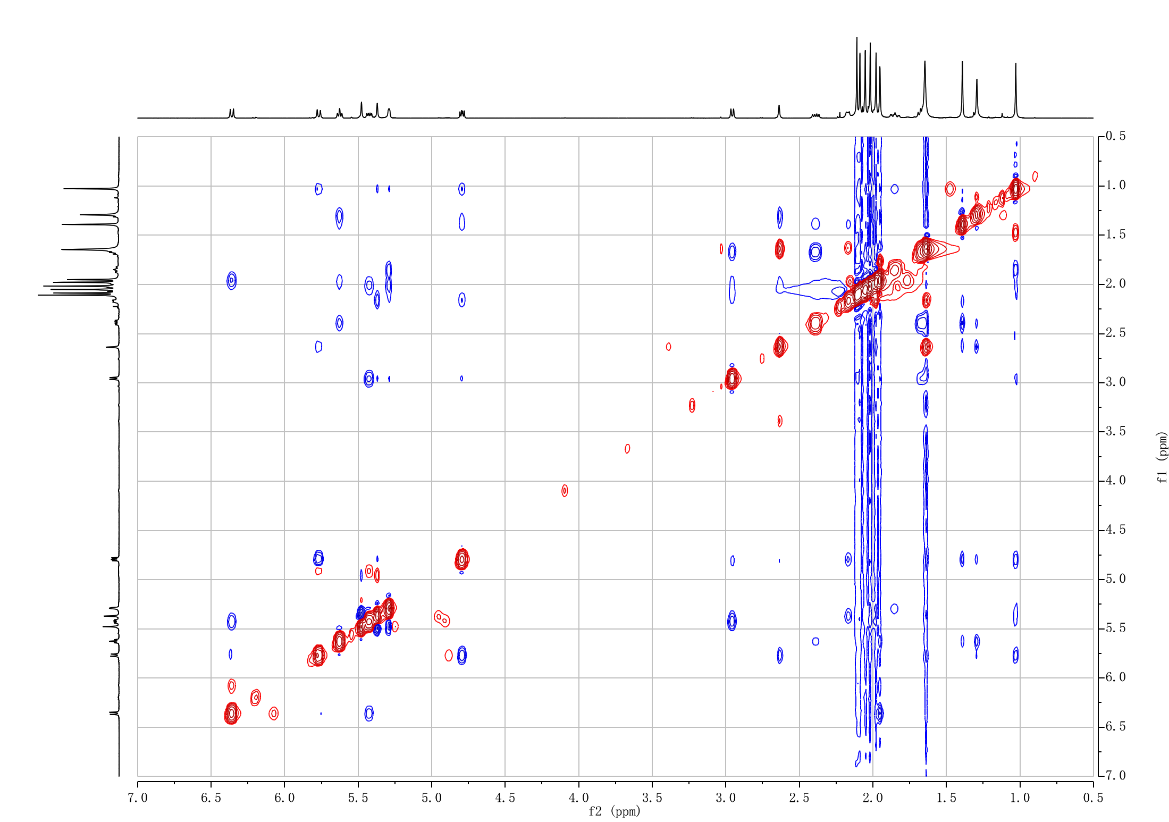

**Figure S64. ^1^H-^1^H NOESY NMR spectrum of 5,10,13-acetyl-10-debenzoyl brevifoliol-2*α*-ol (14).**

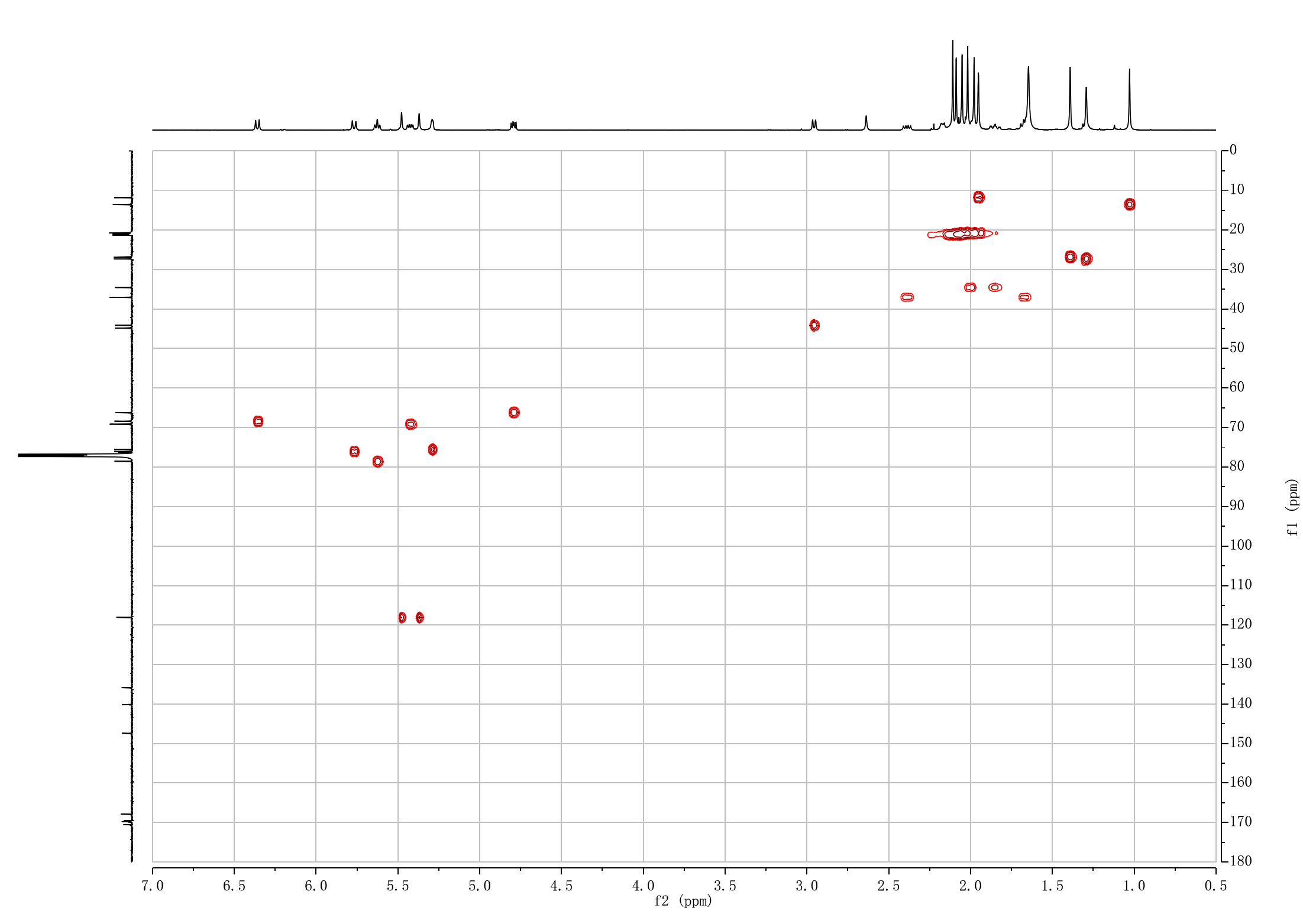

**Figure S65. ^1^H-^13^C HSQC NMR spectrum of 5,10,13-acetyl-10-debenzoyl brevifoliol-2*α*-ol (14).**

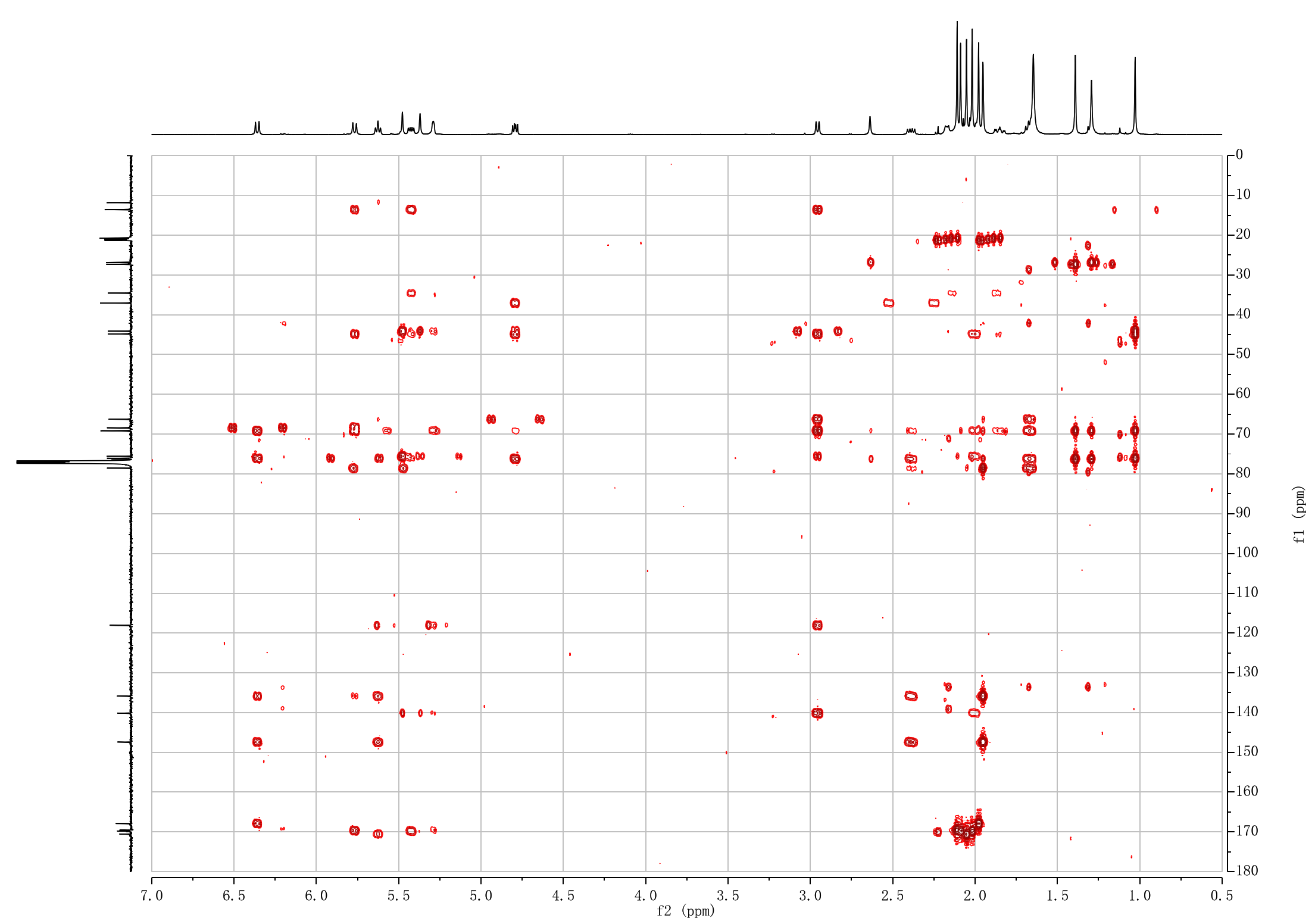

**Figure S66. ^1^H-^13^C HMBC NMR spectrum of 5,10,13-acetyl-10-debenzoyl brevifoliol-2*α*-ol (14).**
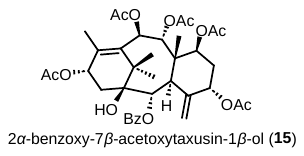

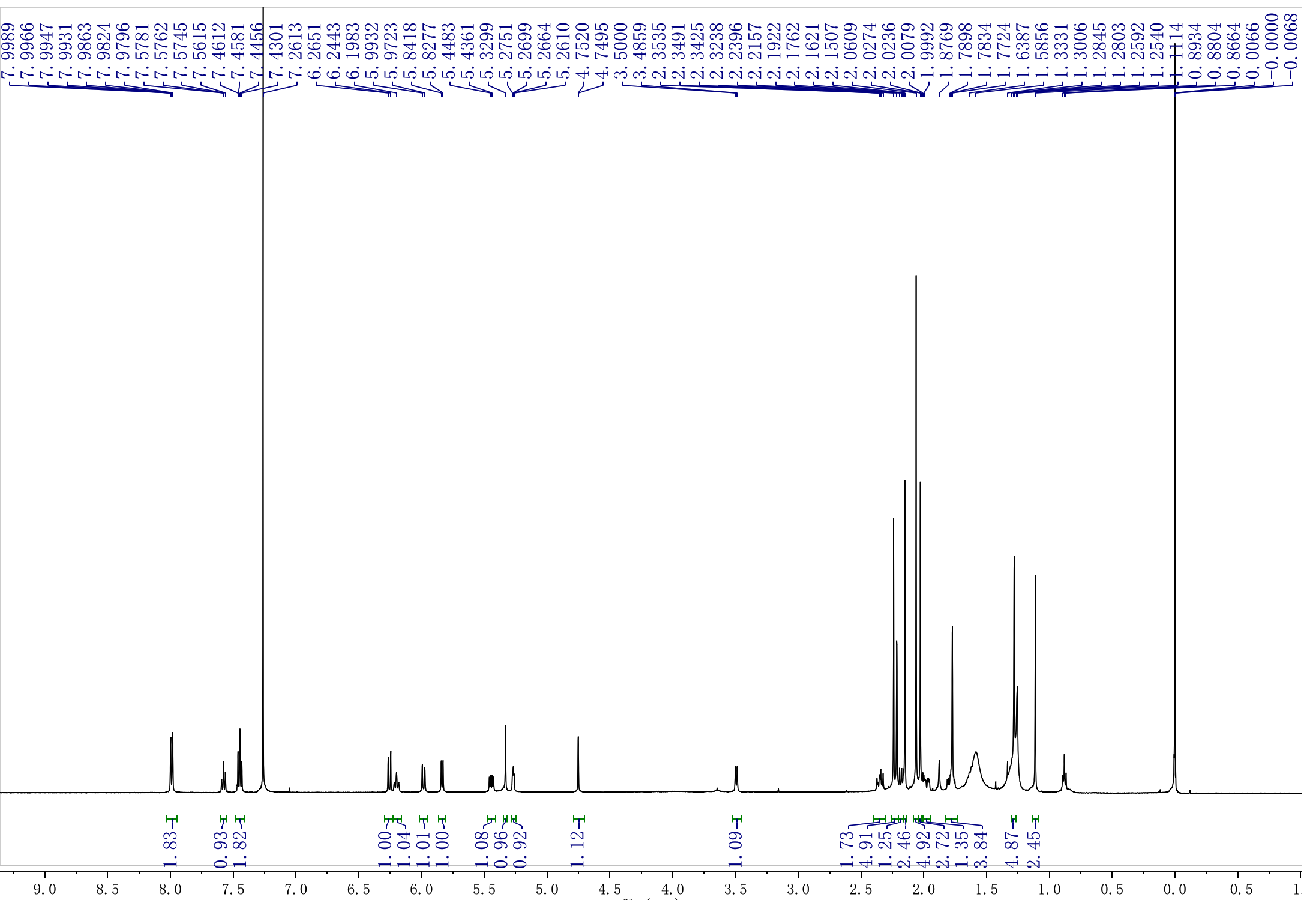

**Figure S67. ^1^H NMR spectrum of 2*α*-benzoxy-7*β*-acetoxytaxusin-1*β*-ol (15).**

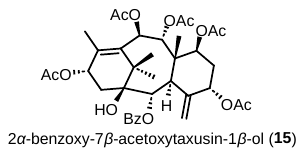

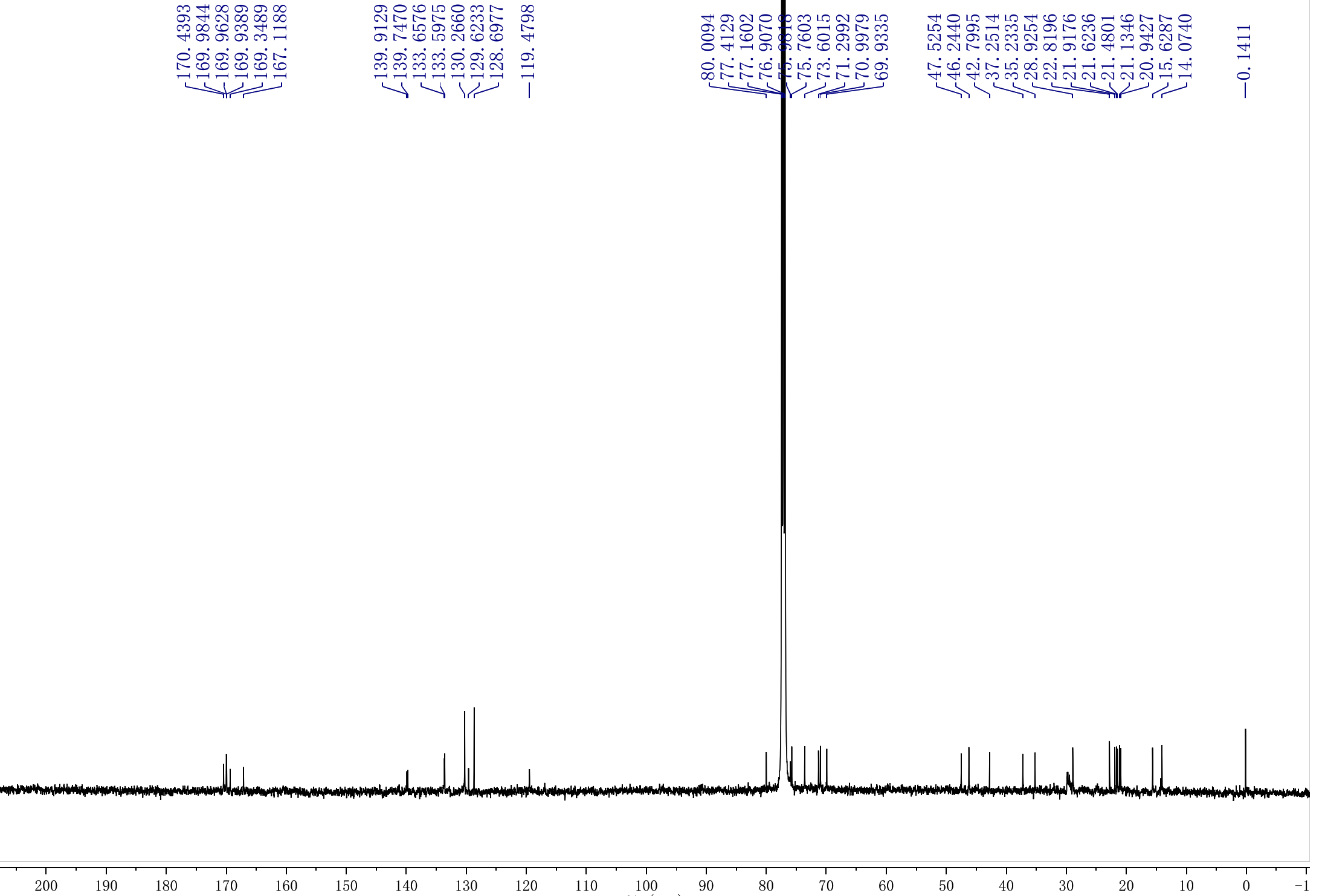

**Figure S68. ^13^C NMR spectrum of 2*α*-benzoxy-7*β*-acetoxytaxusin-1*β*-ol (15).**

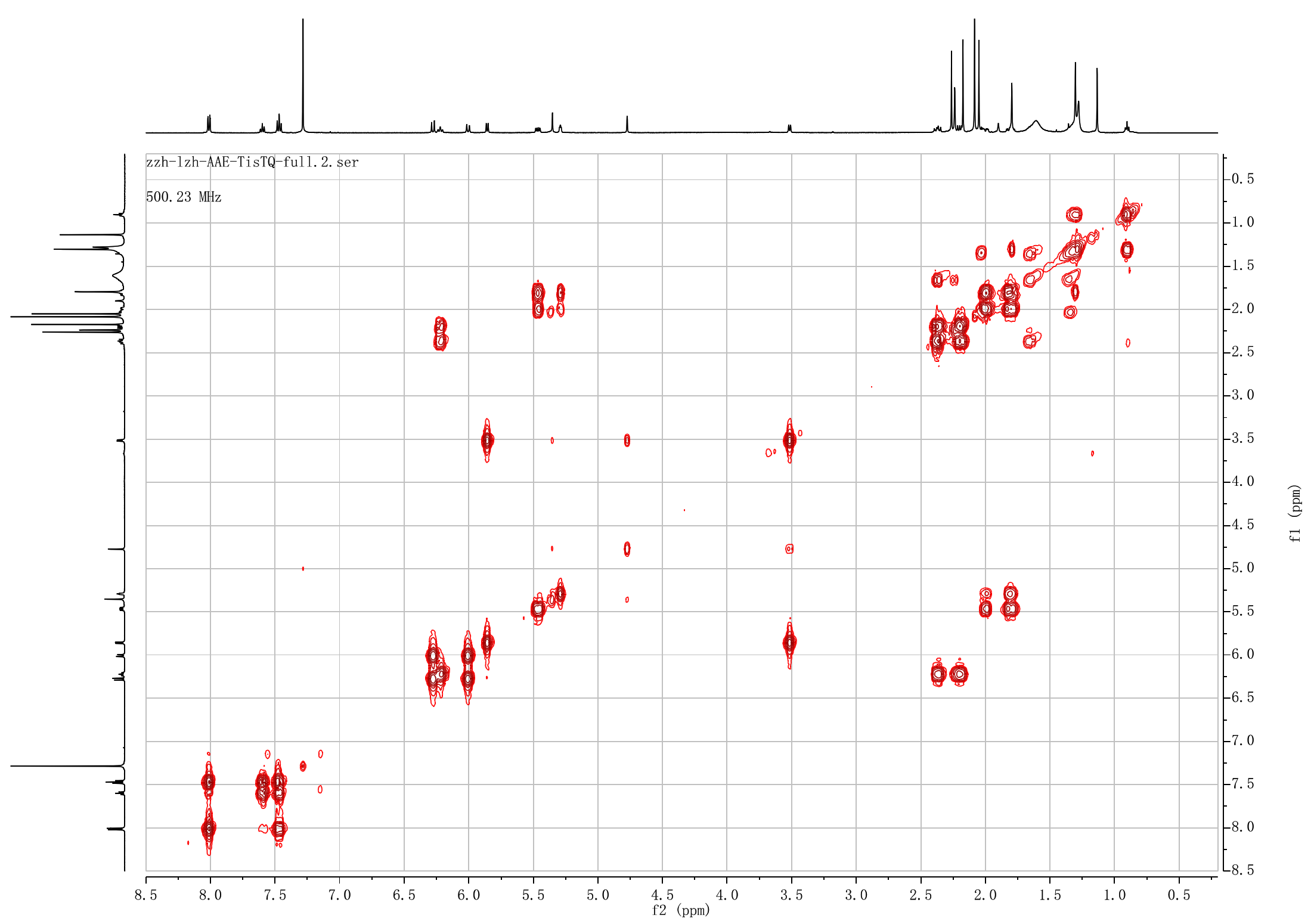

**Figure S69. ^1^H-^1^H COSY NMR spectrum of 2*α*-benzoxy-7*β*-acetoxytaxusin-1*β*-ol (15)**

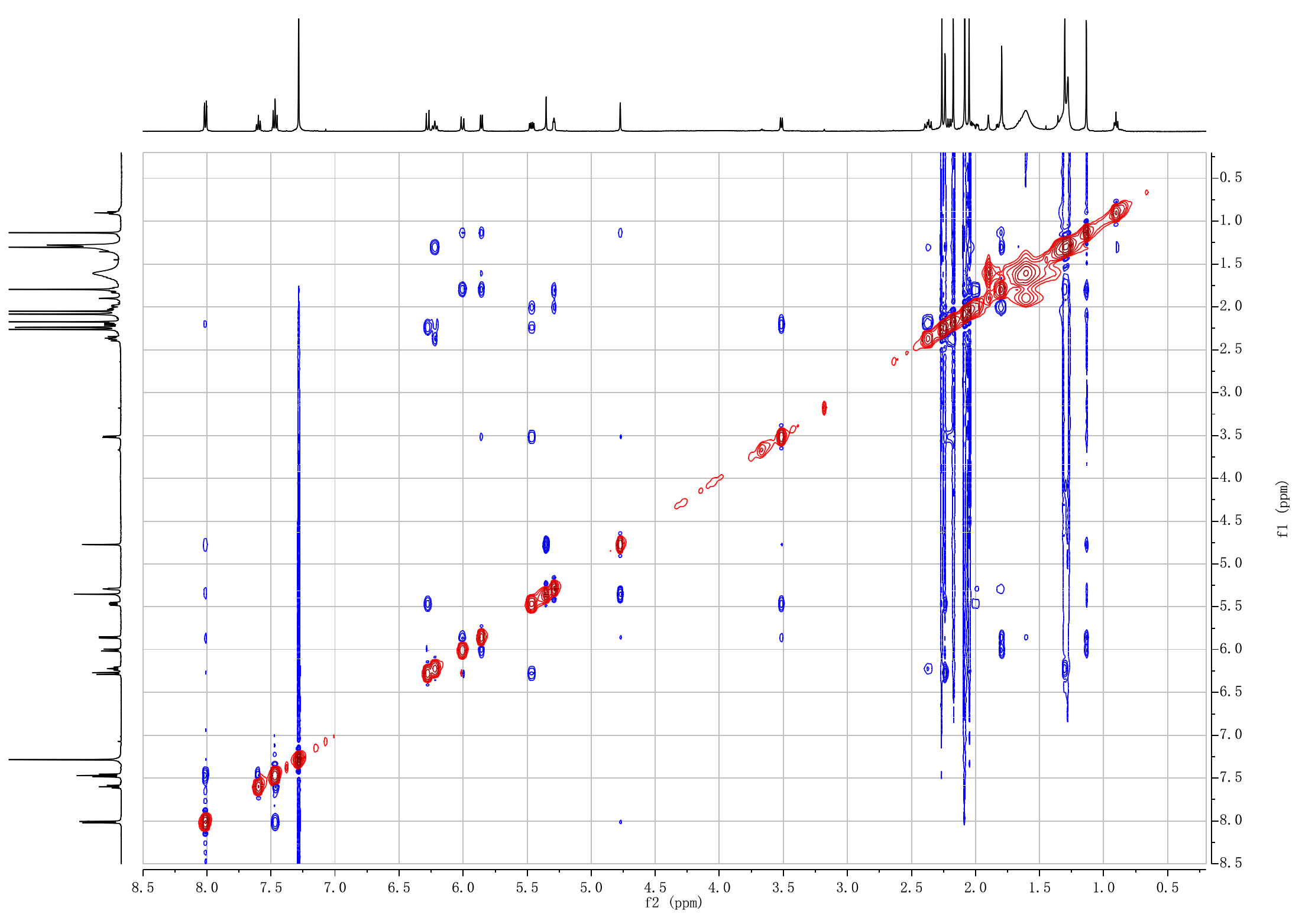

**Figure S70. ^1^H-^1^H NOESY NMR spectrum of 2*α*-benzoxy-7*β*-acetoxytaxusin-1*β*-ol (15)**

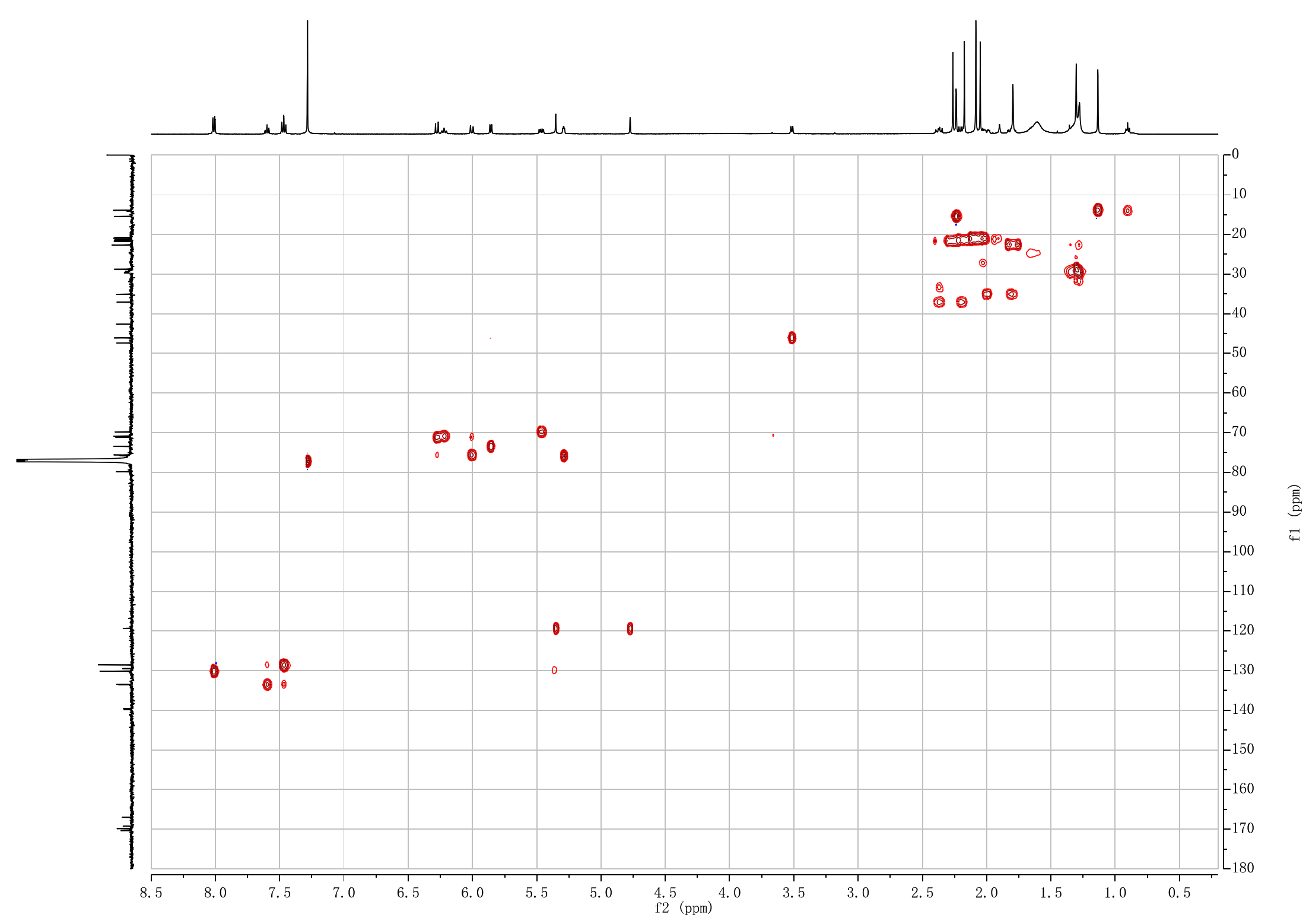

**Figure S71. ^1^H-^13^C HSQC NMR spectrum of 2*α*-benzoxy-7*β*-acetoxytaxusin-1*β*-ol (15).**
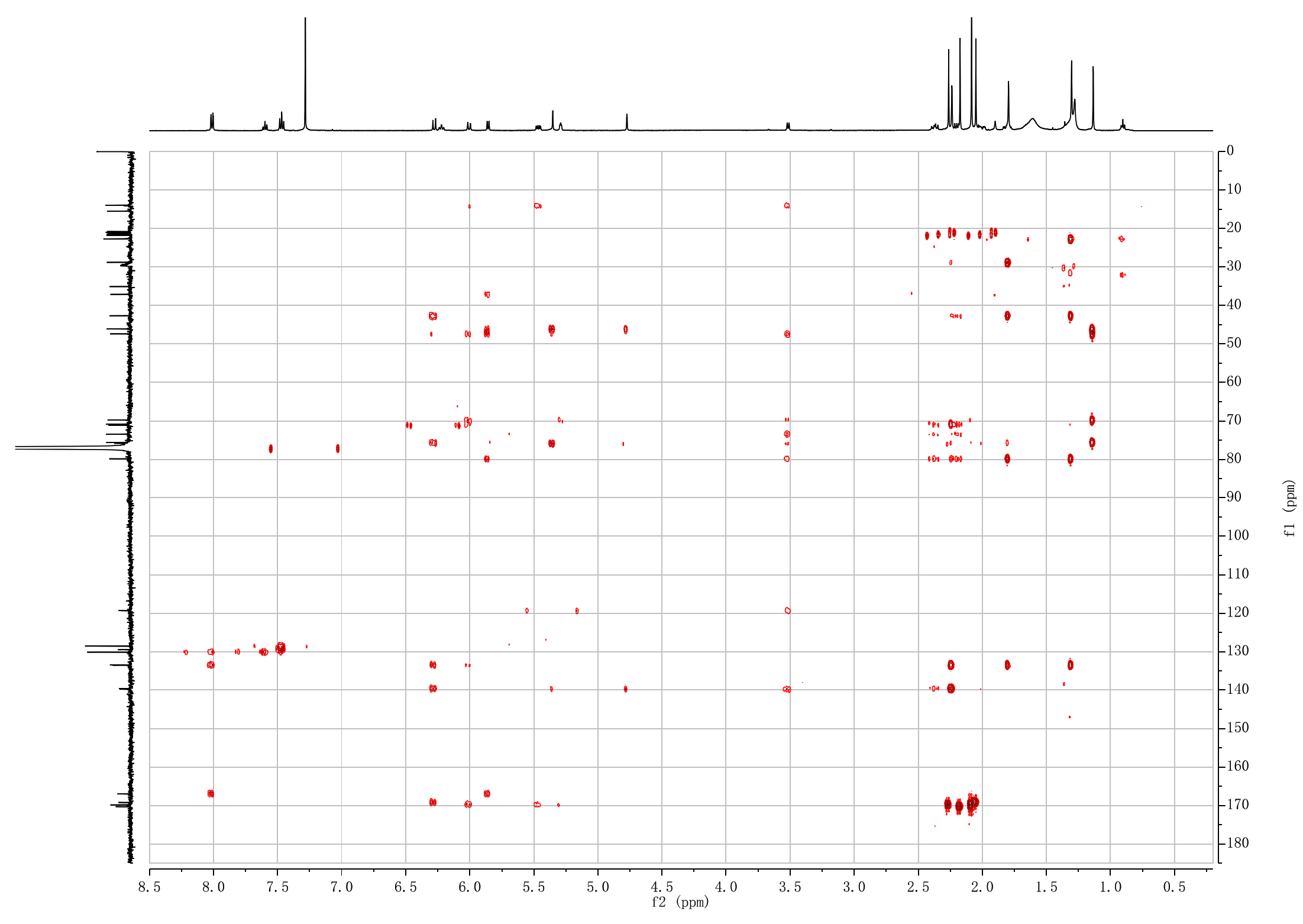

**Figure S72. ^1^H-^13^C HMBC NMR spectrum of 2*α*-benzoxy-7*β*-acetoxytaxusin-1*β*-ol (15).**

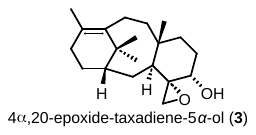

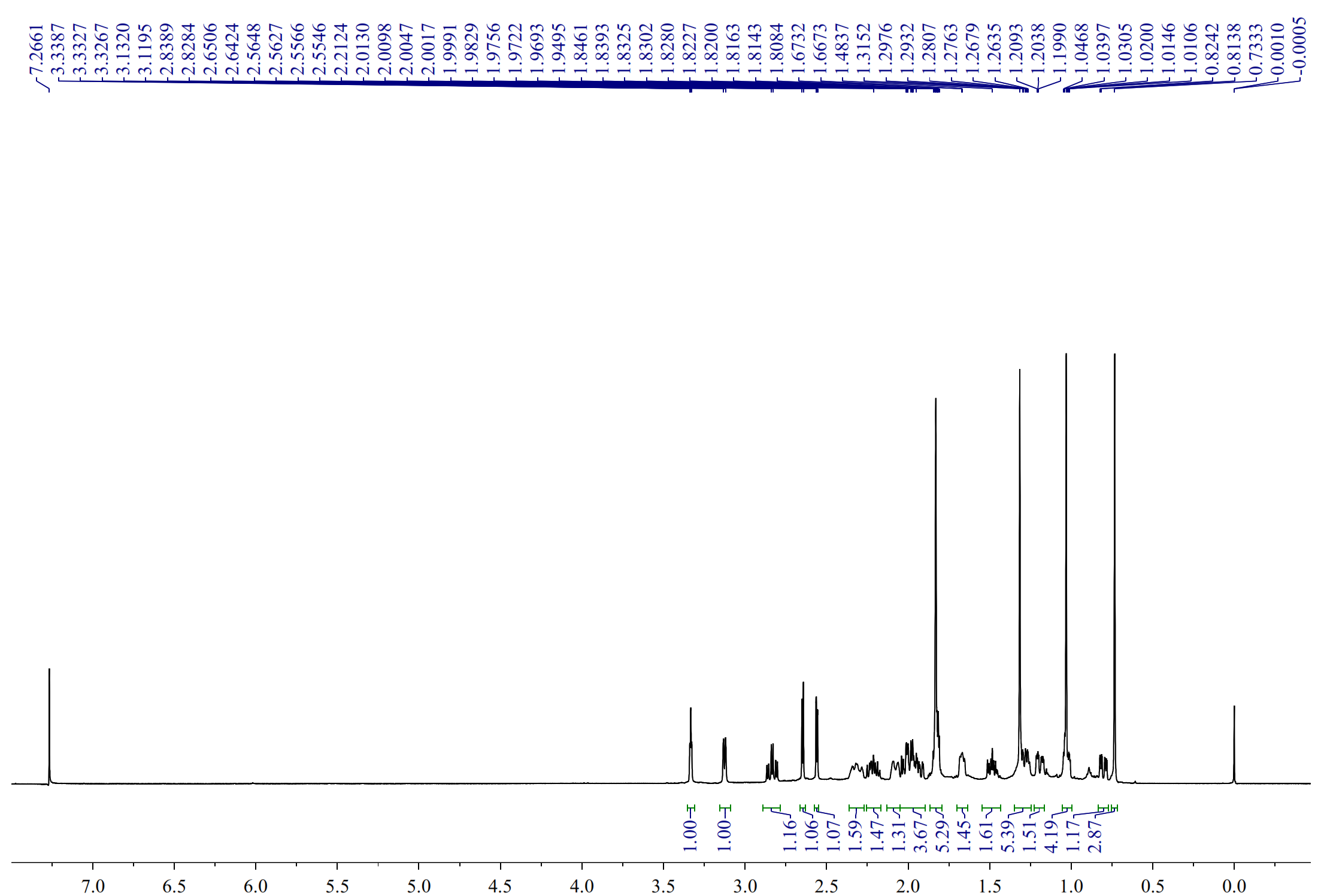

**Figure S73. ^1^H NMR spectrum of 4*α*,20-epoxide-taxadiene-5*α*-ol (3).**

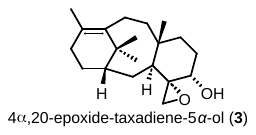

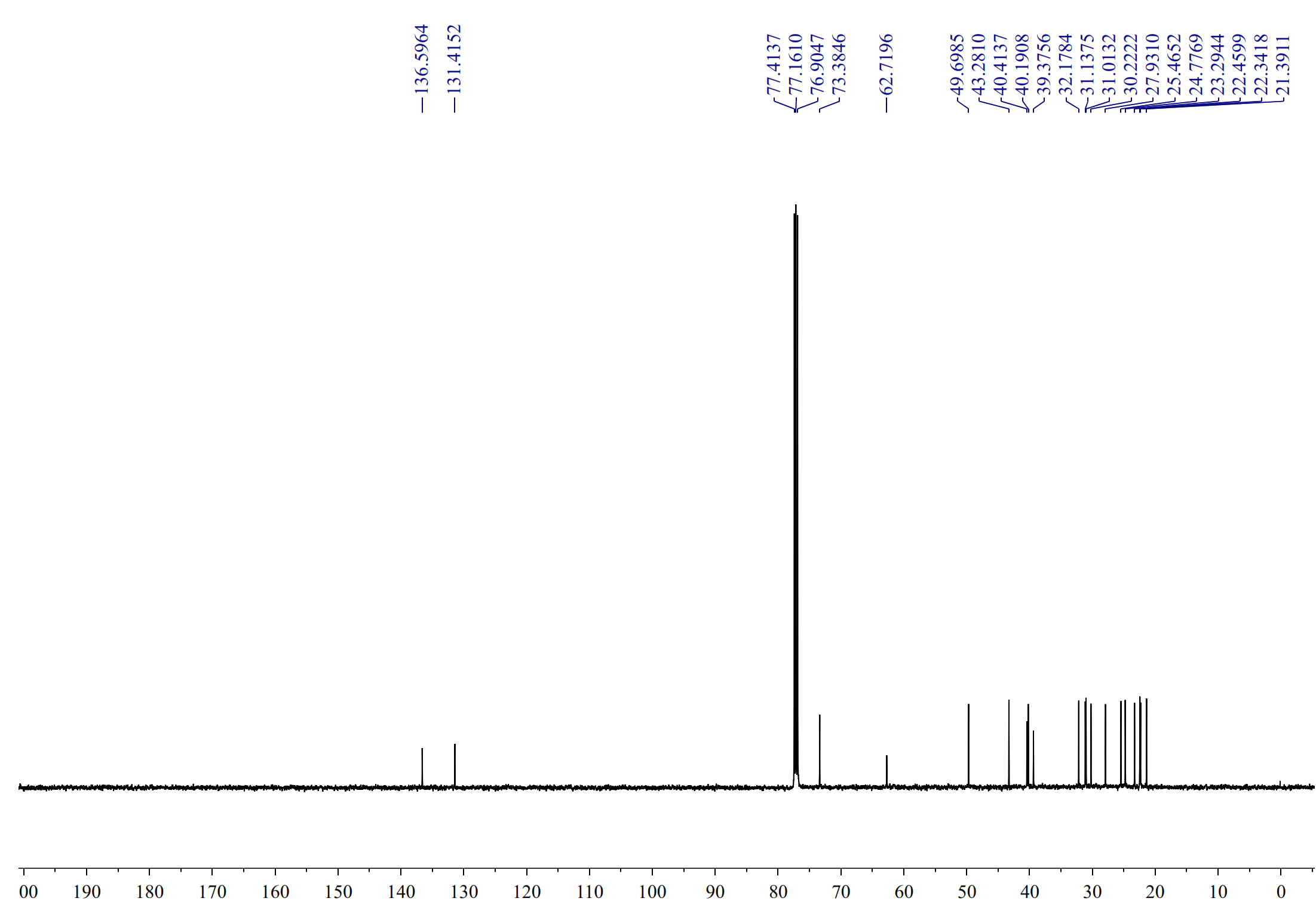

**Figure S74.^13^C NMR spectrum of 4*α*,20-epoxide-taxadiene-5*α*-ol (3).**

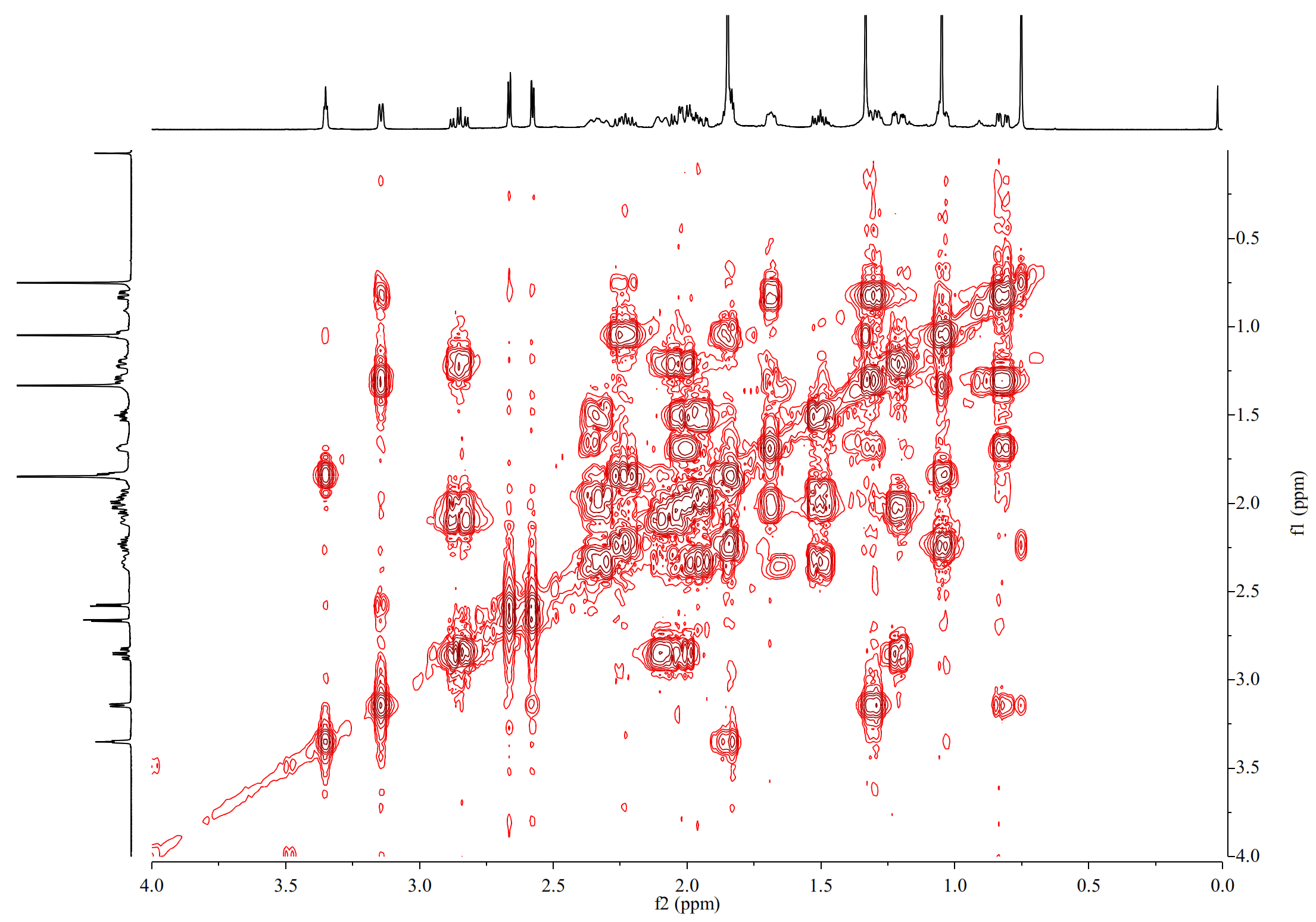

**Figure S75. ^1^H-^1^H COSY NMR spectrum of 4*α*,20-epoxide-taxadiene-5*α*-ol (3).**

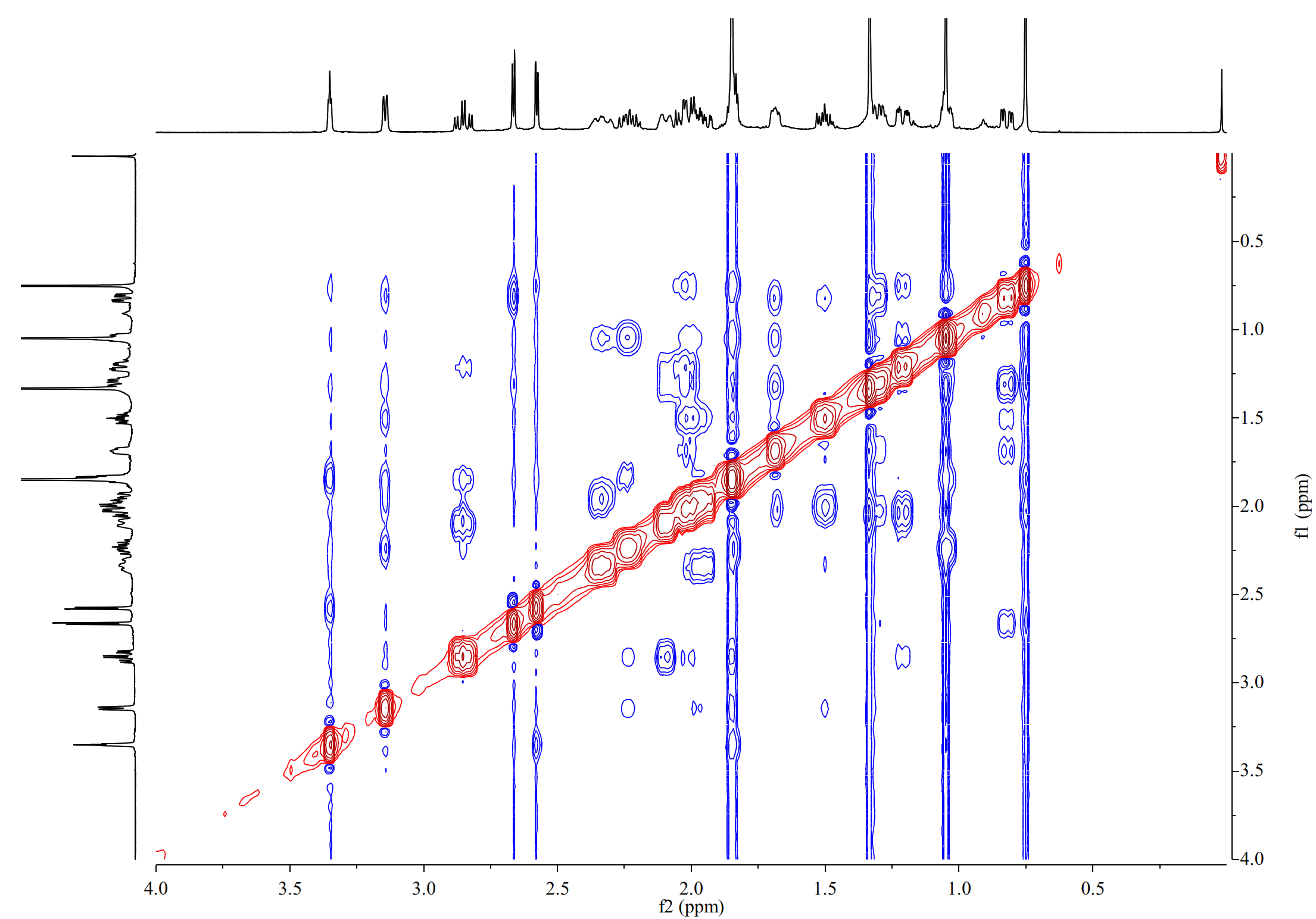

**Figure S76. ^1^H-^1^H NOESY NMR spectrum of 4α,20-epoxide-taxadiene-5*α*-ol (3).**

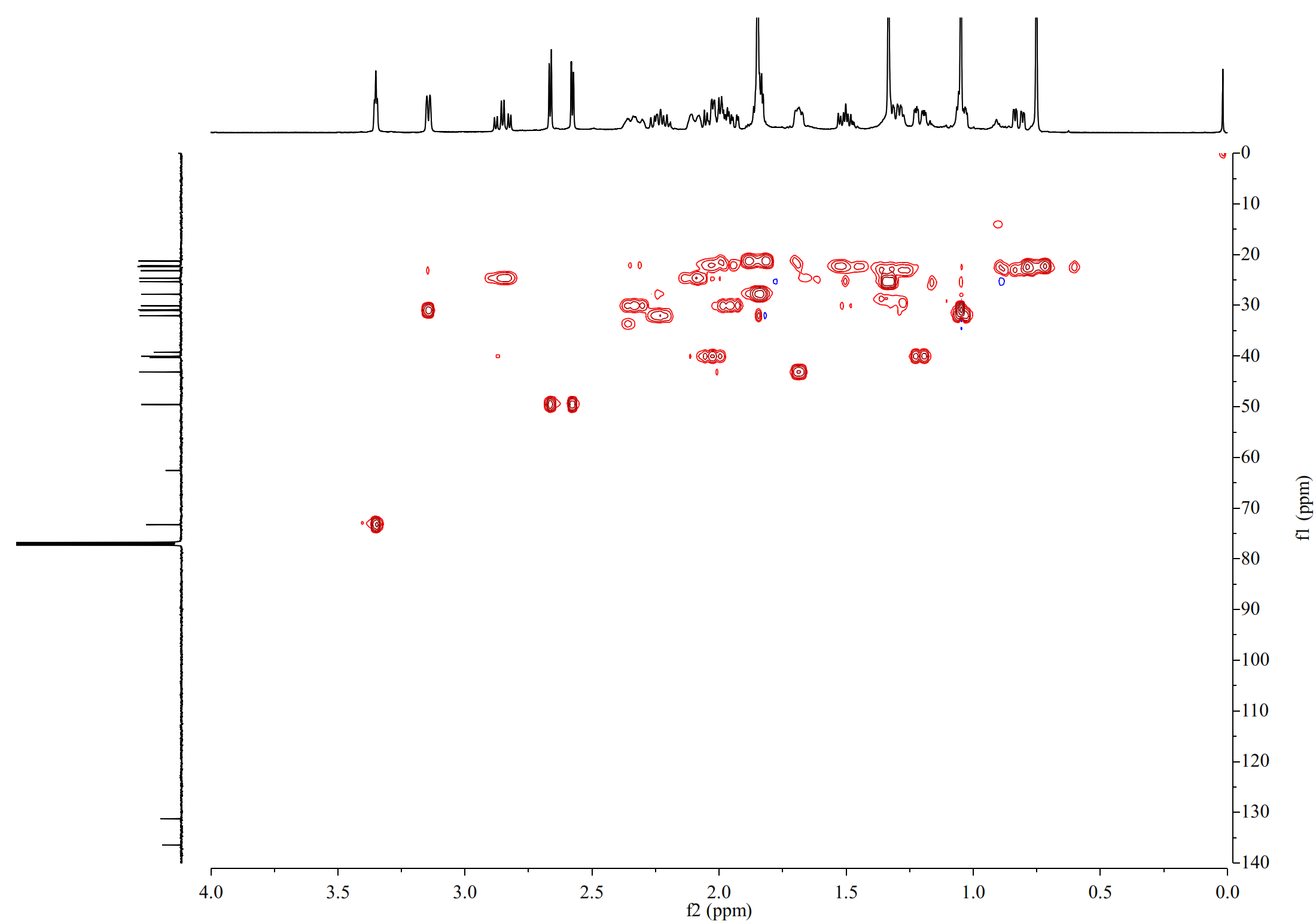

**Figure S77. ^1^H-^13^C HSQC NMR spectrum of 4*α*,20-epoxide-taxadiene-5*α*-ol (3).**

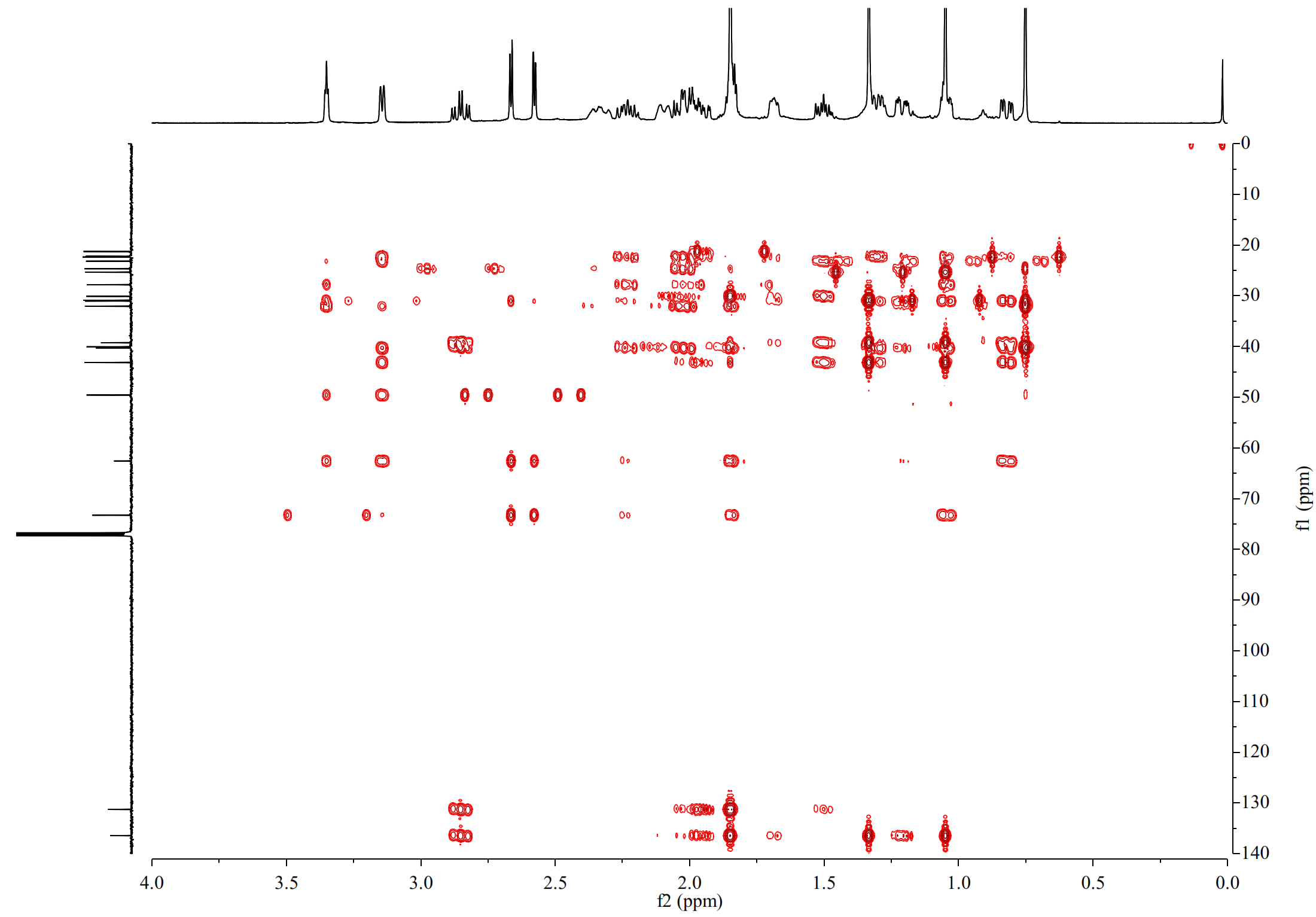
**Figure S78. ^1^H-^13^C HMBC NMR spectrum of 4*α*,20-epoxide-taxadiene-5*α*-ol (3).**

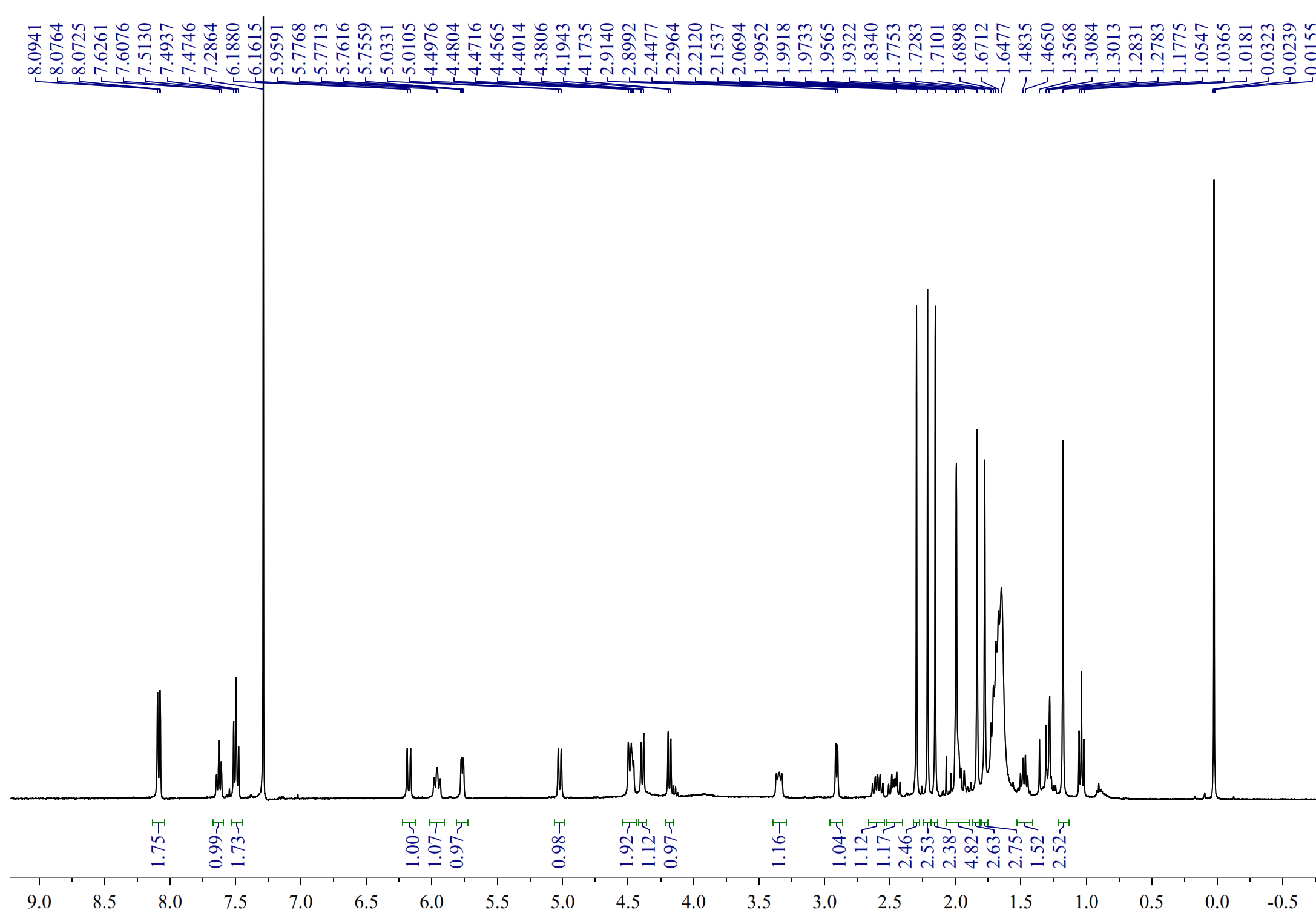

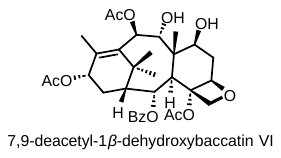

**Figure S79. ^1^H NMR spectrum of 7,9-deacetyl-1*β*-dehydroxybaccatin VI.**

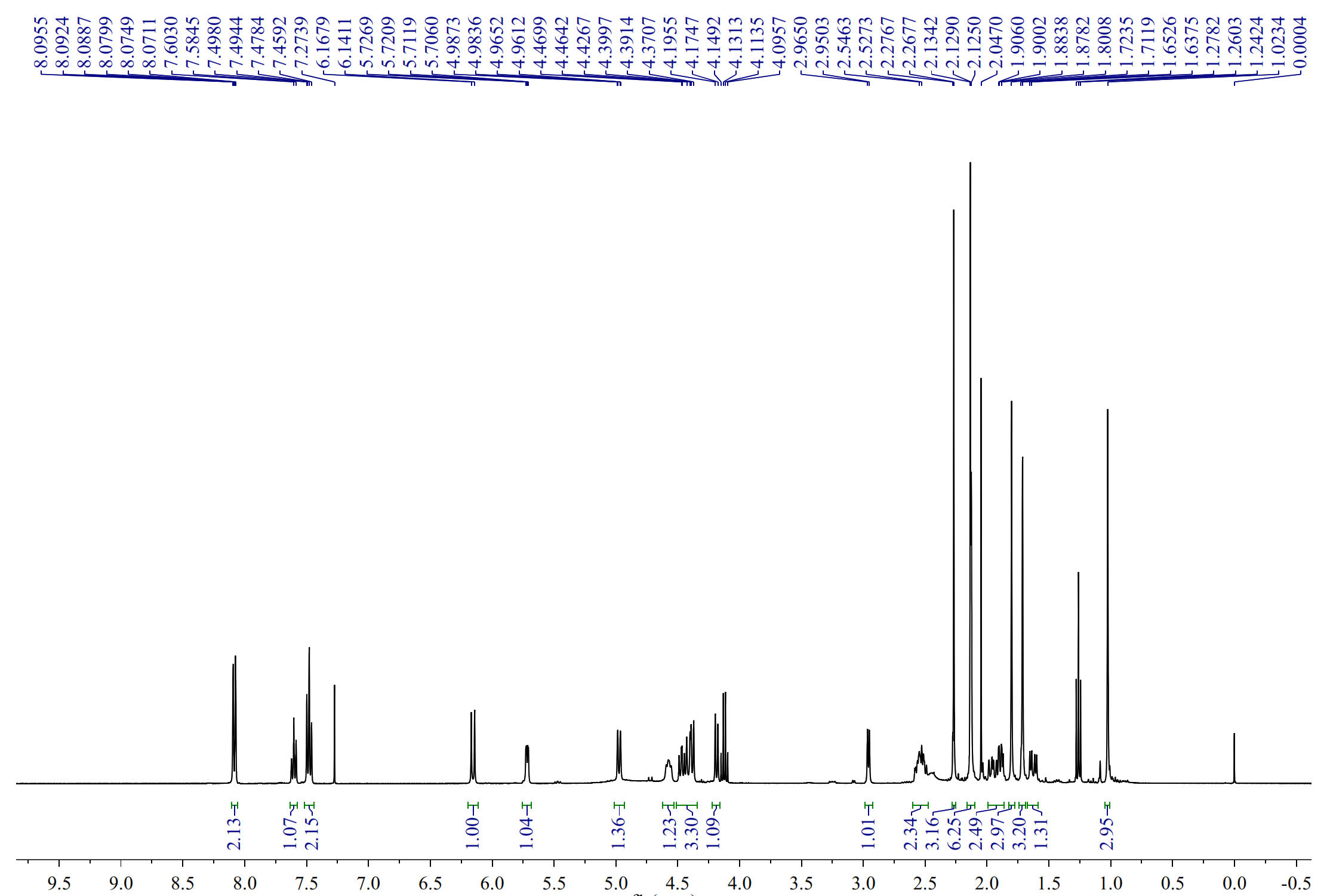

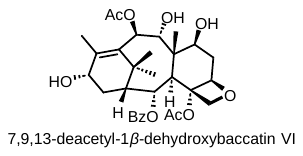

**Figure S80. ^1^H NMR spectrum of 7,9,13-deacetyl-1*β*-dehydroxybaccatin VI.**

**Table S1. Strains used in this study.**

| **Stain** | **Genotype/Description** | **Source** |
| --- | --- | --- |
| BY4742 | MATα, his3Δ1, leu2Δ0, lys2Δ0, ura3Δ0 | EUROSCARF |
| YBD80 | BY4742 {gal80::URA3} | Ref.^2^ |
| YTCPR | YBD80 {deltaDNA::GAL10p-*synTCPR*-TDH3t-LEU2} | Ref.^2^ |
| YT13OH | YBD80 {rDNA::GAL10p-synTCPR-TDH3t/GAL1p-synT13OH-GPM1t-HIS3} | Ref.^2^ |
| YCYP725A55 | YTCPR {rDNA:: GAL1p-CYP725A55-GPM1t-HIS3} | Ref.^2^ |
| YAAE | YBD80 {deltaDNA::GAL1p-synAAE-HXT7t-LEU2} | Ref.^2^ |
| YDBVI | YT02 {rDNA::GAL1p-synAAE4-TDH2t/GAL10p-synTBT-HXT7t/GAL1p-CYP725A55-PDC1t/GAL10p-AT5-PYK1t-KanMX} | Ref.^2^ |
| YSP | YBD80 {deltaDNA:: GAL1p-IDI1-PGIt/GAL10p-*GGPPSaPS*-ENO2t/ GAL1p-ERG19-ADH1t/GAL10p-tHMG*1*-CYC1t/-LEU2} | This study |
| YT5OH | YSP{rDNA::GAL10p-synTCPR-TDH3t/GAL1p-synT5OH-GPM1t-HIS3} | This study |
| YT5OH_M1 | YSP{rDNA::GAL10p-synTCPR-TDH3t/GAL1p-synT5OH_M1-GPM1t-HIS3} |  |
| YFT5OH | YTCPR {rDNA:: GAL10p-synTCPR-TPI1t/GAL1p- synT5OH-GPM1t-HIS3} |  |
| YFT5OH_M1 | YTCPR {rDNA:: GAL10p-synTCPR-TPI1t/GAL1p- synT5OH _M1-GPM1t-HIS3} |  |
| CS2-Cas9 | YDBVI {YPRCt3::TEF1p-Cas9-CYC1t-NAT} | This study |
| CS5 | CS2-Cas9 {X4::TEF1p-tHMG1-RBLt-synT5OH-GAL110p-synFOTO-PGK1t-DIT1t-SaGSGPS-Sa-TCCTDHp-GAL7p-IDI} | This study |
| CHD18 | CS5 {1309 ::CPS1t-synBz-Rg-ENO2p-TEF2p-synBz-4C-IDP1t-HIS5t-synCHD-GPMp-TEF1p-sunKAT-ADH1t} | This study |
| YL-V14 | CS8-BzCHD {delta15::GPM1t-synPT6-ENO2-FBA1t-synT1184-M3-GAL110-synT2OH-PGTt} | This study |
| YL_V17 | YL-V14 {W15::TPI1t- synT1184-M3-HSPENO-GAL7- synT5OH-M1-IDP1t} | This study |
| YL_V20 | YL_V17 {720a:: RPL41Bt-SaGGPPS- Sa-TEF2p -PGK1p-tHMG1} | This study |
| YL02 | BB2 {rDNA::KanMX-GAL110p-Sa-SaSGPS-RBLt} | This study |
| YL03 | YL02{deltaDNA::BLE-DIT1t-SaSGPS-Sa-HSPENO} | This study |
| YL04 | YL03 {YPRCt3::TEF1p-Cas9-CYC1t-NAT} | This study |
| YL3P | YL04 {X4::TEF2p-FOTO1-TPI1t-DIT1t-synPCPR1-GAL110p-synT5OH-RBLt} | This study |
| YL3P-M1 | YL04 {X4::TEF2p-FOTO1-TPI1t-DIT1t-synPCPR1-GAL110p-synT5OH-M1-RBLt} | This study |
| BL21(DE3) | *E. coli* expression chassis | Invitrogen |
| Y28A | BL21 {pET28a} | Ref.^2^ |
| YAT5 | BL21 {pET28a-AT5} | Ref.^2^ |
| CXE18 | BL21 {pET28a-CXE18} | This study |
| CXE16 | BL21 {pET28a-CXE16} | This study |
| T9ox | BL21 {pET28a-T9ox} | This study |
| TOGD5 | BL21 {pET28a- TOGD5} | This study |
| EBAIII | BL21{pETDuet-CXE16-T9ox, pRSFDuet-CXE16-CXE18} | This study |
| EBK | BL21{pETDuet, pRSFDuet} | This study |

**Table S2. NMR spectra data of** **9-dihydro-baccatin III (18) and 2D-NMR (COSY, NOESY and HMBC) correlation.**

| Number | ^1^H NMR (δ ppm) | ^13^C NMR  (δ ppm) | Key HMBC |
| --- | --- | --- | --- |
| 1 |  | 78.76 | 2*β*-H, 3*α*-H, 14-H, 16-H, 17-H |
| 2 | 5.74 (d, *J* = 5.9 Hz) | 73.52 | 3*α*-H, 14-H, |
| 3 | 3.11 (d, *J* = 5.9 Hz) | 46.93 | 2*β*-H, 19-H |
| 4 |  | 82.37 | 3*α*-H, 5*α*-H, 6*β*-H, 20*α*-H |
| 5 | 4.93 (d, *J* = 9.1 Hz) | 84.35 | 6*β*-H, 20*β*-H |
| 6 | 6*α*-H: 2.55(m)  6*β*-H: 1.92(m) | 38.07 |  |
| 7 | 4.44(overlapped) | 73.96 | 6-H, 19-H |
| 8 |  | 45.27 | 2*β*-H, 3*α*-H, 19-H |
| 9 | 4.45(overlapped) | 77.16 | 10*α*-H, 19-H |
| 10 | 6.14 (d, *J* = 10.8 Hz) | 74.04 | 9*β*-H |
| 11 |  | 134.42 | 10*α*-H, 16-H, 17-H |
| 12 |  | 143.04 | 10*α*-H, 14-H |
| 13 | 4.80 (m) | 68.66 | 14-H, 18-H |
| 14 | 14*β* -H: 2.28; 14*α*-H: 2.15 | 38.87 | 2*β*-H |
| 15 |  | 42.82 | 10*α*-H, 14-H, 16-H, 17-H |
| 16 | 1.11(s) | 28.47 | 14*α*-H, 17-H |
| 17 | 1.64(s) | 22.22 | 16-H |
| 18 | 2.12(s) | 15.37 |  |
| 19 | 1.82(s) | 12.65 | 3*α*-H, 7*α*-H |
| 20 | 20*α*-H: 4.32 (d, *J* = 8.2 Hz); 20*β*-H: 4.17 (d, *J* = 8.2 Hz) | 76.91 | 3*α*-H |
| 4-OCOCH_3_ | 2.26(s) | 171.77  23.22 | 4-OCOCH_3_ |
| 10-OCOCH_3_ | 2.15(s) | 170.44  21.52 | 10*α*-H, 10-OCOCH_3_ |
| 2-OCOPh | 7.62(m)  8.40(m)  7.49(m) | 167.19  133.83  130.25 129.43 128.79 | 2*β*-H |

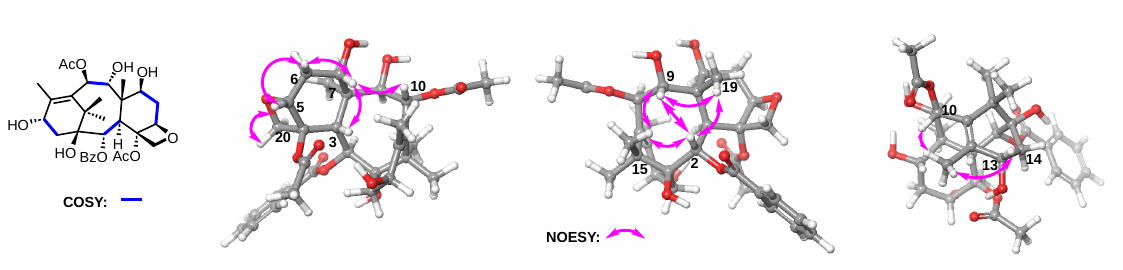

**Table S3: NMR spectra data of baccatin III (19) and 2D-NMR (COSY, NOESY and HMBC) correlation.**

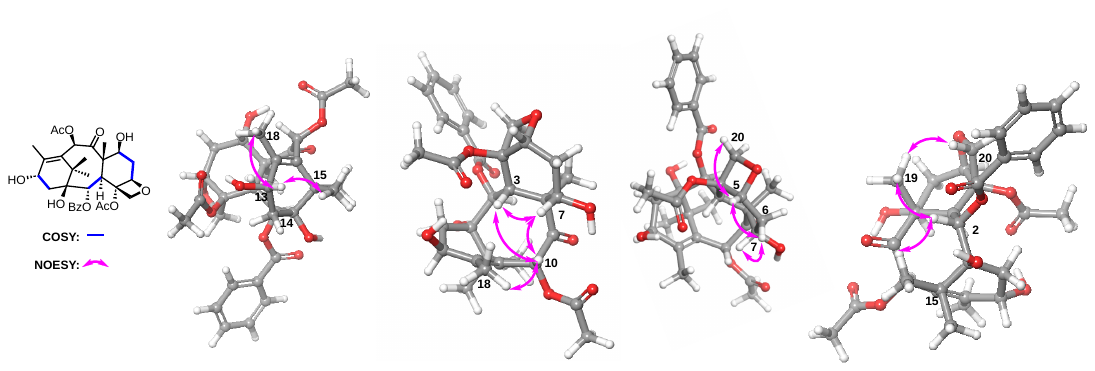

| Number | ^1^H NMR (δ ppm) | ^13^C NMR (δ ppm) | Key HMBC |
| --- | --- | --- | --- |
| 1 |  | 79.24 | 2*β*-H, 3*α*-H, 14-H, 16-H, 17-H |
| 2 | 5.62 (d, *J* = 7.0 Hz) | 75.06 | 3*α*-H, 14-H, |
| 3 | 3.88 (d, *J* = 7.0 Hz) | 46.26 | 2*β*-H, 7*α*-H, 19-H, 20-H |
| 4 |  | 80.95 | 3*α*-H, 5*α*-H, 6-H, 20*α*-H |
| 5 | 4.98 (d, *J* = 7.5 Hz) | 84.60 | 6-H, 20-H |
| 6 | 6*α*-H: 2.56(m)6*β*-H: 1.87(m) | 35.74 | 7*α*-H |
| 7 | 4.47 (dd, *J* = 10.9, 6.7 Hz) | 72.46 | 5*α*-H, 6-H, 10*α*-H, 19-H |
| 8 |  | 58.87 | 2*β*-H, 3*α*-H, 6-H, 19-H |
| 9 |  | 204.30 | 3*α*-H, 7*α*-H, 10*α*-H, 19-H |
| 10 | 6.32 (s) | 76.37 |  |
| 11 |  | 132.01 | 10*α*-H, 16-H, 17-H, 18-H |
| 12 |  | 146.49 | 10*α*-H, 13*β*-H, 14-H, 18-H |
| 13 | 4.89 (t, *J* = 8.2 Hz) | 68.12 | 14-H, 18-H |
| 14 | 2.3 (m) | 38.72 | 2*β*-H |
| 15 |  | 42.85 | 10*α*-H, 14-H, 16-H, 17-H |
| 16 | 1.11 (s) | 21.05 | 14*α*-H, 17-H |
| 17 | 1.11 (s) | 27.11 | 16-H |
| 18 | 2.05 (s) | 15.74 |  |
| 19 | 1.67 (s) | 9.57 | 3*α*-H, 7*α*-H |
| 20 | 20*β*: 4.31 (d, *J* = 8.4 Hz)  20*α*: 4.16 (dd, *J* = 8.4 Hz) | 76.58 | 3*α*-H |
| 4-OCOCH_3_ | 2.28 (s) | 170.80  22.74 | 4-OCOCH_3_ |
| 10-OCOCH_3_ | 2.24 (s) | 171.48  21.05 | 10*α*-H, 10-OCOCH_3_ |
| 2-OCOPh | 7.62  8.40  7.49 | 167.22  133.83 130.24  129.46 128.78 | 2*β*-H |

Compared the NMR data of synthesized Baccatin III this study with literature reported^3^.

| Number | ^1^H NMR (δ ppm) | | ^13^C NMR (δ ppm) | |
| --- | --- | --- | --- | --- |
|  | Synthesized | Literature reported^3^ | Synthesized | Literature reported^3^ |
| 1 |  |  | 79.24 | 80.7 |
| 2 | 5.62 (d, *J* = 7.0 Hz) | 5.62 (d, *J* = 7.2 Hz) | 75.06 | 74.9 |
| 3 | 3.88 (d, *J* = 7.0 Hz) | 3.87 (d, *J* = 7.2 Hz) | 46.26 | 46.1 |
| 4 |  |  | 80.95 | 79.0 |
| 5 | 4.98 (d, *J* = 7.5 Hz) | 4.98 (dd, *J* = 9.7, 2.3 Hz) | 84.60 | 84.4 |
| 6 | 6*α*-H: 2.56(m)6*β*-H: 1.87(m) | 2.56 (ddd, J =14.5, 9.7, 6.6 Hz); 1.85 (ddd, J = 14.5, 10.9, 2.3 Hz) | 35.74 | 35.5 |
| 7 | 4.47 (dd, *J* = 10.9, 6.7 Hz) | 4.47 (dd, *J* =  10.9, 6.6 Hz) | 72.46 | 72.3 |
| 8 |  |  | 58.87 | 58.6 |
| 9 |  |  | 204.30 | 204.2 |
| 10 | 6.32 (s) | 6.32(s) | 76.37 | 76.2 |
| 11 |  |  | 132.01 | 146.5 |
| 12 |  |  | 146.49 | 131.7 |
| 13 | 4.89 (t, *J* = 8.2 Hz) | 4.91-4.88(m) | 68.12 | 67.9 |
| 14 | 2.3 (m) | 2.38 - 2.10 (m) | 38.72 | 38.7 |
| 15 |  |  | 42.85 | 42.6 |
| 16 | 1.11 (s) | 1.10 (s) | 21.05 | 20.9 |
| 17 | 1.11 (s) | 1.10 (s) | 27.11 | 26.9 |
| 18 | 2.05 (d, *J* = 1.4 Hz) | 2.05 (d, *J* = 1.0 Hz) | 15.74 | 15.6 |
| 19 | 1.67 (s) | 1.67 (s) | 9.57 | 9.4 |
| 20 | 20*β*: 4.31 (d, *J* = 8.4 Hz)  20*α*: 4.16 (dd, *J* = 8.4 Hz) | 4.31 (d, *J* = 8.3 Hz, 4.15 (d, *J* = 8.3 Hz) | 76.58 | 76.4 |
| 4-OCOCH_3_ | 2.28 (s) | 2.28 (s) | 170.80  22.74 | 171.4  22.6 |
| 10-OCOCH_3_ | 2.24 (s) | 2.24 (s) | 171.48  21.05 | 170.6  20.9 |
| 2-OCOPh | 7.62  8.40  7.49 | 7.61  8.10  7.48 | 167.22  133.83 130.24  129.46 128.78 | 167.0  133.7  130.1,  129.3,  128.6, |

**Table S4. NMR spectra data of 7-acetoxytaxusin-1,2-diol (13) and 2D-NMR (COSY, NOE and HMBC) correlation.**

| Number | ^1^H NMR (δ ppm) | ^13^C NMR (δ ppm) | Key HMBC |
| --- | --- | --- | --- |
| 1 |  | 79.63 | 2*β*-H, 3*α*-H, 14-H, 16-H, 17-H, 1*β*-OH |
| 2 | 4.07 (m) | 72.33 | 3*α*-H, 2*α*-OH |
| 3 | 3.21 (d, *J* = 6.9 Hz) | 46.61 | 2*β*-H, 19-H |
| 4 |  | 141.14 | 3*α*-H, 20-H |
| 5 | 5.28 (dd, *J* = 4.8, 2.9 Hz) | 75.65 | 3*α*-H, 20-H |
| 6 | α:1.98 (m), *β*: 1.81 (m) | 34.83 | 7*α*-H |
| 7 | 5.36 (dd, *J* = 11.5, 5.5 Hz) | 70.14 | 3*α*-H, 9*β*-H, 19-H |
| 8 |  | 47.26 | 2*β*-H, 3*α*-H, 7*α*-H, 9*β*-H, 10*α*-H,19-H |
| 9 | 5.80 (d, *J* = 10.1 Hz) | 75.94 | 10*α*-H, 19-H |
| 10 | 6.19 (d, *J* = 10.1 Hz) | 71.31 | 9*β*-H |
| 11 |  | 133.72 | 9*β*-H, 10*α*-H, 16-H, 17-H, 18-H |
| 12 |  | 139.21 | 10*α*-H, 14-H, 18-H |
| 13 | 6.17 (overlapped) | 71.23 | 14-H, 18-H |
| 14 | *β*:2.30 (dd, *J* = 15.0, 9.4 Hz); *α*:1.93 (dd, *J* = 15.0, 7.8 Hz) | 37.37 | 2*β*-H, 13*β*-H, |
| 15 |  | 42.27 | 10*α*-H, 16-H, 17-H, 1*β*-OH |
| 16 | 1.29 (s) | 28.79 | 17-H |
| 17 | 1.65 (s) | 22.79 | 16-H |
| 18 | 2.14 (d, *J* = 1.5 Hz) | 15.73 |  |
| 19 | 1.10 (s) | 14.17 | 3*α*-H, 7*α*-H |
| 20 | 5.52 (s), 5.47 (s) | 120.49 | 3*α*-H |
| OCOCH_3_ | 2.21 (s), 2.09 (s), 2.05 (s), 2.04 (s), 2.01 (s) | 21.83, 21.53, 21.46, 21.14, 20.95,170.42, 170.16, 170.02, 169.44, 169.35 |  |

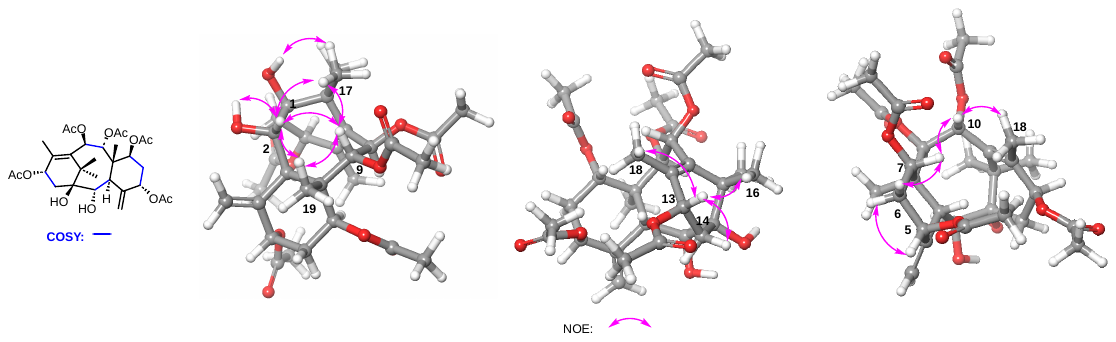

**Table S5. NMR spectra data of 5,10,13-acetyl-10-debenzoyl brevifoliol-2*α*-ol (14) and 2D-NMR (COSY, NOESY and HMBC) correlation**

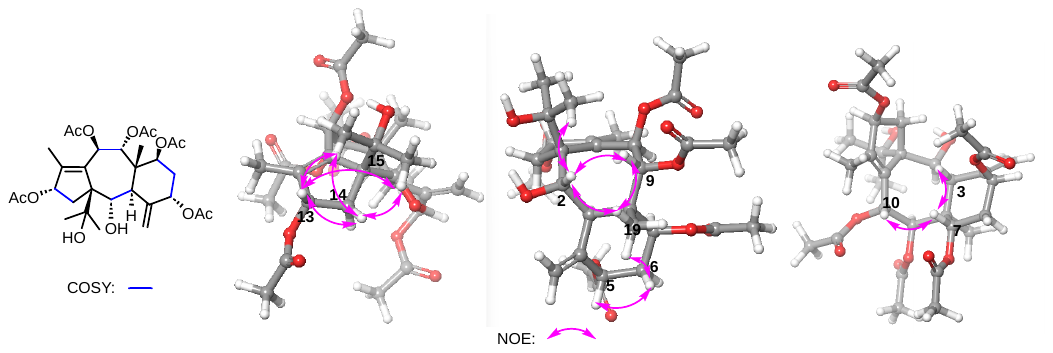

| Number | ^1^H NMR (δ ppm) | ^13^C NMR (δ ppm) | Key HMBC |
| --- | --- | --- | --- |
| 1 |  | 69.28 | 2*β*-H, 3*α*-H, 10-H, 14-H, 16-H, 17-H, |
| 2 | 4.77 (dd, *J* = 9.1, 5.7) | 66.40 | 3*α*-H, 14-H |
| 3 | 2.94 (d, *J* = 9.0 Hz) | 44.28 | 2*β*-H, 5*β*-H, 19-H, 20-H |
| 4 |  | 140.27 | 3*α*-H, 6-H, 20-H |
| 5 | 5.27 (dd, *J* = 4.0, 2.0 Hz | 75.72 | 3*α*-H, 6-H, 20-H |
| 6 | α:1.98 (overlapped),  *β*:1.84 (m) | 34.72 | 7*α*-H |
| 7 | 5.41 (dd, *J* = 11.3, 5.4) | 69.33 | 3*α*-H, 9*β*-H, 19-H |
| 8 |  | 44.98 | 2*β*-H, 3*α*-H, 6-H, 9*β*-H,19-H |
| 9 | 5.75 (d, *J* = 10.7 Hz) | 76.23 | 10*α*-H, 19-H |
| 10 | 6.34 (d, *J* = 10.7 Hz) | 68.58 | 9*β*-H |
| 11 |  | 136.00 | 9*β*-H, 10*α*-H, 13*β*-H, 18-H |
| 12 |  | 147.55 | 10*α*-H, 13*β*-H, 14-H, 18-H |
| 13 | 5.61 (t, *J* = 7.6 Hz) | 78.71 | 14-H, 18-H |
| 14 | *β*:2.37 (dd, *J* = 14.4, 7.3 Hz); *α*:1.66 (overlapped) | 37.21 | 2*β*-H |
| 15 |  | 76.37 | 14-H, 16-H, 17-H, 15-OH |
| 16 | 1.27 (s) | 26.99 | 17-H |
| 17 | 1.37 (s) | 27.46 | 16-H, 15-OH |
| 18 | 1.93 (s) | 11.97 | 13*β*-H |
| 19 | 1.01(s) | 13.73 | 3*α*-H, 7*α*-H, 9*β*-H |
| 20 | 5.46 (s), 5.35 (s) | 118.21 | 3*α*-H |
| OCOCH_3_ | 2.09 (s), 2.07 (s), 2.03 (s), 2.00 (s), 1.96 (s) | 170.67, 169.89, 169.83, 169.67, 168.02, 21.46, 21.36, 21.17, 20.92, 20.86 |  |

**Table S6. NMR spectra data of 2*α*-benzoxy-7*β*-acetoxytaxusin-1*β*-ol (15) and 2D-NMR (HMBC) correlation.**

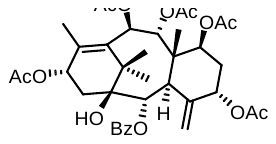

| Number | ^1^H NMR (δ ppm) | ^13^C NMR (δ ppm) | Key HMBC |
| --- | --- | --- | --- |
| 1 |  | 80.01 | 2*β*-H, 3*α*-H, 14-H, 16-H, 17-H, |
| 2 | 5.83 (d, J = 7.0 Hz) | 73.60 | 3*α*-H |
| 3 | 3.49 (d, J = 7.0 Hz) | 46.24 | 2*β*-H, 19-H |
| 4 |  | 139.91 | 3*α*-H, 20-H |
| 5 | 5.27 (dd, J = 4.4, 2.7 Hz) | 75.98 | 20-H |
| 6 | α:1.99 (m), *β*: 1.81 (m) | 35.23 | 7*α*-H |
| 7 | 5.44 (dd, J = 11.5, 5.4 Hz) | 69.93 | 3*α*-H, 9*β*-H, 19-H |
| 8 |  | 47.26 | 2*β*-H, 3*α*-H, 7*α*-H, 9*β*-H, 10*α*-H,19-H |
| 9 | 5.98 (d, J = 10.5 Hz) | 75.76 | 10*α*-H, 19-H |
| 10 | 6.25 (d, J = 10.4 Hz) | 71.30 | 9*β*-H |
| 11 |  | 133.66 | 9*β*-H, 10*α*-H, 16-H, 17-H, 18-H |
| 12 |  | 139.21 | 10*α*-H, 14-H, 18-H |
| 13 | 6.20 (t, *J* = 8.7 Hz) | 71.00 | 14-H, 18-H |
| 14 | *β*:2.38 (m); *α*: 2.20 (m) | 37.25 | 2*β*-H, 13*β*-H, |
| 15 |  | 42.27 | 10*α*-H, 16-H, 17-H, 1*β*-OH |
| 16 | 1.28 (s) | 28.93 | 17-H |
| 17 | 1.77 (s) | 22.82 | 16-H |
| 18 | 2.12 (s) | 15.63 |  |
| 19 | 1.11 (s) | 14.07 | 3*α*-H, 7*α*-H |
| 20 | 5.33 (s), 4.75 (d, J = 1.2 Hz) | 119.48 | 3*α*-H |
| OCOCH_3_ | 2.24 (s), 2.15 (s), 2.06 (s, 6H), 2.03 (s) | 21.92, 21.62, 21.48, 21.13, 20.94, 170.44, 169.98, 169.96, 169.94, 169.35 |  |
| OCOPh | 8.03–7.95 (m, 2H),  7.57 (m, 1H)  7.45 (t, J = 7.8 Hz, 2H), | 167.12,  128.70, 129.62, 130.27, 133.60, |  |

**Table S7. Compared the NMR data of synthesized 9-dihydro-13-acetyl baccatin III (17) this study with literature reported**^3^**.**

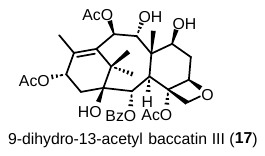

|  | ^1^H NMR (δ ppm) | | ^13^C NMR (δ ppm) | |
| --- | --- | --- | --- | --- |
| Number | Synthesized  (this work) | literature reported^4^ | Synthesized  (this work) | literature reported ^4^ |
| 1 |  |  | 78.92 | 79.51 |
| 2 | 5.76 (d, *J* = 6.0 Hz) | 5.76(d, *J* = 5.95 Hz) | 73.64 | 74.3 |
| 3 | 3.05 (d, *J* = 6.0 Hz) | 3.05(d, J=5.95 Hz) | 47.16 | 49.83 |
| 4 |  |  | 82.16 | 82.85 |
| 5 | 4.96 (d, *J* = 9.0 Hz) | 4.96(dd, *J*=8.72, 1.3 Hz) | 84.17 | 84.83 |
| 6 | 2.55 (m), 1.94 (m) | α: 2.53(ddd, *J* = 14.8, 9.1, 7.5 Hz); β: 1.96, ddd, *J* = 14.8, 9.8, 1.3 Hz) | 38.11 | 38.66 |
| 7 | 4.44 (m) | 4.45 (m) | 74.10 | 74.65 |
| 8 |  |  | 45.10 | 45.50 |
| 9 | 4.45 (overlap) | 4.45 (d, *J* = l0.9 Hz) | 76.70 | 77.51 |
| 10 | 6.18 (d, *J* = 10.6 Hz), | 6.2 (d, *J*= 10.9 Hz) | 73.38 | 73.96 |
| 11 |  |  | 134.93 | 135.74 |
| 12 |  |  | 139.94 | 140.62 |
| 13 | 6.18 (m) | 6.17 (m) | 69.88 | 70.68 |
| 14 | 2.20 (m) | 2.2 (m) | 35.45 | 36.10 |
| 15 |  |  | 43.16 | 43.80 |
| 16 | 1.28 (s) | 1.25 (s) | 23.01 | 23.34 |
| 17 | 1.68 (s) | 1.67 (s) | 28.44 | 29.04 |
| 18 | 1.95 (d, *J* = 1.4 Hz) | 1.92 (d, *J* = 1.4 Hz) | 15.05 | 15.59 |
| 19 | 1.82 (s | 1.89 (s) | 12.63 | 13.25 |
| 20 | 4.31 (d, *J* = 8.3 Hz)  4.16 (d, *J* = 8.3 Hz) | 4.32 (d, *J* = 8.3Hz; 4.17, (d, *J* = 8.3 Hz) | 77.36 | 77.32 |
| 2-OCOPh | 8.09 (m, 2H),  7.61 (m, 1H),  7.48 (m, 2H), | 8.09 (dd, *J*=6.7, 1.4),  7.46 (m),  7.62 (m) | 167.19  133.88, 130.22, 129.34, 128.80 | 167.75,  134.42  130.79,  130.00,  129.37 |
| OCOCH_3_ | 2.28 (s),  2.20 (s)  2.14 (s)  2.01 (s) | 2.3 (s)  2.1 (s)  2.2 (s) | 170.62, 22.73,  170.47, 21.49,  169.49  21.40 | 170.13,  23.57  171.37,  21.96  171.23,  22.07 |

**Table S8. NMR spectra data of 4α,20-epoxide-taxa-11-ene-5*α*-ol (3) and key 2D-NMR (COSY, NOESY and HMBC) correlation.**

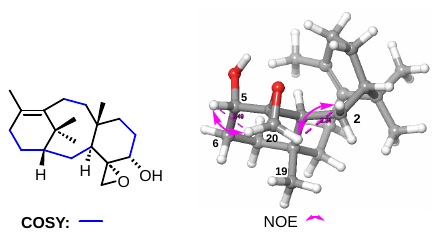

| Number | ^1^H NMR (δ ppm) | ^13^C NMR (δ ppm) | Key HMBC |
| --- | --- | --- | --- |
| 1 | 1.66 (m) | 43.28 | 2-H, 3*α*-H, 16-H, 17-H, |
| 2 | α: 0.80 (ddd, *J* = 14.9, 5.2, 1.8 Hz); β: 1.28 (m), | 23.29 | 3*α*-H |
| 3 | 3.13 (d, *J* = 6.3 Hz) | 31.14 | 2-H, 19-H, 20-H |
| 4 |  | 62.72 | 2-H, 3*α*-H, 5*β*-H, 6-H, 20-H |
| 5 | 3.33 (t, *J* = 3.0 Hz) | 73.38 | 6-H, 7-H, 20-H |
| 6 | 1.83 (m) | 27.93 | 5*β*-H, 7-H |
| 7 | 2.21 (m); 1.03 (m) | 32.18 | 2-H, 3*α*-H, 6-H, 19-H |
| 8 |  | 40.41 | 2-H, 3*α*-H, 6-H, 7-H, 19-H |
| 9 | 2.01 (m); 1.51 (m) | 22.34 |  |
| 10 | 2.31 (m); 1.95 (m) | 30.22 | 9-H |
| 11 |  | 136.60 | 13-H, 18-H |
| 12 |  | 131.42 | 13-H, 18-H |
| 13 | 2.83 (td, *J* = 13.5, 5.3 Hz)  2.10 (m) | 24.78 | 14-H, 18-H |
| 14 | 2.02 (m); 1.19 (ddd, J = 15.4, 5.4, 2.6 Hz) | 40.19 | 2-H, 13-H, |
| 15 |  | 39.38 | 1*β*-H, 9-H, 16-H, 17-H |
| 16 | 1.03 (s) | 31.01 | 17-H |
| 17 | 1.32 (s) | 25.47 | 16-H |
| 18 | 1.83 (s) | 21.39 |  |
| 19 | 0.73 (s) | 22.46 | 3*α*-H, 7-H |
| 20 | 20a: 2.65 (d, *J* = 4.1 Hz)  20b: 2.56 (dd, *J* = 4.1 Hz) | 49.70 | 3*α*-H, 5*β*-H |

**Table S9. NMR spectra data of 7*β*-acetoxytauxin-2*α*-ol (12) and key 2D-NMR (COSY, NOESY and HMBC) correlation**

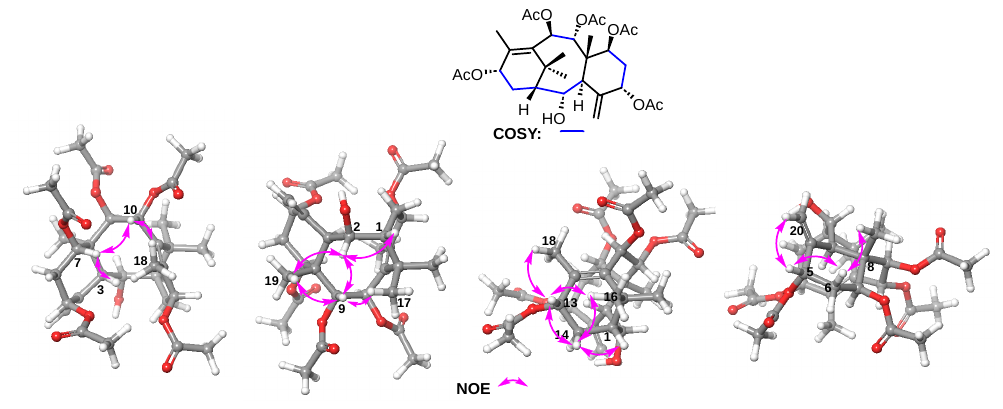

| Number | ^1^H NMR (δ ppm) | ^13^C NMR (δ ppm) | Key HMBC |
| --- | --- | --- | --- |
| 1 | 2.15 (m) | 51.08 | H-3*α*, H-14, H-16, H-17 |
| 2 | 4.28 (m) | 69.34 | H-1*β*, H-3*α* |
| 3 | 3.07 (d, *J* = 6.5 Hz) | 44.90 | H-1*β*, H-19, H-20 |
| 4 |  | 141.45 | H-3*α*, H-6*α*, H-20 |
| 5 | 5.28 (dd, *J* = 4.6, 2.8 Hz) | 75.87 | H-3*α*, H-6, H-20 |
| 6 | 1.99 (m), 1.79(m) | 34.91 | H-7*α* |
| 7 | 5.38 (dd, *J* = 11.4, 5.5 Hz) | 70.32 | H-9*β*, H-6, H-19, C7-OCOCH_3_ |
| 8 |  | 47.26 | H-2*β*, H-3*α*, H-7*α*, H-9*β*, H-10*α*, H-6, H-19 |
| 9 | 5.78 (d, *J* = 10.2 Hz) | 76.22 | H-7*α*, H-10*α*, H-17, H-18, H-19, C9-OCOCH_3_ |
| 10 | 6.16 (d, *J* = 10.2 Hz) | 71.71 | H-9*β*, H-19 |
| 11 |  | 133.21 | H-9*β*, H-10*α*, H-13*β*, H-16, H-17, H-18 |
| 12 |  | 136.96 | H-10*α*, H-13*β*, H-14*β*, H-18 |
| 13 | 6.00 (m) | 70.75 | H-14, H-18, C13-OCOCH_3_ |
| 14 | 2.57 (dt, *J* = 14.8, 9.5 Hz); 1.34 (m) | 27.92 | H-1*β*, H-13*β*, H-16 |
| 15 |  | 37.71 | H-1*β*, H-10*α*, H-14*α*, H-16, H-17 |
| 16 | 1.19 (s) | 32.09 | H-1*β*, H-17 |
| 17 | 1.70 (s) | 27.85 | H-2*β*, H-16 |
| 18 | 2.16 (s) | 15.56 |  |
| 19 | 1.07 (s) | 14.10 | H-3*α*, H-7*α*, H-9*β* |
| 20 | 20a: 5.52 (t, *J* = 1.3 Hz)  20b: 5.46 (d, *J* = 1.4 Hz) | 120.48 | H-3*α*, H-5*β* |
| OCOCH_3_ | 2.22 (s)  2.10 (s)  2.05 (s)  2.03 (s)  2.00 (s) | 170.59, 170.22  170.18, 170.00  169.36, 21.87  21.59, 21.47  21.16, 20.96 |  |

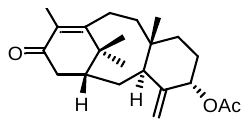
 **5*α*-acetoxytaxadiene-13-one (6)**

^1^H NMR (500M NMR, CDCl_3_) δ ppm: 5.27 (brs, 1H, 5*β*-H), 5.08 (s, 1H, 20-H), 4.73 (s, 1H, 20-H), 3.14 (brs, 1H, 3*α*-H), 3.03-2.94 (m, 1H, 10-H), 2.88 (dd, *J* = 19.2, 7.0 Hz, 1H, 14-H), 2.37 (d, *J* = 13.9 Hz, 1H, 10-H), 2.25-2.14 (m, 2H, 9-H and 1*β*-H), 2.12-2.02 (m, 1H, 7-H), 2.00 (s, 3H, 18-H), 1.98-1.96 (m, 3H, 22-H), 1.92 (d, *J* = 18.7 Hz, 1H, 14-H), 1.80-1.75 (m, 2H, 6-H), 1.75-1.70 (m, 2H, 2-H), 1.45 (s, 3H, 16-H), 1.49-1.40 (m, 1H, 9-H), 1.14 (s, 3H, 17-H), 1.17-1.09 (m, 1H, 7-H), 0.68 (s, 3H, 19-H);

^13^C NMR (125M NMR, CDCl_3_) δ ppm: 200.14(C13), 170.54(C21), 162.57(C11), 150.06(C4), 133.11(C12), 111.71(C20), 75.95(C5), 41.06(C1), 40.56(C15), 40.51(C8), 40.02(C14), 38.44(C9), 37.02(C17), 36.92(C3), 33.40(C7), 28.31(C6), 27.10(C10), 26.79(C2), 24.68(C16), 22.30(C19), 21.63(C22), 13.58(C18).

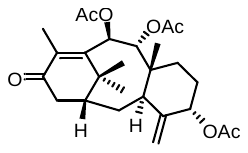
 **Taxusin-13-one**

^1^H NMR (600 MHz, CDCl_3_) δ ppm: 6.09 (d, *J* = 10.4 Hz, 1H, 10*α*-H), 5.86 (d, *J* = 10.6 Hz, 1H, 9*β*-H), 5.31 (t, *J* = 2.6 Hz, 1H, 5*β*-H), 5.21 (s, 1H, 20-H), 4.84 (s, 1H, 20-H), 2.98 (d, *J* = 5.3 Hz, 1H, 3*α*-H), 2.92 (dd, *J* = 19.6, 7.2 Hz, 1H, 14*β*-H), 2.23 (s, 3H, 18-H), 2.18-2.16 (m, 1H, 1*β*-H), 2.08 (s, 3H, 24-H), 2.05 (s, 3H, 22-H), 1.96 (s, 3H, 26-H), 1.94-1.88 (m, 2H, 14*α*-H and 6-H), 1.85-1.68 (m, 5H, 2-H, 6-H and 7-H), 1.64(s, 3H, 16-H), 1.13 (s, 3H, 17-H), 0.76 (s, 3H, 19-H);

^13^C NMR (150MHz, CDCl_3_) δ ppm: 200.32(C13), 170.46(C23), 170.25(C21), 169.91(C25), 152.24(C11), 148.70(C4), 137.84(C12), 113.60(C20), 76.64(C9), 75.72(C5), 73.54(C10), 43.39(C8), 41.02(C1), 39.83(C14), 39.81(C15), 37.62(C3), 37.32(C17), 27.68 (C7), 27.44(C2), 26.40(C6), 25.56(C16), 21.54(C26), 21.07(C24), 20.92(C22), 17.47(C19), 13.85(C18).

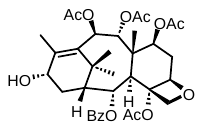
**13-deacetyl-1-dehydroxybaccatin VI (21)**

^1^H NMR (500 MHz, Chloroform-d) δ 8.18 – 8.08 (m, 2H), 7.70 – 7.60 (m, 1H), 7.54 – 7.47 (m, 2H), 6.18 (d, J = 11.2 Hz, 1H, 10*α*-H), 6.01 (d, J = 11.2 Hz, 1H, 9*β*-H), 5.86 (dd, J = 6.0, 2.4 Hz, 1H, 2*β*-H), 5.58 (dd, J = 9.6, 7.8 Hz, 1H, 7*α*-H), 4.98 (d, J = 8.2 Hz, 1H, 5*α*-H), 4.60 (t, J = 8.2 Hz, 1H, 13*β*-H), 4.42 (d, J = 8.4 Hz, 1H, 20-H), 4.17 (dd, J = 8.4, 1.0 Hz, 1H, 20-H), 3.10 (d, J = 5.9 Hz, 1H, 3*α*-H), 2.64 – 2.49 (m, 2H, 14*β*-H and 6*α*-H), 2.30 (s, 3H), 2.24 (d, J = 1.5 Hz, 3H, 18-H), 2.13 (s, 3H), 2.11 (s, 3H), 2.01 (s, 3H), 1.97 (dd, J = 8.7, 2.3 Hz, 1H, 1*β*-H), 1.89 (ddd, J = 15.1, 9.6, 1.5 Hz, 1H, 6*β*-H), 1.84 (s, 3H, 16-H), 1.65 – 1.61 (m, 1H, 14*α*-H), 1.61 (s, 3H, 19-H), 1.03 (s, 3H, 17-H).

^13^C NMR (126 MHz, CDCl3) δ 171.86, 170.32, 170.04, 169.31, 165.07, 142.15(C12), 133.65, 132.91(C11), 129.96, 129.91, 128.76, 84.20 (C5), 81.82(C4), 76.79(C20), 75.69(C9), 72.07(C7), 71.72(C10), 71.56(C2), 67.64(C13), 47.26(C1), 45.99(C8), 44.23(C3), 37.86(C15), 34.78(C6), 31.54(C17), 30.22(C14), 26.76(C16), 23.10, 21.59, 21.17, 20.97, 15.31(C18), 13.02(C19).

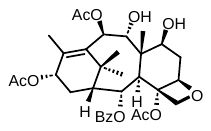
**7,9-deacetyl-1*β*-dehydroxybaccatin VI**

^1^H NMR (400 MHz, Chloroform-*d*) δ 8.13 – 8.04 (m, 2H), 7.63 (t, *J* = 7.4 Hz, 1H), 7.49 (t, *J* = 7.7 Hz, 2H), 6.17 (d, *J* = 10.6 Hz, 1H, 10α-H), 5.96 (t, *J* = 9.0 Hz, 1H, 13β-H), 5.77 (dd, *J* = 6.1, 2.2 Hz, 1H, 2β-H), 5.02 (d, *J* = 9.0 Hz, 1H, 5α-H), 4.48 (m, 2H, 7α-H and 9β-H), 4.39 (d, *J* = 8.3 Hz, 1H, 20-H), 4.18 (d, *J* = 8.3 Hz, 1H, 20-H), 3.39 – 3.29 (m, 1H, 3α-H), 2.91 (d, *J* = 5.9 Hz, 1H), 2.66 – 2.54 (m, 1H), 2.53 – 2.40 (m, 1H), 2.30 (s, 3H), 2.21 (s, 3H), 2.15 (s, 3H), 2.00 (s, 3H), 2.00 – 1.89 (m, 1H), 1.83 (s, 3H), 1.78 (s, 3H), 1.74 – 1.64 (m, 1H), 1.47 (q, *J* = 7.3 Hz, 1H), 1.18 (s, 3H).

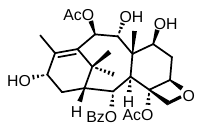
**7,9,13-deacetyl-1*β*-dehydroxybaccatin VI**

^1^H NMR (400 MHz, Chloroform-d) δ 8.11 – 8.06 (m, 2H), 7.63 – 7.58 (m, 1H), 7.48 (t, J = 7.7 Hz, 2H), 6.15 (d, J = 10.7 Hz, 1H, 10α-H), 5.72 (dd, J = 6.0, 2.4 Hz, 1H, 2β-H), 4.97 (dd, J = 8.9, 1.5 Hz, 1H, 5α-H), 4.57 (t, J = 7.8 Hz, 1H, 13β-H), 4.51 – 4.34 (m, 3H, 7α-H, 9β-H and 20-H), 4.19 (d, J = 8.3 Hz, 1H, 20-H), 2.96 (d, J = 5.9 Hz, 1H, 3α-H), 2.60 – 2.48 (m, 2H), 2.27 (s, 3H), 2.13 (s, 3H), 2.13 (s, 3H), 1.99 – 1.86 (m, 2H), 1.80 (s, 3H), 1.71 (s, 3H), 1.63 (dd, J = 15.6, 6.1 Hz, 1H), 1.02 (s, 3H).

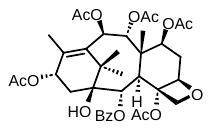
**baccatin VI (16)**

1H NMR (500 MHz, Chloroform-d) δ 8.10 (dt, *J* = 8.4, 1.2 Hz, 2H), 7.61 (td, *J* = 7.5, 1.4 Hz, 1H), 7.48 (td, *J* = 7.8, 1.3 Hz, 2H), 6.21 (dd, *J* = 11.3, 1.2 Hz, 1H, 10*α*-H), 6.17 (t, *J* = 8.8 Hz, 1H, 13*β*-H), 6.00 (d, *J* = 11.3 Hz, 1H, 9*β*-H), 5.87 (d, J = 6.0 Hz, 1H, 2*β*-H), 5.55 (t, J = 8.7 Hz, 1H, 7*α*-H), 4.97 (d, *J* = 9.0 Hz, 1H, 5*α*-H), 4.33 (d, *J* = 8.4 Hz, 1H, 20-H), 4.13 (d, *J* = 8.3 Hz, 1H, 20-H), 3.18 (d, *J* = 6.0 Hz, 1H, 3*α*-H), 2.50 (dt, J = 15.7, 8.5 Hz, 1H), 2.29 (s, 3H), 2.23 – 2.21 (m, 1H), 2.20 (s, 3H), 2.11 (s, 6H), 2.07 – 2.02 (m, 1H), 2.03 (s, 3H), 2.00 (s, 3H), 1.92 – 1.85 (m, 1H), 1.79 (s, 3H), 1.60 (s, 3H), 1.29 – 1.19 (m, 1H), 1.23 (s, 3H).

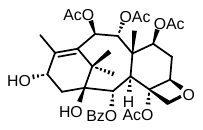
**13-deacetoxybaccatin VI (22)**

1H NMR (500 MHz, Chloroform-d) δ 8.16 – 8.09 (m, 2H), 7.64 – 7.58 (m, 1H), 7.48 (t, *J* = 7.8 Hz, 2H), 6.19 (d, *J* = 11.3 Hz, 1H, 10*α*-H), 5.99 (d, *J* = 11.2 Hz, 1H, 9*β*-H), 5.89 – 5.82 (m, 1H, 2*β*-H), 5.55 (dd, *J* = 9.7, 7.8 Hz, 1H, 7*α*-H), 4.94 (d, *J* = 9.4 Hz, 1H, 5*α*-H), 4.84-4.76 (m, 1H, 13*β*-H), 4.34 (d, *J* = 8.3 Hz, 1H, 20-H), 4.14 (dd, *J* = 8.4, 1.1 Hz, 1H, 20-H), 3.30 – 3.19 (m, 1H, 3*α*-H), 2.56 – 2.45 (m, 1H), 2.35 – 2.28 (m, 1H), 2.27 (s, 3H), 2.20 (s, 3H), 2.18 – 2.12 (m, 1H), 2.11 (s, 3H), 2.10 (s, 3H), 2.00 (s, 3H), 1.86 (ddd, J = 15.1, 9.7, 1.6 Hz, 1H), 1.74 (s, 3H), 1.60 (s, 3H), 1.09 (s, 3H).
